# Genomes reveal age and demographic consequence of ultrafast adaptive radiation

**DOI:** 10.1101/2025.03.07.638630

**Authors:** David A. Marques, Joana I. Meier, Matthew D. McGee, Marcel P. Haesler, Salome Mwaiko, Mary A. Kishe, Anthony Taabu-Munyaho, Lauren J. Chapman, Colin A. Chapman, Sylvester B. Wandera, Catherine E. Wagner, Laurent Excoffier, Ole Seehausen

## Abstract

New species typically evolve over several million years^1^. However, rates of speciation and ecological diversification vary by orders of magnitude across the tree of life^1–3^, with the fastest shown by some adaptive radiations. Eight hundred endemic species of cichlid fishes emerged and formed entire food webs in Lake Victoria and nearby lakes in East Africa. According to Victoria’s paleolimnological history^4^, five hundred may have arisen within the past 16,700 years, but molecular phylogenies estimated a much older origin^5–8^. We reconstruct the age and demography of all Lake Victoria region radiations from whole genomes. We show that indeed, in Lake Victoria all trophic guilds diverged <16,700 years ago, corresponding to between 537 and nearly 30000 speciation events per species per million years, the fastest speciation rate in metazoans^1,2^. Cichlid radiations in lakes Edward, Albert and Kivu too began <20,000 years ago, an order of magnitude faster than previously thought. Evolutionary transitions between trophic levels led to divergence in effective population sizes as predicted by the trophic pyramid of numbers concept^9^ and replicated across three parallel food web radiations. Our results demonstrate that classical theory of trophic interactions in ecologically assembled food webs applies equally to food webs that assembled through rapid adaptive radiation.

## Main Text

Adaptive radiations, when single lineages evolve into ecologically diversified species flocks^10^, are often associated with a burst of rapid successive speciation events^11^. However, since adaptive radiations are mostly studied from a macroevolutionary perspective long after they had unfolded, with extinction rates likely varying too, it remains unclear how punctuated in time and rapid speciation bursts in adaptive radiations really are^12^ and at what rate ecological diversification follows^10,13–15^. Furthermore, even though the ecology of adaptive radiation is rather well understood^10^, the demographic consequences of rapid transitions between trophic levels associated with speciation in adaptive radiations, have never been studied. Recent large adaptive radiations with many young sympatric species at multiple trophic levels can help to close these knowledge gaps.

Here, we reconstruct the age and demographic history of multiple adaptive radiations of haplochromine cichlid fishes in the Lake Victoria region, East Africa, together referred to as the “Lake Victoria region superflock” (LVRS)^16,17^ (Fig. 1c). The largest and most stunning radiation consists of at least 500 species that have evolved in and are endemic to Lake Victoria^16–18^. Others include 80 species in Lake Edward, 15 in Lake Kivu, ten in Lake Albert, and two species in Lake Saka^16–20^. In each lake with ten or more species, cichlids form multi-level trophic networks with guilds feeding on plants, planktonic algae, firmly attached filamentous algae, diatoms attached loosely to filamentous algae, detritus (primary consumers), insect larvae, zooplankton, shrimps, mollusks (secondary consumers), as well as guilds specialized on biting off scales or stealing eggs and juveniles from other cichlids, or preying on other fish (tertiary consumers)^16–18^.

**Fig. 1.**
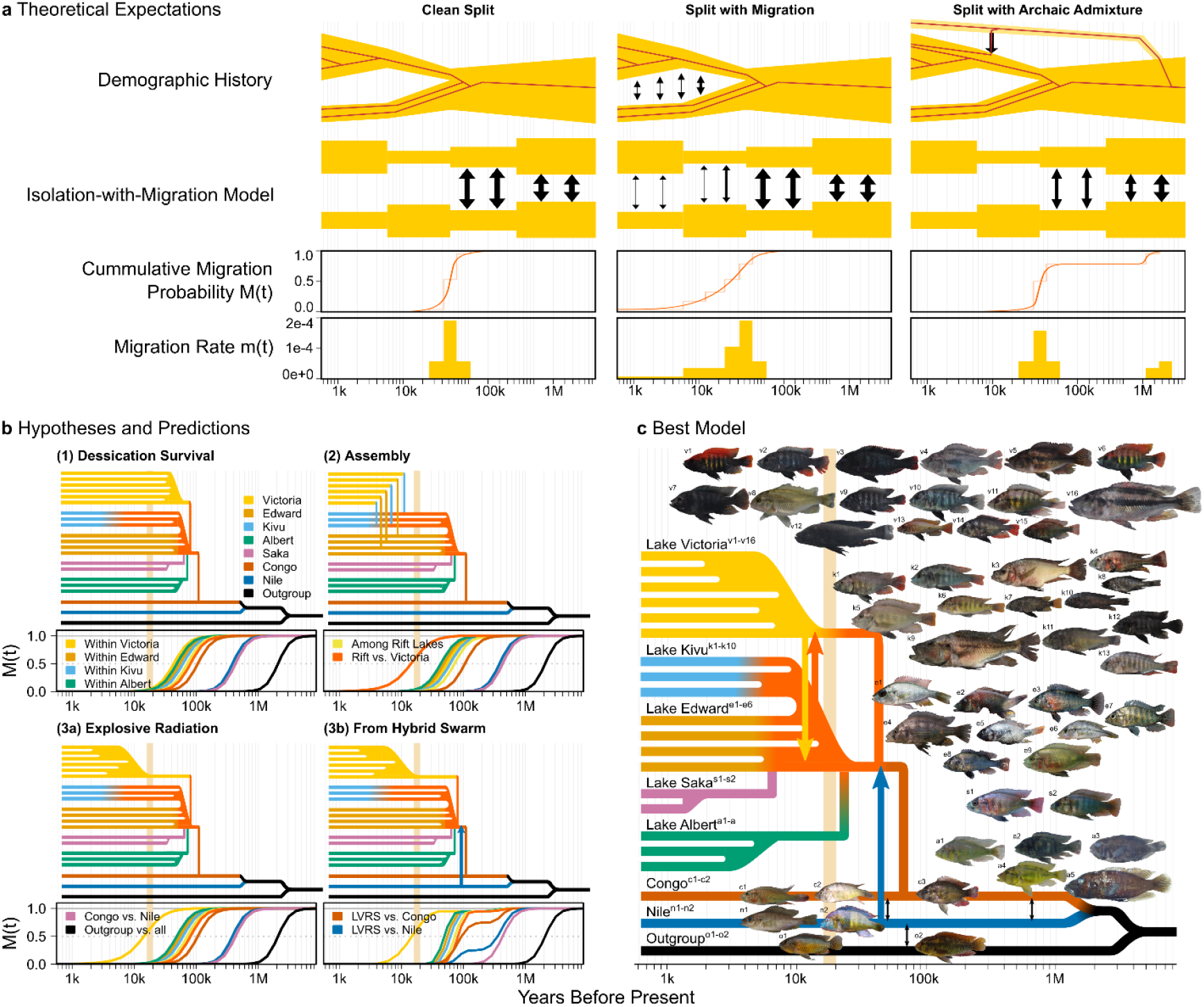
Evolutionary history hypotheses and expected coalescence patterns among Lake Victoria region cichlids. **a**, Three speciation scenarios and expected coalescence patterns inferred with MSMC-IM: if a species splits instantaneously into two isolated species (‘clean split’), the split time coincides with a 50% probability that the lineages have merged backwards in time – a cumulative migration probability M(t) of 50%^42,43^ – and a peak in migration rate m(t). Gene flow after speciation (‘split with migration’) extends the migration episode in m(t) into more recent time and lets M(t) exceed 50%^43^ at the split time, but both still peak at the split time. Gene flow from a third species (‘split with archaic admixture’) delays the complete merging of lineages backwards in time – an M(t) value of 100% – to the split time from the third species, leads to multiple migration m(t) episodes and to an M(t) plateau proportional to the admixture contribution^43^. **b**, Four hypotheses for the evolutionary history of Lake Victoria region cichlid radiations and expected lineage merging patterns within and between LVRS radiations, sister lineages and outgroups for M(t). **c**, Summary of the best supported model of explosive radiations from a hybrid swarm in Lake Victoria, the shared Kivu / Edward radiation, in Lake Albert and Lake Saka, supported by genetic exchange between Western rift lake and Victoria radiations at their base. Species included in this study: Lake Victoria, v1: *Pundamilia nyererei*, v2: *Pundamilia pundamilia*, v3: *Mbipia mbipi*, v4: *Labrochromis* sp. “stone”, v5: *Paralabidochromis chilotes*, v6: *Pundamilia* sp. “nyererei-like”, v7: *Neochromis greenwoodi*, v8: *Macropleurodus bicolor*, v9: *Pundamilia* sp. “pundamilia-like”, v10: *Neochromis omnicaeruleus*, v11: *Astatotilapia* sp. “nubila swamp red”, v12: *Lipochromis* sp. “velvet black cryptodon”, v13: *Yssichromis pyrrhocephalus*, v14: *Lithochromis* sp. “yellow chin”, v15: *Enterochromis cinctus* “E”, v16: *Harpagochromis vonlinnei;* Lake Albert, a1: *Astatotilapia* sp. “aeneocolor-like 1”, a2: *Thoracochromis wingatii*, a3: *Neochromis* sp. Albert, a4: *Thoracochromis mahagiensis*, a5: *Thoracochromis avium*; Lake Saka, s1: Incertae sedis (I.s.) sp. “yellow-red chest” Saka, s2: I. s. sp. “blue” Saka; Lake Edward, e1: *Astatotilapia oregosoma*, e2: *Psammochromis schubotzi*, e3: *Ptyochromis* sp. “Rotbrust” Edward, e4: *Gaurochromis* sp. “huge” Edward, e5: *Enterochromis nigripinnis*, e6: *Astatotilapia schubotziellus*, e7: I. s. sp. “steilstirn” Edward, e8: *Haplochromis* sp. “cf. serridens”, e9: *Haplochromis limax*; Lake Kivu, k1: I. s. *scheffersi*, k2: I. s. *crebridens*, k3: I. s. *occultidens*, k4: I. s. *graueri*, k5: I. s. sp. “littoral light Paralabidochromis-like sp2”, k6: I. s. *paucidens*, k7: I. s. sp. “demersal sp1”, k8: I. s. *kamiranzovu*, k9: I. s. *vittatus*, k10: I. s. *olivaceus*, k11: I. s. sp. “littoral black Mbipia-like “, k12: I. s. sp. “littoral black Pundamilia-like “, k13: I. s. sp. “littoral light Haplochromis-like “; Congo, c1: *Astatotilapia* sp. “cf. katavi” Rukwa, c2: *Astatotilapia* sp. “Yaekama”, c3: *Astatotilapia stappersi*; Nile, n1: *Thoracochromis gracilior*, n2: *Thoracochromis pharyngalis*; outgroups, o1: *Pseudocrenilabrus multicolor*, o2: *Astatoreochromis alluaudi*.

### Lake Victoria region superflock debate

When and how each LVRS radiation evolved has been heavily debated^5–8,21–27^. Paleolimnology of the lakes and geology of the basins revealed a complex history of lake origination, desiccation, and refilling, and basin capture between the Nile and the Congo^28^. Lake Victoria originated only 500,000 years ago in a depression between the West and East African rifts, but has completely dried out repeatedly, including during the most recent arid period from 20,200 to 16,700 years ago^4,29–32^. The Western rift lakes Albert, Edward, and Kivu were part of a single large paleo-lake Obweruka from 7.5 to 2.5 million years (MY) ago and fractured into separate basins with the Ruwenzori and Virunga mountains uplifts 2.5 MY^33^ and 12,000 years ago^34^. All three Western rift lakes presumably persisted through the recent arid period, with shallower lakes Edward and Albert reduced to small, possibly alkaline, lakes^35,36^.

All LVRS cichlid radiations together form a monophyletic lineage^6,7,26,37^ that descends from a hybrid swarm that had formed between divergent and previously geographically isolated lineages from the Upper Nile and the upper Congo^26^. The most recent common ancestor of all LVRS cichlids was estimated to have lived between 100,000-300,000 years ago^5,8,26^, but depending on molecular clock assumptions, estimates ranged from 10,000 years^6^ to 4.7 MY^25^ (Supplementary Note 1). The Lake Victoria cichlid radiation itself forms a monophyletic clade that descends from a hybrid swarm between other refugial LVRS cichlid lineages^37^. The most recent common ancestor of all Lake Victoria cichlids was estimated to have lived 89,000-132,000 years ago^5,8^, but these times are in conflict with the post-desiccation lake age of <16,700 years^4,29,30,32^ and have led to an enduring debate about the origin of the lake’s endemic cichlids and its trophic diversity^4,5,22–24^.

While non-linearity of molecular clocks in the past 1-2 million generations^38^, uncertainty in calibration points^27^ and hybridization^26,37^ might explain why previous molecular clock age estimates are older than the lake, three hypotheses on the origin of Victoria’s cichlid radiation have been put forward (Fig. 1b). (1) Desiccation survival: the Lake Victoria radiation unfolded before the recent arid period and contemporary species from several trophic guilds survived desiccation^5^. (2) Colonization assembly: the Lake Victoria radiation consists of species and trophic guilds that evolved earlier either in Lakes Edward, Albert or Kivu and invaded Lake Victoria post-desiccation^5,6^. (3) Explosive radiation: a single lineage rapidly radiated in the modern Lake Victoria, facilitated either by (3a) a very large standing genetic variation^21,39^ or (3b) the renewed hybrid swarm origin of the Lake Victoria radiation^26,37,40,41^. Testing these hypotheses is of fundamental importance to understanding how fast bursts of speciation and ecological diversification can occur.

### Whole genome demographic reconstruction

Reconstructing the evolutionary history of rapid adaptive radiations from DNA sequence data is challenging because of incomplete lineage sorting and hybridization leading to conflicting evolutionary histories across the genome^44^, complicating age estimation with methods that assume bifurcating trees and lineage sorting. We take advantage of whole genomes to estimate local evolutionary histories along chromosomes with the multiple sequentially Markovian coalescent algorithm^42^ (MSMC2). MSMC2 estimates pairwise coalescence times between all haplotypes in a sample along chromosomes and from this infers coalescence rates for different time periods in the past, both within and between species. We fit isolation-with-migration (IM) models to these coalescence rates^43^ (MSMC-IM) for each pair of species to infer episodes of gene flow and historical effective population sizes (N_e_)^42^.

Thinking backwards in time, during speciation two lineages merge back into one lineage and the between-species coalescence rate increases from zero to 100% of the within-species rate^42^ (Fig. 1a). MSMC-IM approximates this ratio with the cumulative migration probability M(t)^43^, the probability of lineages to have merged at time t. Expectations for M(t) and episodes of migration m(t) depend on whether following speciation, two species continue to exchange genes, or if archaic admixture from hybridization with additional species has occurred^42,43^ (Fig. 1a), both likely scenarios for Lake Victoria region cichlids^26,37,45^. Consequently, for the three hypotheses on the origin of the Lake Victoria radiation, we expect different patterns and timings of lineage merging within and between LVRS radiation members, sister lineages and outgroups (Fig. 1b).

We sequenced 112 whole genomes of 56 cichlid species, sampled such as to maximize the eco-morphological diversity within each lake radiation of the LVRS, and included their sister lineages and outgroups (Fig. 1c, Supplementary Table 1). To parametrize the MSMC2 and MSMC-IM reconstruction, we estimated a single base pair mutation rate of 2.4 × 10^−9^ mutations per generation and base pair for the Victoria cichlid *Gaurochromis hiatus* by sequencing 11 genomes of a three-generation family (Supplementary Note 2). The MSMC-IM method resolved changes in historical N_e_ and estimated gene flow between lineages in a time frame between ∼700-12,000 years and ∼6 MY ago, depending on the number of haplotypes included per species^42^, ideally suited to resolve the evolutionary history of the LVRS radiations.

**Table 1.** Speciation rate estimates. Speciation rate (λ) and speciation interval (SI) estimates for LVRS radiations, assuming a maximally balanced (log) or maximally unbalanced (lin) tree, see Methods.

| LVRS radiation | No. of species | Age [y] | Speciation rate [no. of spec. events per MY] |  | Speciation interval [years] |  |
| --- | --- | --- | --- | --- | --- | --- |
| | | | $\lambda_{\log}$ | $\lambda_{\text{lin}}$ | SI<sub>log</sub> | SI<sub>lin</sub> |
| Victoria | 500 | <16,700 | 537 | 29,880 | 1,863 | 33 |
| Edward / Kivu | 95 | <20,000 | 328 | 4,700 | 3,044 | 213 |
| Albert | 10 | <10,000 | 332 | 900 | 3,010 | 1,111 |
| Saka | 2 | <2,000 | 500 | 500 | 2,000 | 2,000 |

### Young age of all LVRS radiations

We found that all Lake Victoria radiation guilds rapidly merge (backwards in time) into an ancestral lineage during the refilling of the lake following the end of the arid period ∼16,700 years ago (Fig. 2c), with an 59-82% probability of lineages having merged 16,000 years ago (Fig. 2a) and migration peaking during or at the end of the arid period (Fig. 2d, Supplementary Fig. 1). This finding suggests very recent speciation and evolution of trophic guilds in modern Lake Victoria – a radiation age of around 16,700 years. Importantly, coalescence between members of different guilds is consistent with an early but not an immediate origin of all trophic guilds upon refilling of the lake: predators, paedophages, snail crushers and specialized insect pickers likely merged most anciently and thus diverged first in the radiation, followed by rocky shore specialists and most recently open water cichlids (Supplementary Note 3). A second migration peak in the most recent 2,000 years may be explained by hybridization and further speciation during major environmental change in the offshore habitat of Lake Victoria around that time^45,48^ associated with increased cichlid abundances in that habitat^46^. A third migration peak 30,000-80,000 years ago is consistent with archaic admixture during the formation of the LVRS hybrid swarm lineage^26^ (Supplementary Note 4).

**Fig. 2.**
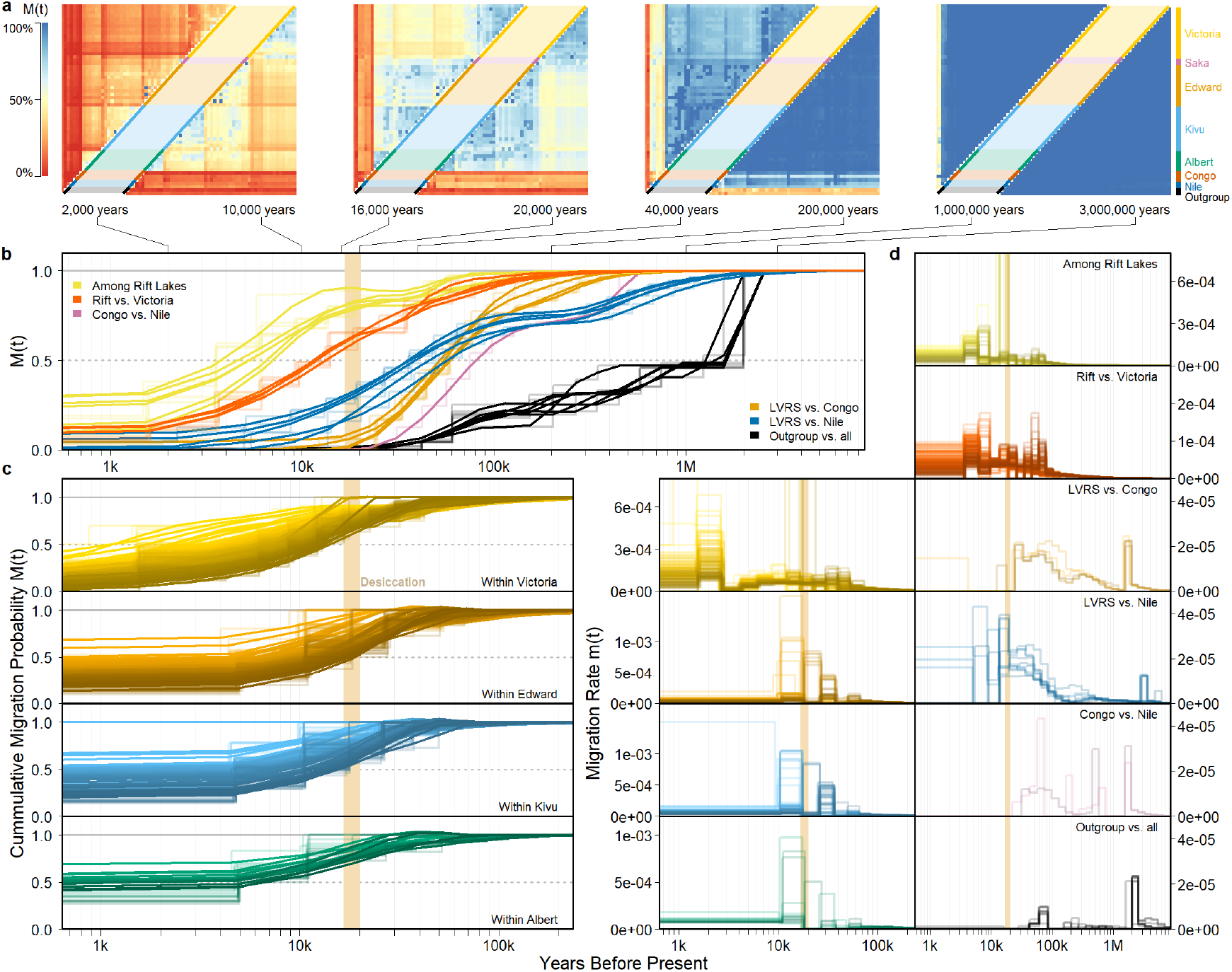
Observed lineage merging among Lake Victoria region cichlids, sister lineages and outgroups. Cumulative migration probability M(t) (**a-c**) and migration rate m(t) (**d**) illustrate the timing of lineages merging in a common ancestor for species of haplochromine cichlids from the Lake Victoria region superflock radiations, Nile and Congo sister lineages and one outgroup (*Astaoreochromis alluaudi*). **a**, M(t) heat maps for different time points illustrate that same-lake species merge more recently into an ancestral lineage backwards in time than species from different lakes, especially between Lake Victoria and Western rift lakes. **b**, Coalescence of LVRS radiation cichlids, sister lineages and the outgroup reconstructed from four haplotypes per species, of one representative species each. Western rift lake cichlids lineages merge with each other more recently than they merge with Lake Victoria cichlids in a common ancestor, consistent with parallel radiations in Lake Victoria and Western rift lakes. Complete lineage merging of all LVRS cichlids only 50,000-100,000 years ago and a plateau between the major and complete lineage merging with Nile cichlids support the hybrid origin of all LVRS cichlid lineages. **c**, Coalescence within each LVRS radiation reconstructed from six (Victoria) and two haplotypes (Western Rift lakes) per species. In each lake radiation, species merge in a common ancestor during or more recently than the arid period 16,700-20,200 years ago, consistent with very recent, explosive radiations. **d**, Migration rates within (left) and between (right) LVRS radiations, sister lineages and outgroups reconstructed from six (within Victoria radiation) and two haplotypes (all others) per species. Major migration episodes around the arid period 16,700-20,200 years are consistent with explosive radiations in Victoria and Western rift lakes following the arid period. See Supplementary Figs 1-4 for further details.

Surprisingly, Western rift lake cichlids also merge very recently into an ancestral lineage: already 20,000 years ago, the probability of having merged in a common ancestor is 54-100% for all Lake Edward cichlids, 16,000 years ago the probability is 50-99% for all Lake Kivu cichlids, and even just 10,000 years ago the probability is 66-88% for Lake Albert cichlids (Fig. 2a, c, Supplementary Figs. 2-4). Much like for Lake Victoria cichlids, most of the lineage merging between species within Western rift lakes occurs in a migration episode either after the end of or during the arid period, with a second older migration episode 30,000-80,000 years ago (Fig. 2d, Supplementary Figs. 2-4), coincident with the LVRS hybrid origin^26^ (Supplementary Note 4). In crater lake Saka, complete lineage merging between the two species occurs already in the last just 1,000-2,000 years (Supplementary Fig. 5), consistent with the very young geological age of this crater lake and weak genetic species differentiation^19^ suggesting a more recent speciation event. Radiations are thus of an estimated age of 16,000-20,000 years for Edward and Kivu, 10,000 years for Lake Albert and less than 2,000 years for Lake Saka.

Importantly, LVRS cichlids inhabiting the same lake merge more recently and rapidly into a common ancestor than those inhabiting different lakes (Fig. 2a, 3, Supplementary Figs. 6-26), suggesting largely independent^37^, parallel adaptive radiations within lakes, with two exceptions. First, many Kivu and Edward cichlid species merge faster in a common ancestor between lakes than within lake (Fig. 3f,j), suggesting they represent a single adaptive radiation shared today between the two lakes that were connected until 12,000 years ago, consistent with their polyphyly in phylogenetic trees^37^. Second, a few species, including piscivorous and paedophagous predator species in lakes Victoria and Edward tend to merge faster between lakes than within lake in the most recent time period following the desiccation (Supplementary Figs. 7, 11), consistent with substantial across-lake excess allele sharing between these guild members^37^ due to genetic exchange between the Western Rift lake and Victoria lineages during the unfolding of each radiation. Migration peaks around the arid period 16,700-20,200 years ago between many Lake Victoria and Western rift lake species (Fig. 2d) is further consistent with genetic exchange not only limited to these guilds^37^.

**Fig. 3.**
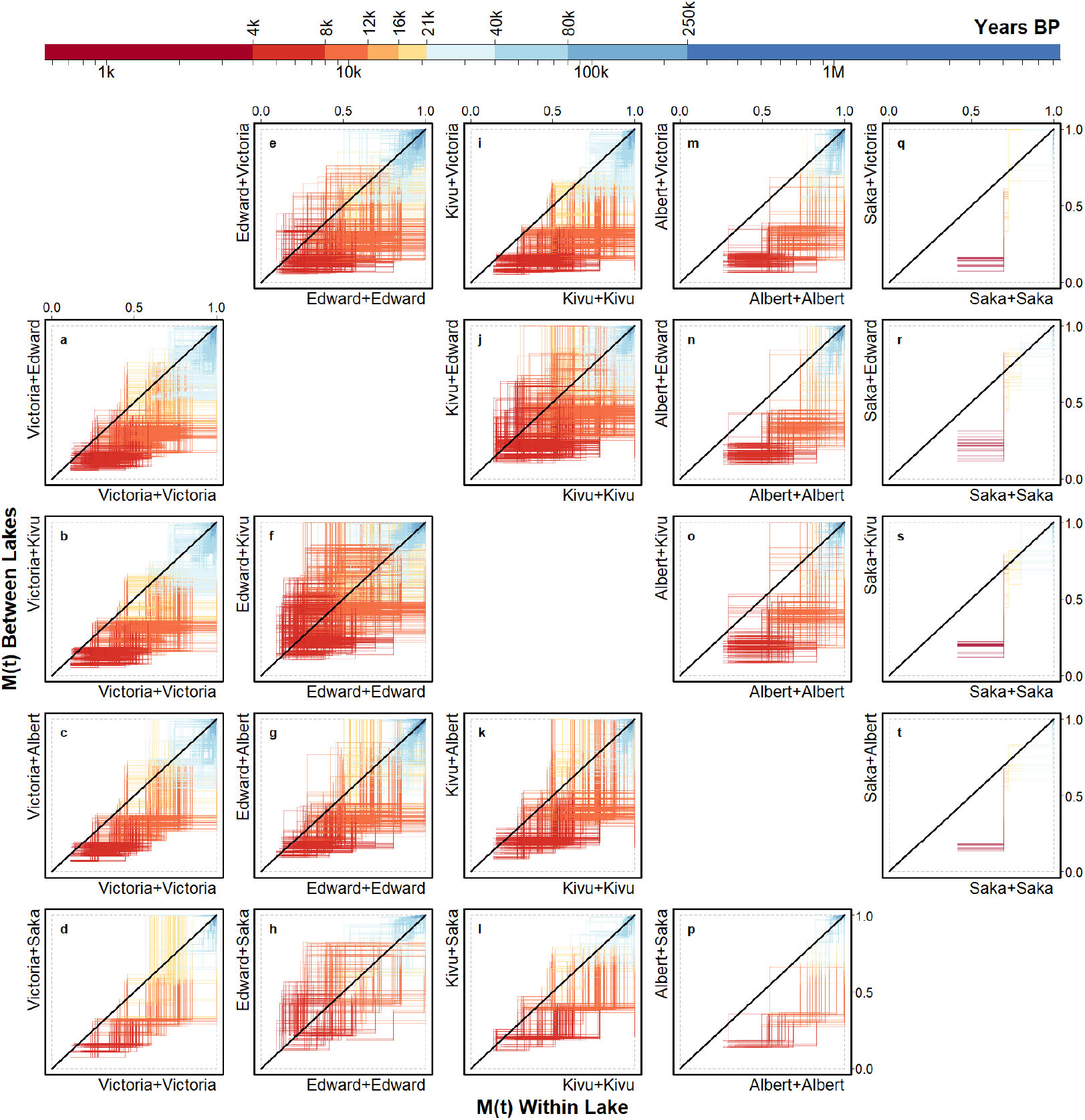
Observed lineage merging among Lake Victoria region cichlids from the same and different lakes. Each trajectory shows the probability of a species having merged in a common ancestor with another species from the same lake (x axis) vs. with another species from a different lake (y axis), during different time intervals indicated by the color scale. Trajectories below the diagonal indicate faster lineage merging between species from the same lake, than with species from another lake – consistent with speciation and thus radiation within a lake, a pattern predominant in lakes Victoria (**a-d**), Albert (**m-p**) and Saka (**q-t**). In contrast, the symmetrically distributed trajectories in comparisons of lake Kivu and Edward species (**f, j**) suggest a single, shared radiation. While Saka species merge more rapidly into a common ancestor within lake (**q-t**), some Edward species do merge more rapidly with these Saka cichlids than with other Edward species (**h**), suggesting that the recent Saka radiation is nested in the Edward radiation. A few trajectories (**a, e**) involving two predator (*Harpagochromis vonlinnei, Prognathochromis* sp. 2 Edward) and two paedophage species (*Lipochromis* sp. “velvet black cryptodon”, *Lipochromis taurinus*) from Victoria and Edward, respectively, and *Astatotilapia* sp. “Rotbrust” Edward pass the diagonal line in recent times, suggesting recent gene exchange between lakes, at the base of both radiations or even later migration from Lake Victoria to Lake Edward (see Supplementary Figs. 7-26 for details). Cumulative migration probabilities M(t) reconstructed from two haplotypes per species are shown for all comparisons.

Hypothesis (1), that today’s Lake Victoria radiation survived the desiccation of Lake Victoria, is inconsistent with the recent merging in a common ancestor of all trophic groups in Lake Victoria. Rather, pre-existing cichlid diversity appears to neither have survived in the Lake Victoria basin, nor in Western rift lakes despite lakes Albert and Edward not drying completely^35,36^. Hypothesis (2) of an assembly of the food web in Lake Victoria through colonization of pre-existing trophically diverse Western rift lake species^5,6^, is inconsistent with faster lineage merging within Lake Victoria than with species of other lakes, and with the monophyly of the Victoria radiation^37^. Hypothesis (3) of an explosive radiation in Lake Victoria is consistent with the recent lineage merging within Lake Victoria and faster coalescence within than between lakes. Given the apparently recent age of Western rift lake cichlid radiations, they too could be considered such explosive radiations. However, hypothesis (3) does not fully explain our results. Instead, we added to hypothesis (3) elements of hypotheses (2) to include genetic exchange with Western rift lakes at the base of the radiation, to build a new best supported model based on the results outlined above (Fig. 1c). After the desiccation and subsequent refilling of Lake Victoria, refugial cichlid populations from headwaters of the Lake Victoria basin^37^ admixed with migrants from the Lake Edward system into a hybrid population that seeded the recent explosive adaptive radiation in Lake Victoria as the new lake refilled and expanded, leading to parallel evolution of trophic networks in Lake Victoria, the Edward / Kivu radiation and in Lake Albert (Supplementary Note 5).

The young age of all LVRS cichlid radiations leads to speciation rates one to two orders of magnitude higher than anything else known in the tree of life^1–3,24^. Five hundred species evolving in the Victoria radiation since the lake refilled 16,700 years ago corresponds to speciation rates between 537 and 29,880 speciation events per species per million years or a speciation interval of between 33 and 1,863 years between successive speciation events (Table 1), depending on whether maximally balanced or completely unbalanced trees are assumed to estimate these per species speciation rates (see Methods). The younger than expected age of Western rift lake cichlid radiations likewise ranks their speciation rates to a similar order of magnitude (Table 1).

### Demographic consequence of adaptive radiation

We discovered a strong relationship between evolution of tropic position and change in effective population size (N_e_) within each LVRS radiation (Fig. 4c-f, Supplementary Note 6). In Lake Victoria, the most recent N_e_ estimates for the 11 species representing major ecological guilds sort them by trophic position and known relative abundances^18^ as predicted by theory (Fig. 4a): we estimate the lowest recent population sizes for the lake-wide distributed paedophage *Lipochromis* sp. “velvet black cryptodon” and predator *Harpagochromis vonlinnei*, as expected by their high trophic position as tertiary consumers and low to very low local census numbers^18^. The highest N_e_ is shown by the pelagic zooplanktivore *Yssichromis pyrrhocephalus* and the demersal detritivore *Enterochromis cinctus*, both also distributed across the lake, but feeding at lower trophic positions and being highly abundant open water species^18^.

**Fig. 4.**
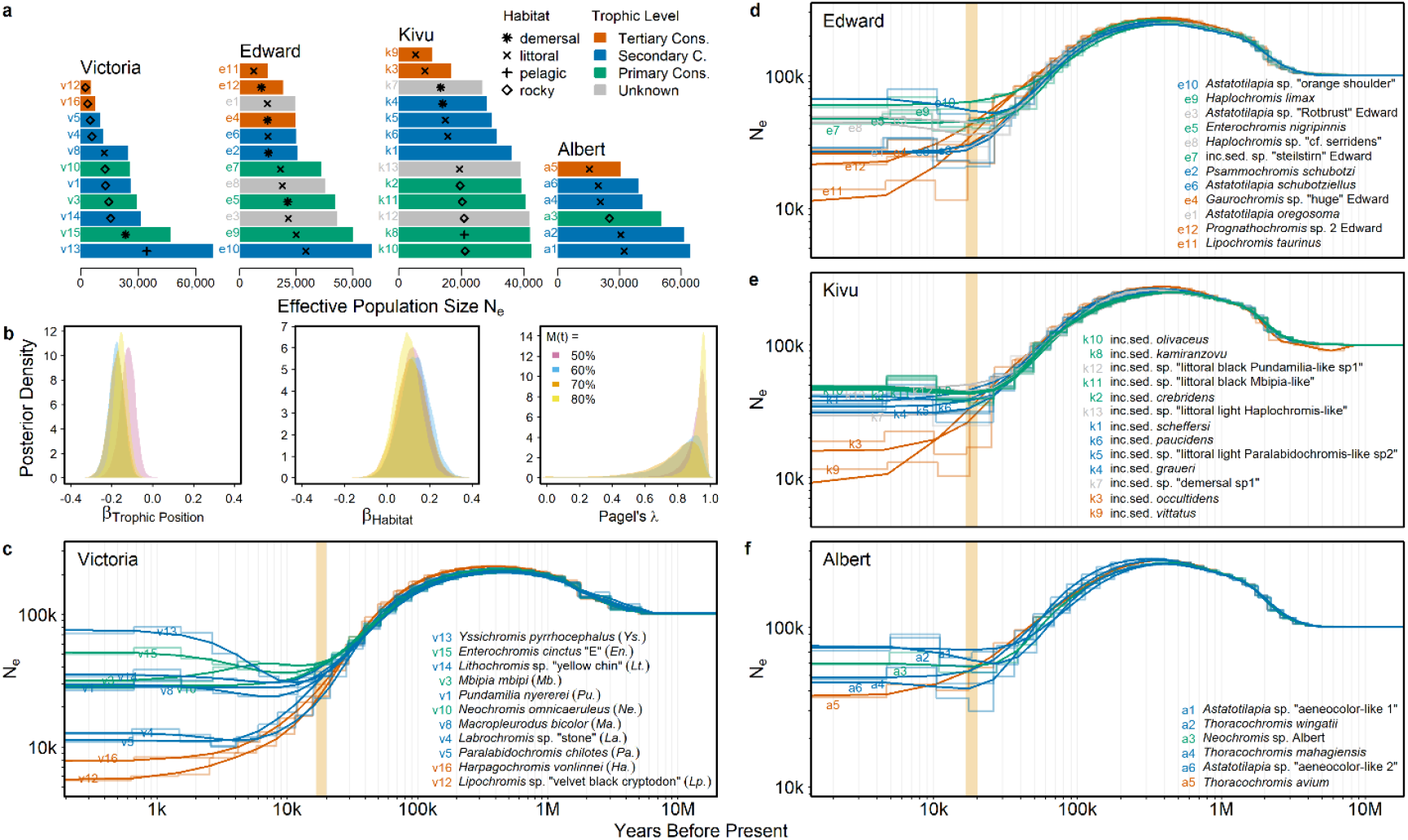
Repeated evolution of trophic pyramids of numbers in Lake Victoria region cichlid adaptive radiations. **a**, Recent effective population size estimates (N_e_) correspond to trophic position and abundance of endemic species in the adaptive radiations of haplochromine cichlids in lakes Victoria, Edward, Kivu and Albert. Predators and paedophages (tertiary consumers) occupying high trophic levels consistently show the lowest effective population sizes in each radiation, while primary consumers such as algae scrapers and super-abundant secondary consumers such as pelagic zooplanktivores show the highest N_e_. The color code indicates trophic levels of each species with symbols and corresponding species names shown in panels **c-f** and species sorted by increasing N_e_. **b**, A Bayesian regression using divergence time estimates from M(t)=50-80% lineage merging shows that height of trophic position has a strong negative effect on N_e_, together with shared evolutionary history (Pagel’s λ) and weak positive effect of open water habitat. **c-f**, The evolution of effective population sizes suggests that trophic niche evolution across all adaptive radiations of cichlids in Lakes Victoria (**b**), Edward (**h**), Kivu (**i**) and Albert (**j**) had immediate demographic consequences with effective population sizes diverging at the same time as the species started to diverge. Estimates from six haplotypes per species are shown in panel **c** and of two haplotypes per species in panels **d-f**.

Intermediate N_e_ estimates are shown by species restricted to patchily distributed littoral habitat, such as the epilithic algae browser *Neochromis omnicaeruleus* and epilithic algae grazer *Mbipi mbipi* limited to rocky shores, reef-restricted zooplanktivores *Lithochromis* sp. “yellow chin” and *Pundamilia nyererei*, and the littoral snail eater *Macropleurodus bicolor*. Among them, two trophically specialized rocky shore species with much lower census sizes^18^ show even lower N_e_: the big-lipped insectivore *Paralabidochromis chilotes* and the pharyngeal mollusk crusher *Labrochromis* sp. “stone”. The same trajectory and hierarchy of N_e_ evolution is found among ecological guild representatives within Lake Edward, Lake Kivu and Lake Albert, where species that have evolved a high trophic position show lower recent N_e_ and species feeding at low trophic positions show larger recent N_e_ estimates (Fig. 4a, d-f).

We validated this observation in a comparative analysis that accounts for non-independence due to shared evolutionary history but relaxes the assumption of tree-like evolution. Instead of a phylogeny-based covariance matrix, we used time estimates at 50% to 80% probability of the species pairs having merged in a common ancestor backwards in time to build a covariance matrix (see Methods). A Bayesian regression of trophic position and habitat (littoral vs. open water) on N_e_ across all LVRS radiations revealed a consistent strong negative effect of trophic position and a weak positive effect of open water habitat on N_e_ (Fig. 4b). At the same time, a Pagel’s λ estimate close to one (Fig. 4b) indicates that shared evolutionary history also strongly predicts variance in N_e_, i.e. that more recently diverged species, such as species from the same lake, tend to show more similar effective population sizes. Food web and habitat niche evolution in these adaptive radiations has thus led to similar, predictable demographic consequences associated with the ecology of the emerging species, and this is replicated across multiple cichlid radiations. The onset of demographic divergence and diversification (Fig. 4) coinciding temporally with the time of divergence between members of different guilds (Supplementary Figs. 1-4) furthermore is in line with predictions from adaptive radiation theory that new ecological roles evolve around the same time that new species arise^10^.

### Reconciling cichlid and lake histories

Our demographic reconstructions shed new light on a longstanding debate about the age and rates of diversification in one of the largest extant metazoan adaptive radiations. We show that the actual speed of both speciation and ecological diversification are one to two orders of magnitude higher than anything else known in the tree of life^1–3,24^. Our approach avoids problematic assumptions for mitochondrial substitution rates or phylogenetic placement of fossils, circularity induced by assuming paleolimnological constraints, or biases due to non-recombining markers, and approaches ignoring non-treelike evolution.

Our reconstruction suggesting divergence between all major guilds^37,40^ in Lake Victoria within the last 16,700 years resolves earlier conflicts between paleolimnological and phylogeographic reconstructions^5–8,21–27^. That the Western rift lake cichlid radiations are nearly as young as the Lake Victoria radiation is surprising and emphasizes both the susceptibility of lacustrine cichlid radiations to environmental change^45,49^ and the remarkable ability of the Victoria region haplochromine lineage to repeatedly form explosive adaptive radiations^37,40^.

### The ecology of evolved trophic networks

We discovered that evolutionary transitions in the ecological niche across trophic levels led to divergence in effective population size consistent with predictions by the trophic pyramid of numbers concept^9^ both in direction and magnitude. Because variation in N_e_ has repercussions on evolutionary rates, such predictable demographic consequences of trophic level evolution may have important implications for the further course of speciation and adaptive radiation^50^. The evolutionary predictability of the classical relationship between ecology and demography is demonstrated by its replicability across three parallel cichlid radiations in lakes Victoria, Albert and Edward / Kivu. Whole genomes and new population genomic methods can thus give unprecedented insight into the intricate feedback between ecology and evolution even in the most challenging systems, explosive adaptive radiations that occurred on very recent timescales, as exemplified by magnitudes faster speciation, ecological diversification and food web evolution with demographic consequences uncovered here for Lake Victoria region cichlid fish.

## Methods

### Taxonomic sampling

To reconstruct the demographic history of LVRS cichlids, we used a phylogenetically, geographically, and ecologically fully nested sampling design: all lakes, multiple ecological guilds in each lake, all major guilds and genera in Lake Victoria. To accomplish this, we selected 56 species and sequenced one to five individuals per species (112 genomes). The various contrasts between these species encompass all major divergence events between lakes and guilds within lakes. From each ecological guild we sampled one species representing a large number of endemic species from the same adaptive radiation, e.g. the paedophage *Lipochromis* sp. “velvet black cryptodon” represents a genus containing ∼15 species specialized on stealing eggs and juveniles from other mouthbrooding cichlids in Lake Victoria, the pelagic zooplanktivore *Yssichromis pyrrhocephalus* and the demersal detritivore *Enterochromis cinctus* each represent a clade of 15 to 20 different but ecologically similar species from Lake Victoria. The contrasts we investigated are (i) between guilds within the Lake Victoria radiation, (ii) between LVRS radiations Victoria, Albert, Edward, Kivu, and Saka, (iii) between LVRS radiations and the founding Congolese and Upper Nile lineages, and (iv) between any of these and a distant outgroup species endemic to the Lake Victoria region *Astatoreochromis alluaudi* or *Pseudocrenilabrus multicolor* (Supplementary Table 1).

Samples were collected with gill nets, hook and line, or minnow traps and euthanized with an overdose of MS-222. This was done in compliance with national and international guidelines for ethical treatment of animals. Field campaigns took place between 1995 and 2019 (Supplementary Table 1) in collaboration with the Tanzania Fisheries Research Institute and the Ugandan National Fisheries Resources Research Institute. Exports were approved by the Ministry of Agriculture, Livestock and Fisheries in Tanzania, the Ministry of Agriculture, Animal Industry and Fisheries in Uganda, and the Department of Fisheries in Zambia.

To estimate the mutation rate in a Lake Victoria cichlid, we used a three-generation family of the Lake Victoria haplochromine cichlid *Gaurochromis hiatus*, in which two F1 full siblings produced F2 offspring. We established this family from fish caught in the Mwanza Gulf of Lake Victoria in 2014 and bred in the fish lab facilities at the University of Bern, Switzerland (Supplementary Fig. 27). The two F0 grandparents, two F1 parents and seven F2 offspring were euthanized with MS-222 according to national ethical regulations under animal experimentation and holding permits BE65/18 and BE1/10 issued by the Cantonal government of Bern, Switzerland.

### DNA extraction, whole genome re-sequencing and alignment

DNA extraction and PCR-free library preparation followed the procedure described in Meier *et al*.^26^. We performed whole genome resequencing of the 112 wild-caught LVRS cichlids, sister lineages and outgroup species and 11 *Gaurochromis hiatus* family members for mutation rate estimation. The majority of individuals were sequenced on an Illumina HiSeq 3500 (Illumina Inc.) to a mean depth of 21.7x (range: 9-49x) and *Gaurochromis* family on an Illumina NovaSeq 6000 (Illumina Inc.) to a mean depth of 50.5x (range: 34-76x, Supplementary Table 1), producing 150bp paired-end reads, at the University of Bern, Switzerland. We aligned reads to an anchored version^51^ of the *Pundamilia nyererei* reference genome^52^ using bowtie2 v2.2.3^53^ in local alignment mode and allowing one mismatch in seed alignment.

### Mutation rate estimation

We called variants and genotypes in the *Gaurochromis* family with three different methods to achieve maximal sensitivity in detecting possible *de novo* mutations. We used the GVCF/HaplotypeCaller pipeline of GATK v3.5^54^, freebayes v1.3.1^55^ and joint calling with HaplotypeCaller of GATK v4.1.2.0^56^. The first method generated base pair resolved GVCF files with HaplotypeCaller, and we specified a contamination fraction of 10%, to consider bases with a minimum quality of 20 and to call and emit variants with quality 30 or higher only. We combined GVCF files into a VCF file containing monomorphic and variant sites of the 11 individuals, using the GATK tool GenotypeGVCF. The second method, freebayes, called jointly variants and genotypes from alignments of the 11 individuals. We set freebayes to consider only bases with quality 20 or higher, positions covered by at least six reads, to down-sample regions covered by more than 500 reads, to skip regions covered by more than 5,500 reads, to output genotype qualities and all monomorphic sites at base pair resolution as GVCF entries and disabled clumping of complex haplotypes. The third method also jointly called variants and genotypes from the 11 alignments using HaplotypeCaller of GATK v4.1.2.0 but did not output monomorphic sites. We specified a contamination fraction of 10%, a minimum base quality of 20, a calling threshold of minimum quality 30 for a variant and a minimum mapping quality per read of one.

To identify candidate mutations, we applied two sets of stringent filters to the calls produced by all three methods following^57–59^. The first set of filters identified the total number of ‘callable sites’ including monomorphic sites, SNPs and indel variants. We removed sites with unusually low or high depth by computing the mean site depth ±1.5 times the interquartile range of site depths across all 11 individuals and removing sites with depths outside these boundaries in bcftools v1.6^60^. We further removed regions in the reference genome with more than one 35-kmer self-mapping using the SNPable tool with stringency 0.5^61^, as well as regions masked by a default run of RepeatMasker v4.0.6^62^. We also removed regions showing a high depth across multiple Lake Victoria region and outgroup cichlid species, where in 30 or more individuals’ depth exceeded mean depth plus 1.5 times interquartile range among >500 genomes^37,40^ aligned in the same way as outlined above. We further removed sites with a genotype showing quality below 30 or depth below 10 in any of the eleven individuals.

The second set of filters was intended to identify candidate *de novo* single base pair mutations for each parent offspring trio. Each retained candidate *de novo* single base pair mutation had to satisfy the following criteria: (i) homozygous reference genotype call in both parents, grandparents (only for F2 as grandparents not sequenced for F1 individuals) and same generation siblings. (ii) Parents not containing any reads supporting the alternate allele. (iii) Heterozygous genotype in the focal individual supported by >25% of the reads. (iv) Bi-allelic SNPs only. To exclude issues arising from mapping, structural variation or reference bias causing erroneous SNP calls or genotypes, each candidate *de novo* single base pair mutation detected by at least one of the three variant and genotype calling methods was manually curated into a list of final *de novo* mutations. This was done by visually inspecting the raw read alignments for all 11 individuals using IGV v2.5.3^63^.

We estimated the mutation rate from the final number of *de novo* single base pair mutations, divided by the product of (1) ploidy, (2) the total number of callable sites from the freebayes workflow and (3) the number of parent-offspring trios (i.e. generation transitions), i.e. μ = N_DeNovoMutations_ (2 * N_CallableSites_ * N_Trios_), with N_Trios_ being nine as we had two F0 vs. F1 trios and seven F1 vs. F2 trios, following Smeds *et al*.^59^. We estimated 95% confidence intervals for the number of mutations observed by computing the 2.5% and 97.5% quantile under a chi-square distribution with degrees of freedom 2 x N and 2 x (N+1) for lower and upper bounds, with N being the number of observed mutations^64,65^, and transforming those boundary mutation numbers to mutation rates with the same formula as outlined above.

### MSMC analysis

We used the HaplotypeCaller / GATK v3.5 approach to call variants and genotypes for the 112 wild-caught cichlids, generating base pair resolved GVCF files, followed by GenotypeGVCF creating VCF files containing monomorphic and variant sites. From this dataset, we used bcftools to remove genotypes covered by less than five reads, sites with more than 50% missing data, insertion / deletion polymorphisms and multiallelic sites. We also removed regions that overlap the masks outlined above and sites exceeding the mean site depth by 1.5 times the interquartile range within the dataset. In addition, sites with a quality-by-depth profile smaller than 2.0 (QD<2.0), with a Fisher strand bias greater than 50 (FS>50), a strands odds ratio greater than three (SOR>3.0), a mapping quality rank sum test smaller than -12.5 (MQRankSum<-12.5) and a read position rank sum test smaller than -8 (ReadPosRanksum<-8.0) were also excluded. We extracted phase information from reads with extractPIRs and applied read-backed phasing to SNPs in the dataset as implemented in shapeit v2.r900^66,67^. From combined phased SNPs and monomorphic sites, we generated BED files of sequenced sites and VCF files with biallelic SNPs for each individual. MSMC2 input for different combinations of individuals was generated using the generate_multihetsep.py script^68^.

We ran MSMC2 v2.1.2^68,69^ for different combinations and numbers of haplotypes specified with -I for 20 iterations^42,68^. We used a recombination over mutation rate parameter of -r 10 reflecting a magnitude lower mutation rate (see Main text) than recombination rate^51^ in Victoria cichlids. We converted the MSMC2 coalescence rates to time intervals and effective population sizes with the same mutation rate and assuming a two year generation time, as Lake Victoria haplochromine cichlids typically reach sexual maturity at one to two years of age, depending on their ecology, and live and reproduce for several years^18,70^. We computed relative cross-coalescence rates between species with the Python script combineCrossCoal.py^68^. We used the MSMC2 output to build isolation with migration models with MSMC-IM run with default parameters^43^ and to estimate effective population (N_e_) size within species, cumulative migration probabilities M(t) and migration rates m(t) between species. We fit a smoothing spline to M(t) estimates on a log10(time) x-axis using the R-function smooth.spline with smoothing parameter spar=0.4 to estimate the proportion of the genome of two species coalescing in a common ancestor at different time points with the R-function uniroot. We performed all analyses on the Euler computer cluster at ETH Zürich, Switzerland.

### Speciation intervals and rates

Given pervasive non-treelike evolution in the young adaptive radiations of the Lake Victoria region cichlids, we estimated speciation rates with a method not relying on a resolved phylogenetic tree. Instead we assumed either a linear or a logarithmic diversification model following McCune^71^, leading to maximum and minimum bounds for speciation rate^24,71^, respectively. The linear model assumes a maximally unbalanced tree with all speciation along a single branch leading to a maximum per tip speciation rate estimate, while the logarithmic model assumes a maximally balanced tree where every branch in the tree splits again leading to a minimum per tip speciation rate estimate^71^. We used the number of known species in each cichlid radiation^16–20,72^ and the oldest time estimate for when the majority of the genome of each species pair has coalesced in an common ancestor in a radiation (Table 1), assuming a single ancestral species^26^ (N_0_=1), to compute net speciation intervals (SI), the average time between two successive speciation events along each branch, as SI_lin_ = t / (N_t_ – N_0_) and SI_log_ = (t * log_e_(2)) / (log_e_(N_t_) – log_e_(N_0_)). We converted these speciation intervals to speciation rates (λ) reported in number of speciation events along a branch per million years: λ = 1,000,000 / SI. We did not estimate SI and λ for Lake Saka, as it contains only two species with much uncertainty in divergence time.

### Trophic evolution, habitat and effective population size

We analyzed the relationship between trophic position, habitat and most recent effective population size across the LVRS radiations in an ‘evolutionary comparative’ Bayesian framework – adopting phylogenetic comparative methods while relaxing the assumption of tree-like evolution. We restricted the analysis to members of the large LVRS adaptive radiations Victoria, Albert, Edward and Kivu, and used Upper Nile cichlids as an outgroup. To incorporate the effects of non-independence due to shared evolutionary history, we generated a covariance matrix from MSMC-IM time estimates at 50% to 80% probability of the species pairs having merged in a common ancestor. We used estimates from two haplotypes for all comparisons except for within Lake Victoria comparisons, for which six haplotype estimates were available. We used the ultrametric function in the R package ape v5.5^73^ to impute six missing time estimate too young to be accurately estimated with MSMC2, with all imputed values ranging from 2,307 to 3,528 years. We divided the matrix by the maximum time estimate and subtracted it from one and used the function nearPD from the R package Matrix v1.3 to compute the nearest approximate positive definite matrix. We used the Bayesian framework implemented in STAN via the R package brms v2.16.1^74,75^. We included log10-transformed most recent effective population size estimates as dependent variable, habitat (littoral, open water) as a binary and trophic position (primary consumers: herbivores, secondary consumers: insectivores, zooplanktivores, molluscivores, tertiary consumers: piscivores and paedophages) as monotonic predictors. We used normally distributed priors with mean zero and standard deviation five for intercept and effect sizes of habitat and trophic position, and exponentially distributed priors with a rate parameter of 20 for sigma (σ) and a rate parameter of 10 for σ’s standard deviation (SD). σ and SD were used to estimate a non-linear Bayesian model equivalent to Pagel’s λ^76^, capturing how much shared evolutionary history, approximated by the covariance matrix, explains the response variable effective population size, with the hypothesis that σ^2^ / (σ^2^+ SD^2^) = 0.

## Acknowledgments

We would like to thank Pamela Nicholson, Cord Drögemüller, and Tosso Leeb from the sequencing platform at the University of Bern, Switzerland, for whole genome sequencing services and support, as well as Niklaus Zemp and Aria Minder from the Genetic Diversity Center at ETH Zürich, Switzerland, for bioinformatics support. We acknowledge TAFIRI for hosting O.S. and team members during fieldwork in Tanzania, M. Kayeba, M. Haluna and H. Mrosso for assistance with Tanzanian fieldwork, and NaFiRRI for hosting O.S. and M.D.M. during fieldwork in Uganda, and the Makerere University Biological Field Station for hosting O.S., L.J.C and C.A.C in Uganda. We thank Guy Periat and his team for sampling in Lake Kivu. We thank the Aquatic Ecology and Evolution group at the University of Bern Institute of Ecology & Evolution and the Fish Ecology and Evolution Department at the Eawag for discussion and feedback at all stages of the project. This research was funded by Swiss National Science Foundation grant 31003A_163338 to O.S., L.E. and R. Bruggmann, and by National Science Foundation grant DEB-1556963 to C.E.W.

