## Supplementary Information for "Genomes reveal age and demographic consequence of ultrafast adaptive radiation"

Marques et al.

### Supplementary Notes

*Supplementary Note 1: Phylogenetic estimates of LVRS and Victoria cichlid ages*

Previous phylogenetic studies estimated vastly different times to the most recent common ancestor (TMRCA) of all LVRS cichlids and all Victoria cichlids, probably due to different molecular clock assumptions and sometimes erroneous inclusion of outgroup taxa to the Victoria or LVRS radiations. The oldest TMRCAs of 4.7 MY (95% confidence interval 3.2-6.3 MY) for the LVRS cichlids and 3.0 MY (2.3-3.9 MY) for the Victoria cichlids were estimated from microsatellite data[1], but the study included the taxon *Astatoreochromis alluaudi* into both the Victoria and LVRS lineages, which has previously[2,3] been found and reconfirmed[4,5] to be a distantly related outgroup. The second oldest TMRCA of 716,000 years for the LVRS cichlids was estimated from the nuclear ITS-1 intron sequence based on one substitution rate assumption, while another substitution rate assumption for the same gene suggested a TMRCA of 10,000 years[2]. The earliest study, using two mitochondrial genes and a rate assumption for the mitochondrial cytochrome b gene, estimated a TMRCA of less than 200,000 years for the Victoria cichlids[3]. Faster evolving mitochondrial control region sequences used in another study and a substitution rate calibrated on the geological origin of Lake Malawi[6], led to a TMRCA estimate of 98,000 to 132,700 years for Victoria cichlids[7]. A third study used one nuclear and two mitochondrial genes and estimated a TMRCA for LVRS cichlids to 273,000 (±216,000) years and 189,000 (±133,000) years, respectively, when calibrated either on the assumption of an origin of all cichlids before the Gondwana breakup or on assumptions from the scant cichlid fossil record[8]. Under these two assumptions, the TMRCA of the Victoria cichlids was 120,000 (±110,000) and 89,000 (±74,000) years[8], respectively. Another study using the mitochondrial control region and SINE insertion patterns and three Victoria cichlid species from the genus *Yssichromis* estimated a TMRCA of 210,000 and 30,000 years for these three species[9] under the same two calibration assumptions[8], but also estimated a TMRCA of 732,000 years[9] under a second mutation rate[10]. In all of these studies, the non-linearity of molecular clocks in the past 1-2 million generations[11] likely led to vast overestimation of very recent times[4], reflected in the large range of estimates as well as in their large confidence intervals where given.

*Supplementary Note 2: Mutation rate estimation*

In the dataset of 11 whole genomes of a three-generation family (Supplementary Fig. 27) of the Lake Victoria cichlid *Gaurochromis hiatus*, 577,885,642 callable sites passed stringent quality filters (see Methods) and of 596 candidate *de novo* single base pair mutations in parent-offspring trios, 25 *de novo* single base pair mutations remained after manual curation of read alignments (Supplementary Table 2).

Nine *de novo* mutations are confirmed F0 germline mutations transmitted to one to five of the seven F2 individuals, while the remaining 16 are likely germline mutations in F1 individuals. Six mutations are transversions, resulting in a transversion to transition ratio of 0.23. Three mutations are C to T substitutions at CpG sites (Supplementary Table 2). Dividing the number of observed mutation events (n=25) by the product of organism ploidy (n=2), number of parent-offspring trios (n=9) and number of callable sites (n=577,885,642) resulted in an estimate of $2.4 \times 10^{-9}$ single base pair mutations per generation and base pair with 95% confidence interval $1.6 \times 10^{-9}$ to $3.5 \times 10^{-9}$, comparable to other pedigree-derived haplochromine cichlid mutation rates[12]. The corresponding mutation rate excluding C to T substitutions at CpG sites is $2.1 \times 10^{-9}$ single base pair mutations per generation and base pair with 95% confidence interval $1.3 \times 10^{-9}$ to $3.2 \times 10^{-9}$.

*Supplementary Note 3: Order of lineage divergence in Lake Victoria*

The Victoria paedophages (represented by the genus *Lipochromis*), piscivores (*Harpagochromis*), snail crushers (*Labrochromis*), and anatomically specialized insect pickers (*Paralabidochromis chilotes*) merge most anciently in a common ancestor among Victoria cichlids with 50% probability of having merged between 5,816 and 14,139 years ago, suggesting that these lineages diverged first in the explosive Victoria radiation (Supplementary Fig. 1). Between the anatomically specialized guilds of rocky shore cichlids, such as epilithic algal browsers (*Neochromis*), epilithic algal grazers (*Mbipia*), reef insectivores and reef planktivores (*Pundamilia*), as well as oral shelling molluscivores (*Macropleurodus*), the 50% probability of having merged in a common ancestor is 3,096-7,166 years ago. Most recently, the 50% probability of having merged in a common ancestor between detritivores' (*Enterochromis*) and pelagic zooplanktivores' (*Yssichromis*) is 2,232 years ago, while among rocky shore cichlids, the 50% probability is reached 2,293-6,557 years ago (Supplementary Fig. 1). The recent emergence of open water guild members coincides with an increase of haplochromine cichlids in offshore habitat between 3,800 and 2,300 years ago inferred from fossil cichlid teeth found in sediment cores[13]. This order of divergence among representatives of different guilds is consistent with the adaptive radiation in Lake Victoria continuing to generate new clades that subsequently evolved into new trophic guilds throughout its rapid evolutionary history, as proposed by theory[14] and supported by the evolutionary history of novel, endemic guilds in the Victoria radiation[5].

Speciation in the Victoria radiation continued within guilds into the 500 species currently known[15–18]. We estimated the divergence of two of what we expected to be among the most recent speciation events, for two pairs of congeneric species: *Pundamilia* sp. "pundamilia-like"/*P.* sp. "nyererei-like" and *Neochromis greenwoodi*/*N. omnicaeruleus*. In both congeneric species pairs, the 50% probability of lineages having merged in a common ancestor falls into the last 1,200 years (Supplementary Fig. 28), below the inference limit of the MSMC-IM method and thus not in conflict with a population-based demographic modelling estimate of 314 years (95% confidence interval: 134-642 years) divergence time for the former of these species pairs[19].

*Supplementary Note 4: Admixture and age of founding lineages*

The Congo and Upper Nile parental lineages of the LVRS show most lineage merging during two migration episodes 40,000-100,000 years and 400,000-800,000 years ago, with complete merging of lineages during a third migration episode around 2 MY ago and a plateau in M(t) between these episodes (Fig. 2b, d, Supplementary Fig. 29). The most recent migration episode 40,000-100,000 years ago overlaps with the complete lineage merging of all LVRS lineages (Fig. 2b, d, Supplementary Fig. 29), as well as with the period of most rapid lineage merging between either Congo or Upper Nile cichlids with all LVRS lineages 30,000-80,000 years ago (Fig. 2b, d, Supplementary Fig. 30-31). This observation suggests that the LVRS hybrid swarm[4] was founded 30,000-80,000 years ago, during which gene exchange also occurred between the founding Congo and Upper Nile lineages. The second episode 400,000-800,000 years ago coincides with the initial formation of Lake Victoria 400,000 years ago and the establishment of the modern Lake Victoria region drainage system changing outflow from the Congo to the Nile[20]. This would likely have been the first time that larger faunal exchange became possible between the old drainage of Paleolake Obweruka (the modern Upper Nile region) and the central-western Tanzanian Malagarasi / Upper Congo drainage. The third episode around 2 MY ago corresponds well with phylogenomic divergence times estimates of 1.63-5.79 MY between Congo and Upper Nile lineages based on molecular clocks calibrated with paleolimnological constraints and non-cichlid fossils[4]. The two more recent migration episodes thus likely reflect repeated admixture in secondary contact after the Upper Nile and Congo lineages first split ~2 MY ago.

The distantly related outgroup *Astatoreochromis alluaudi*, a lineage that did not radiate in the Lake Victoria region despite being widely-distributed, starts to merge with all other lineages steadily before 60,000 years ago and most rapidly 2 MY ago, with a minor migration episode around 50,000-80,000 years and a major migration episode 2-6 MY ago (Fig. 2b,d, Supplementary Fig. 32), suggesting a split time several million years ago, consistent with phylogenetic estimates[4,21], and surprisingly, some limited, more recent gene flow with the lineages that founded the LVRS radiations (Fig. 2b,d).

*Supplementary Note 5: Parallel adaptive radiations*

Our results are consistent with parallel adaptive radiations[5] in Lake Victoria and multiple Western rift lakes: genomes coalesce more recently within Lake Victoria and within each Western Rift lake than between Lake Victoria and Western rift lakes (Fig. 3, Supplementary Fig. 6-26). The monophyly of Lake Victoria cichlids in phylogenomic analyses and PCA grouping are consistent with adaptive radiations evolving in parallel[4,5]. Lineage merging between Lake Edward and Lake Kivu species however overlaps broadly with lineage merging within these lakes (Supplementary Fig. 6), suggesting that these two assemblages share their evolutionary origin in a single adaptive radiation that unfolded during or shortly after the arid period 16,700-20,200 years ago. A shared Kivu / Edward radiation is in line with the paraphyly of these assemblages relative to each other in phylogenetic analyses[4,5]. Lineage merging between Lake Albert species is more recent than with Edward and Kivu species (Supplementary Fig. 6), suggesting that the Lake Albert radiation is another radiation. Likewise, the Lake Saka species pair merges more recently in a common ancestor than with other Western rift lake species or Lake Victoria species (Supplementary Fig. 6), with the next most recent lineage merging being with Lake Edward species, consistent with the hypothesis that an Edward ancestor founded this small crater lake radiation. In conclusion, the adaptive radiations of haplochromine cichlids in Lake Victoria, Lake Albert, Lake Saka and the combined Lakes Edward / Kivu evolved in largely in parallel, in line with phylogenomic and earlier population genomic analyses [4,5], as thus did their trophic networks.

*Supplementary Note 6: Demography pre-radiation*

All lake endemic LVRS members and the Upper Nile lineage show similar effective population size ($N_e$) trajectories prior to radiations in each lake, with a large ancestral population size of ~200,000 individuals decreasing to smaller sizes beginning some 100,000-200,000 years ago (Figs. 3c-f). This large ancestral population size estimate might be inflated due to population structure arising from the hybrid origin of all LVRS and Victoria radiations[4,5]. The formation of the LVRS hybrid swarm combined two divergent gene pools into one, and as such, 'population structure' between these pools should lead to inflated estimates of ancient $N_e$ due to more ancient coalescence of many haplotypes than expected in a non-structured population lacking ancient admixture and hybrid origins[22]. This bias in the estimation of ancestral $N_e$ does however not apply to the most recent estimates of $N_e$ for single species in each radiation, nor would such a bias affect the hierarchy of recent effective population size associated with trophic level and habitat (see Main Text, Fig. 3).

### Supplementary Figures

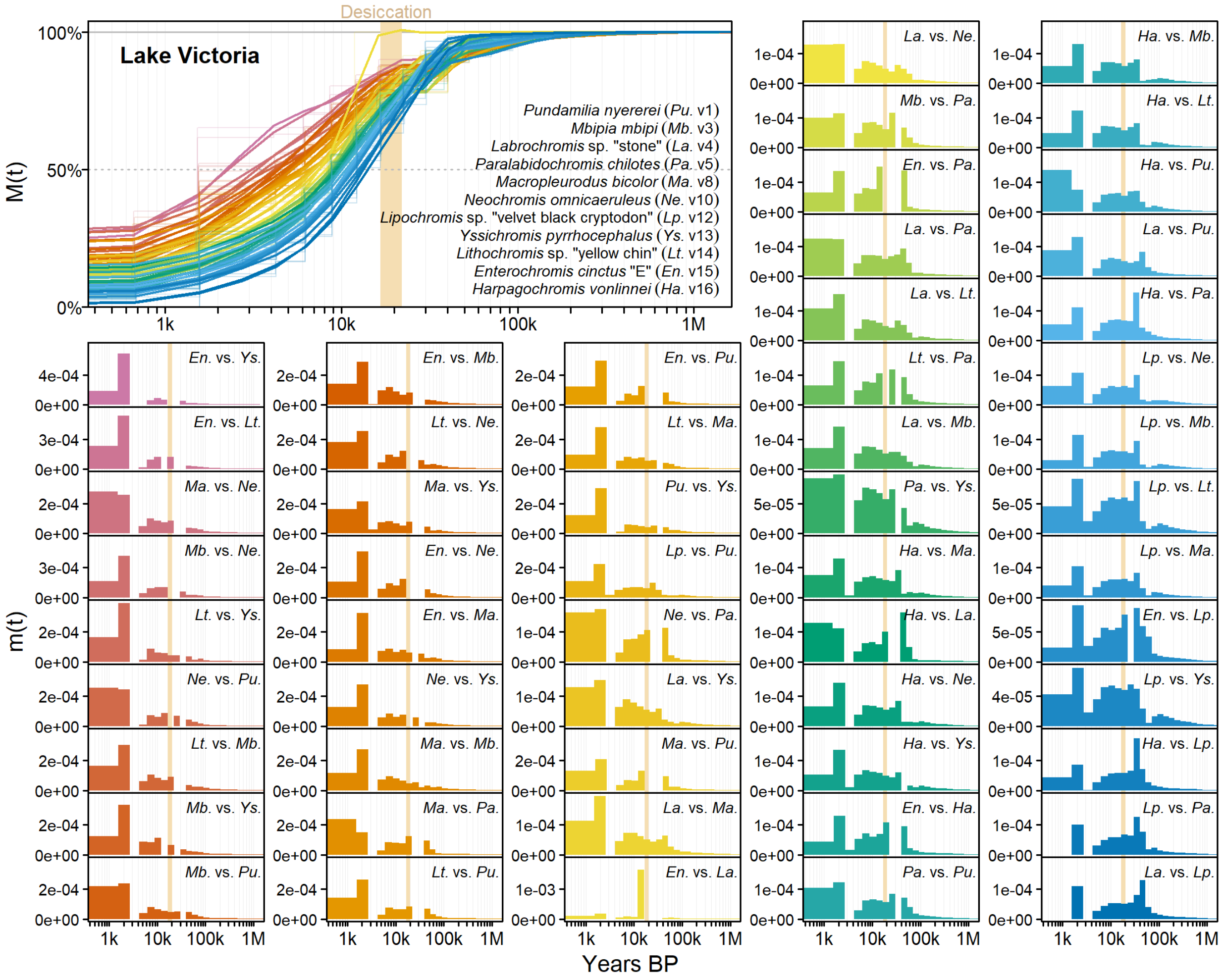

**Supplementary Fig. 1 | Observed linage merging between haplochromine cichlid species from all major guilds of the Lake Victoria radiation.** Cumulative migration probability M(t) and migration rates m(t) between species estimated from six haplotypes per species support the hypothesis of an explosive radiation in Lake Victoria: among all major guilds representatives of the Victoria radiation the probability of merging in an ancestral lineage exceeds 50% in the last 16,700 years, when the modern lake refilled (lake desiccation 16,700-20,200 years ago indicated with a beige vertical bar). Migration rate distributions suggest that guilds might not have arisen simultaneously, with paedophages, predators and shell-eaters (*Lipochromis*, *Harpagochromis*, *Labrochromis*) merging deeper in the past than rock cichlids (*Neochromis*, *Mbipia*, *Pundamilia*, *Macropleurodus*) and most recently open water guild cichlids (*Enterochromis*, *Yssichromis*).

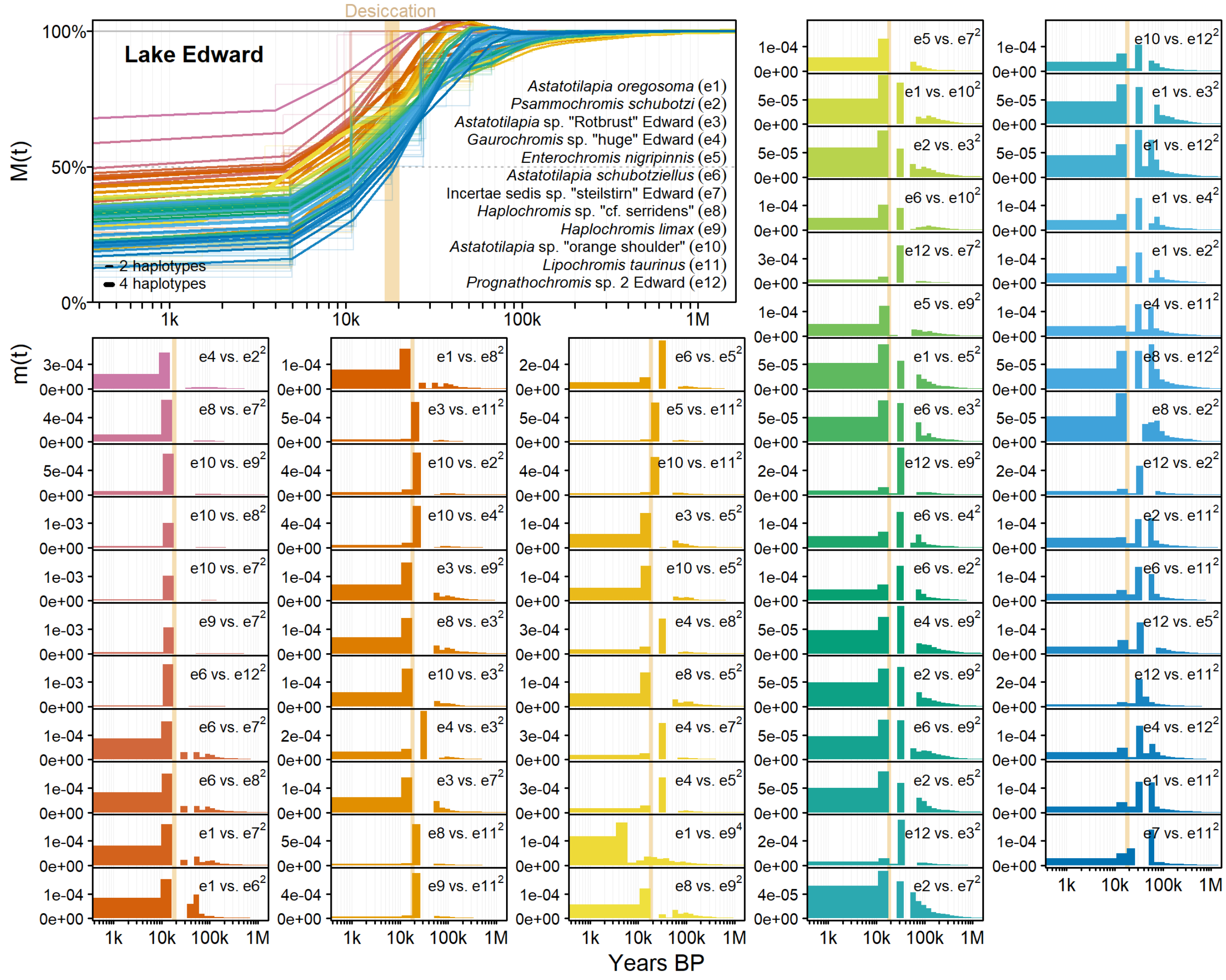

**Supplementary Fig. 2 | Observed linage merging between haplochromine cichlid species from Lake Edward.** Similarly to Lake Victoria, lineage merging among 12 representatives of various guilds from Lake Edward is fastest around the arid period that led to the desiccation of Lake Victoria 16,700-20,200 years ago (indicated with beige vertical bar), consistent with a recent, explosive origin of the shared Edward / Kivu radiation. Line widths and superscripts behind species numbers indicate whether two or four haplotypes per species were used to estimate cumulative migration probability M(t) and migration rates m(t).

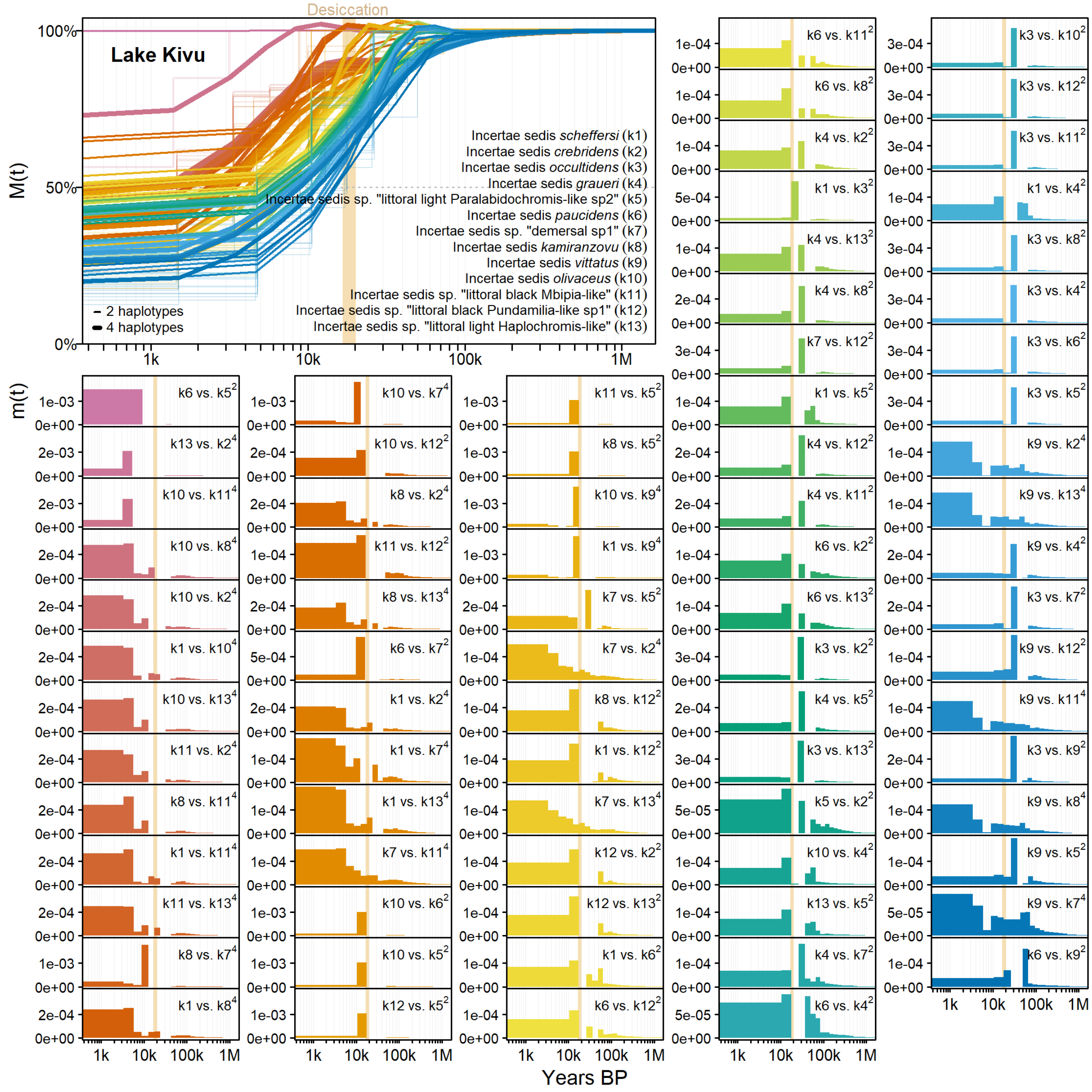

**Supplementary Fig. 3 | Observed linage merging between haplochromine cichlid species from Lake Kivu.** Similarly to Lake Victoria, lineage merging among 13 representatives of various guilds from Lake Kivu is fastest around the arid period that led to the desiccation of Lake Victoria 16,700-20,200 years ago (indicated with beige vertical bar), consistent with a recent, explosive origin of the shared Edward / Kivu radiation. Line widths and superscripts behind species numbers indicate whether two or four haplotypes per species were used to estimate cumulative migration probability M(t) and migration rates m(t).

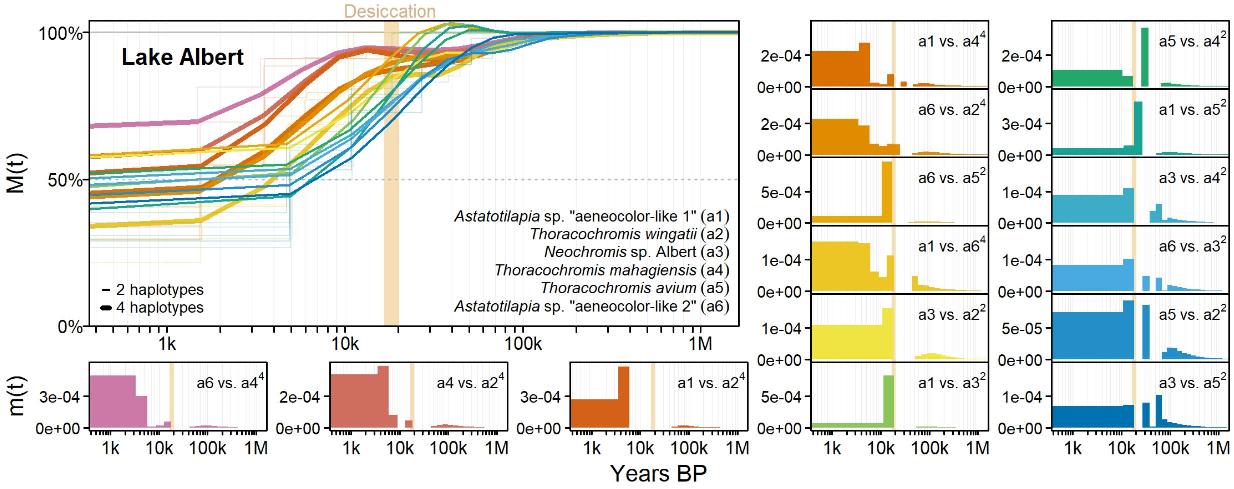

**Supplementary Fig. 4 | Observed linage merging between haplochromine cichlid species from Lake Albert.** The six representatives of various guilds from Lake Albert merge much more recently in a common ancestor than members of the Victoria and Kivu / Edward radiations, suggesting a very recent origin of the Lake Albert haplochromine radiation, despite the lake not desiccating during the arid period 16,700-20,200 years ago (indicated with beige vertical bar). Line widths and superscripts behind species numbers indicate whether two or four haplotypes per species were used to estimate cumulative migration probability M(t) and migration rates m(t).

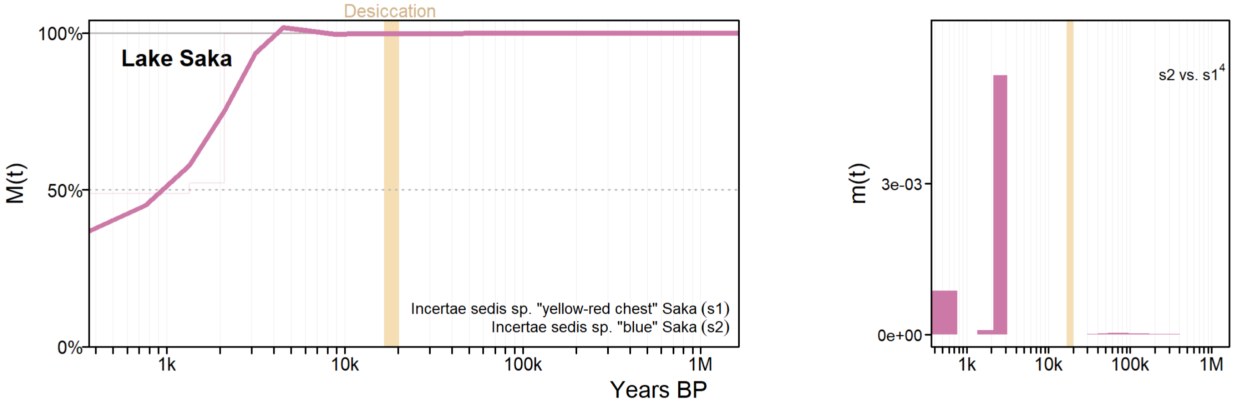

**Supplementary Fig. 5 | Observed linage merging between haplochromine cichlid species from Lake Saka.** The two Lake Saka species merge very recently into a common ancestor, in the last 1,000 years, suggesting recent speciation in this young ecotype pair independent of the arid period that led Lake Victoria to desiccate 16,700-20,200 years ago (beige vertical bar). This estimate is at the lower inference limit of the MSMC-IM method, even with reconstructing cumulative migration probability M(t) and migration rates m(t) with eight haplotypes per species.

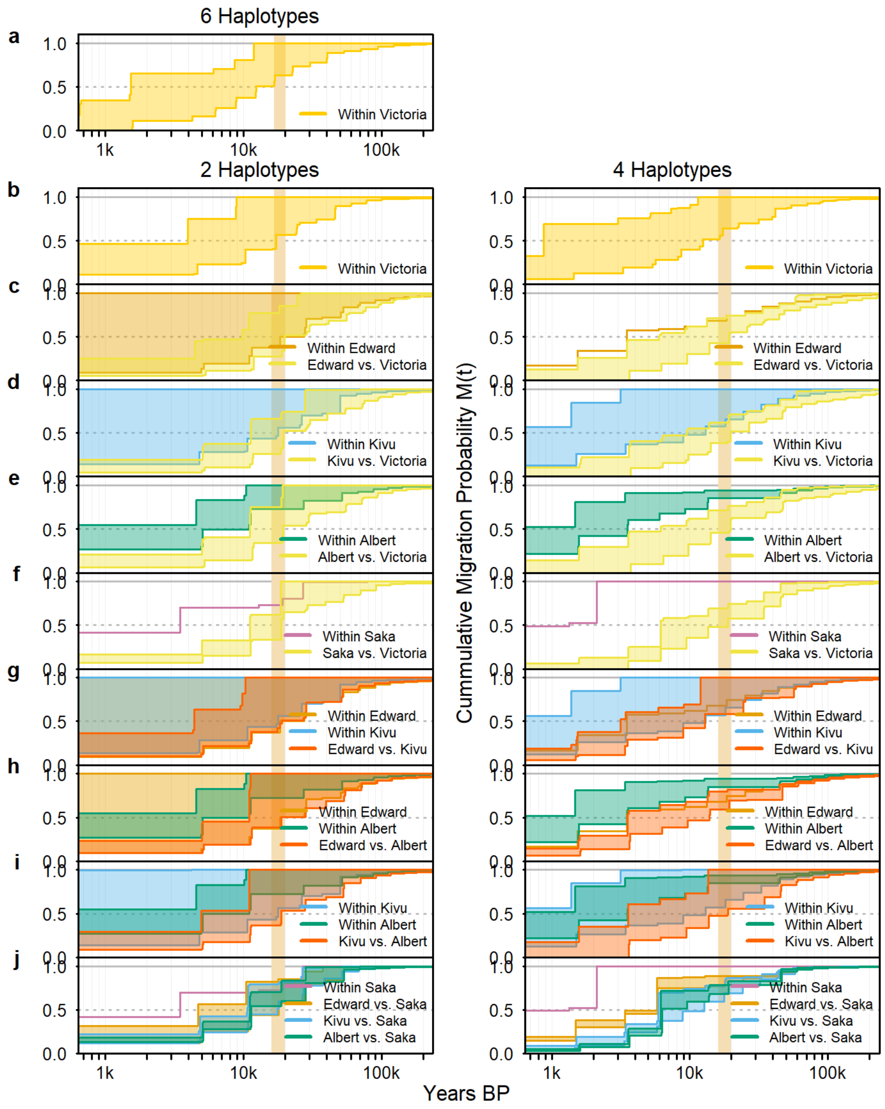

**Supplementary Fig. 6 | Lineage merging between haplochromine cichlid species from the same and different lakes in the Lake Victoria region.** The distribution of cumulative migration probability M(t) curves between all species comparisons from the same ("within") or different ("vs.") lakes in the Lake Victoria region is consistent with parallel, largely independent radiations in lakes Victoria, Albert, Saka and one shared radiation in Kivu and Edward. Lineage merging between and within lake broadly overlaps among the latter two lakes (**g**), while lineage merging is more ancient between than within lake in other comparisons (**c-f**, **h-j**). The distinction is more prominent for estimates based on four haplotypes (right), than two haplotypes (left), as more recent times are better resolved, while the four haplotype comparisons contain fewer species comparisons. Lineage merging between Victoria and Western rift lake radiations around the arid period that desiccated Lake Victoria 16,700-20,200 years ago (beige vertical bar) is consistent with recent genetic exchange before the radiations unfolded[5].

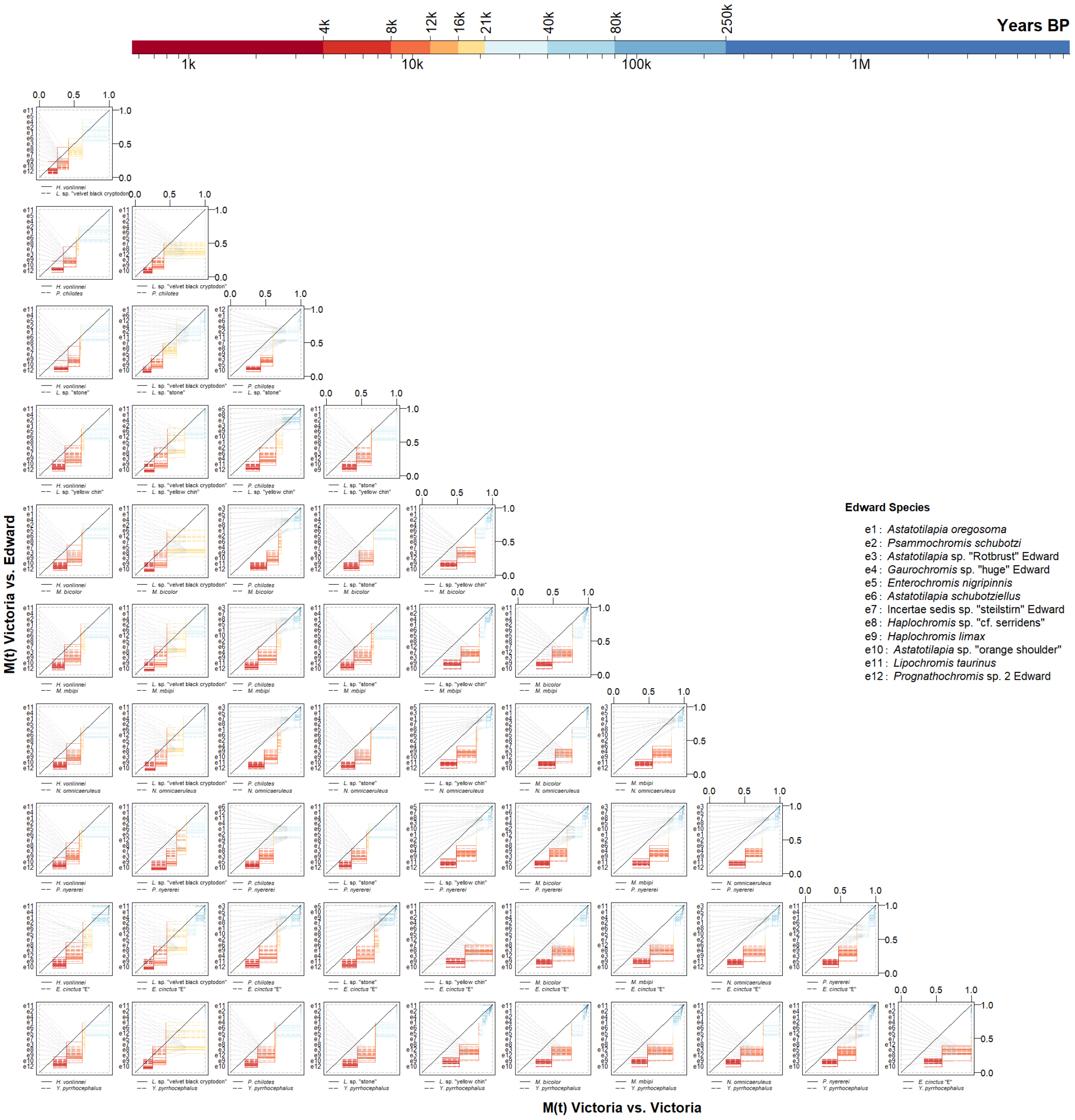

**Supplementary Fig. 7 | Observed lineage merging among Lake Victoria cichlids from the same lake vs. from Lake Edward, resolved by species.** Cumulative migration probability M(t) trajectories reconstructed from two haplotypes per species are shown for species pairs from within and between lakes. The color scale gives the timing of lineage merging.

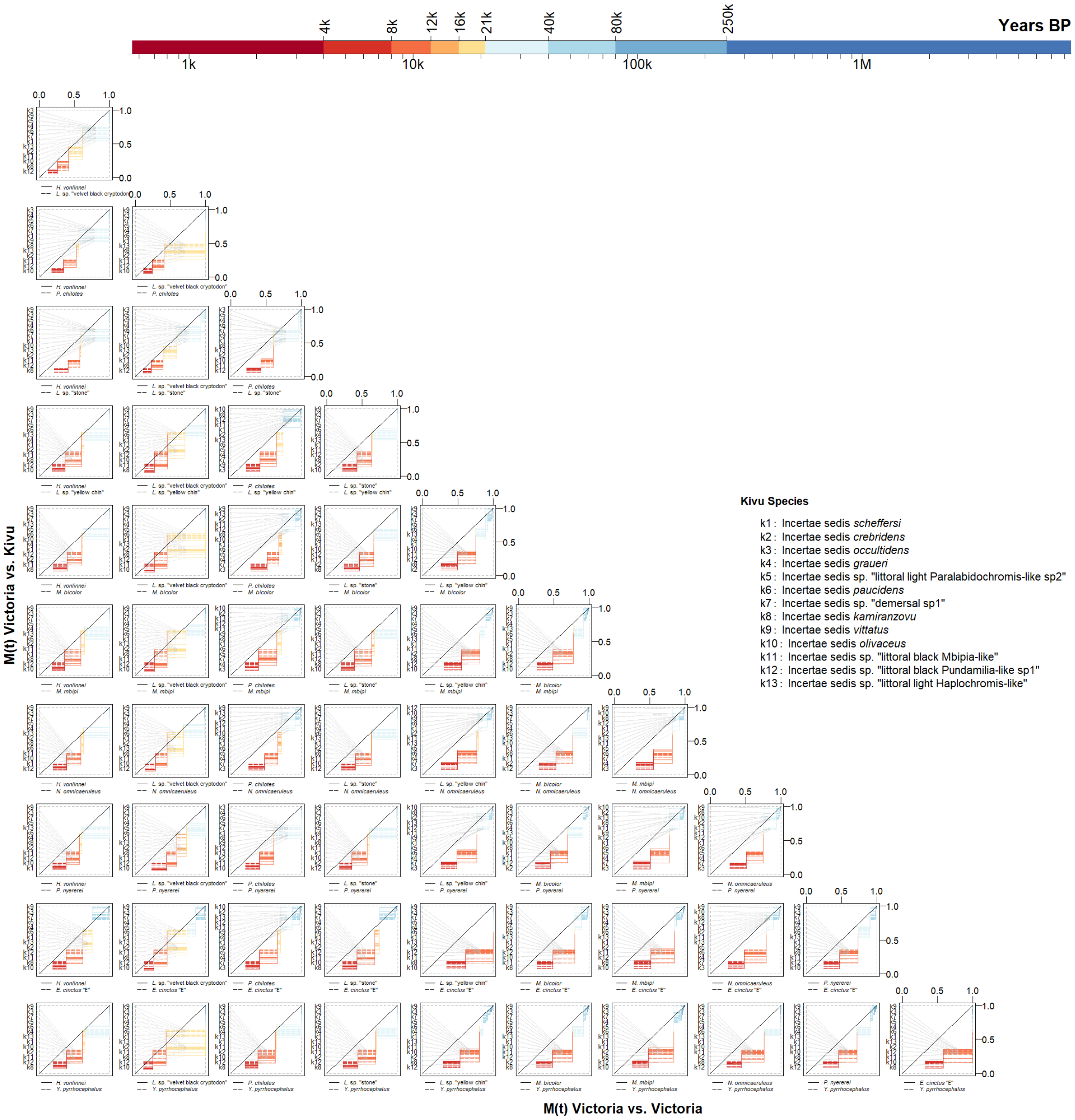

**Supplementary Fig. 8 | Observed lineage merging among Lake Victoria cichlids from the same lake vs. from Lake Kivu, resolved by species.** Cumulative migration probability M(t) trajectories reconstructed from two haplotypes per species are shown for species pairs from within and between lakes. The color scale gives the timing of lineage merging.

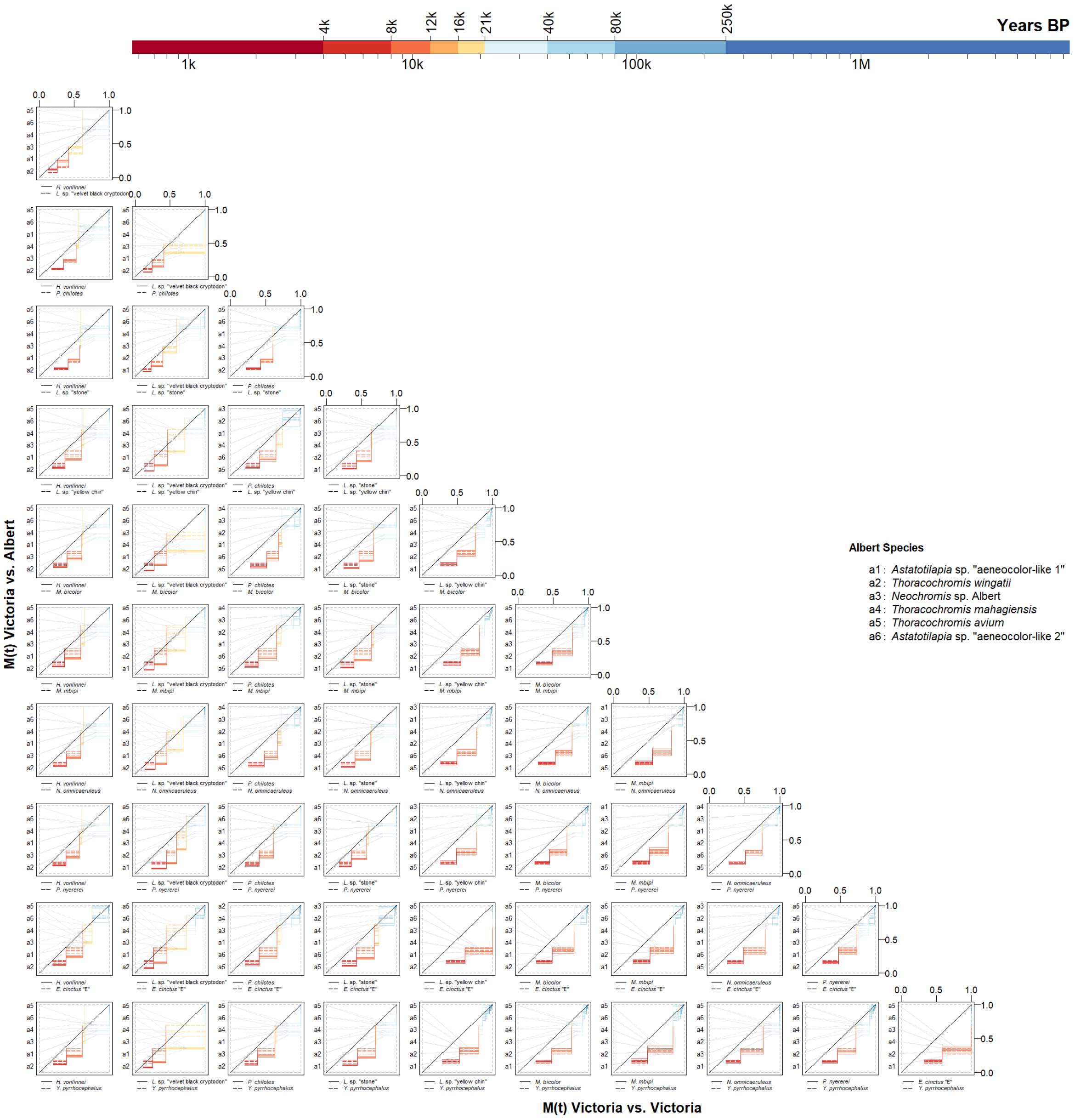

**Supplementary Fig. 9 | Observed lineage merging among Lake Victoria cichlids from the same lake vs. from Lake Albert, resolved by species.** Cumulative migration probability M(t) trajectories reconstructed from two haplotypes per species are shown for species pairs from within and between lakes. The color scale gives the timing of lineage merging.

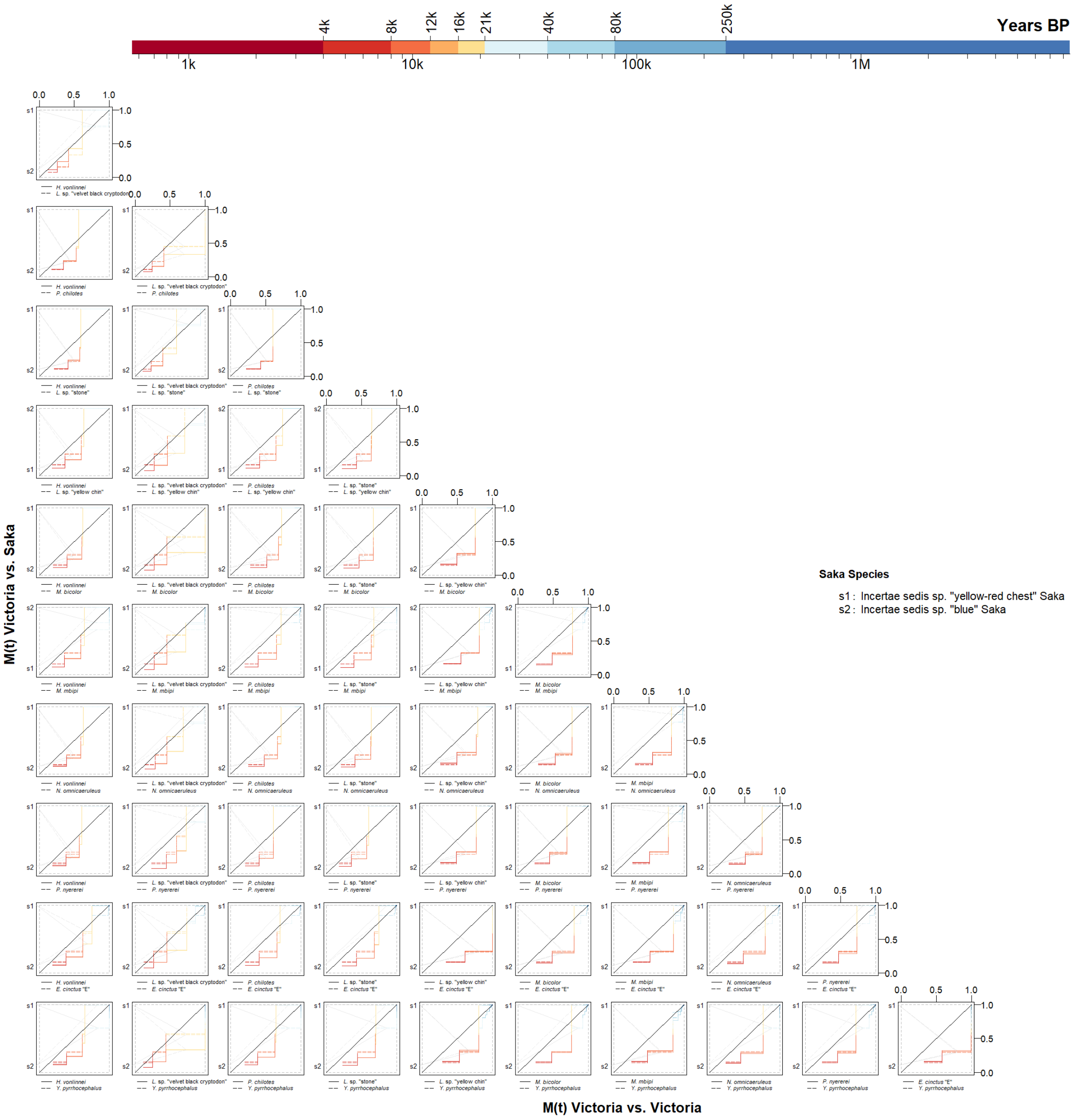

**Supplementary Fig. 10 | Observed lineage merging among Lake Victoria cichlids from the same lake vs. from Lake Saka, resolved by species.** Cumulative migration probability M(t) trajectories reconstructed from two haplotypes per species are shown for species pairs from within and between lakes. The color scale gives the timing of lineage merging.

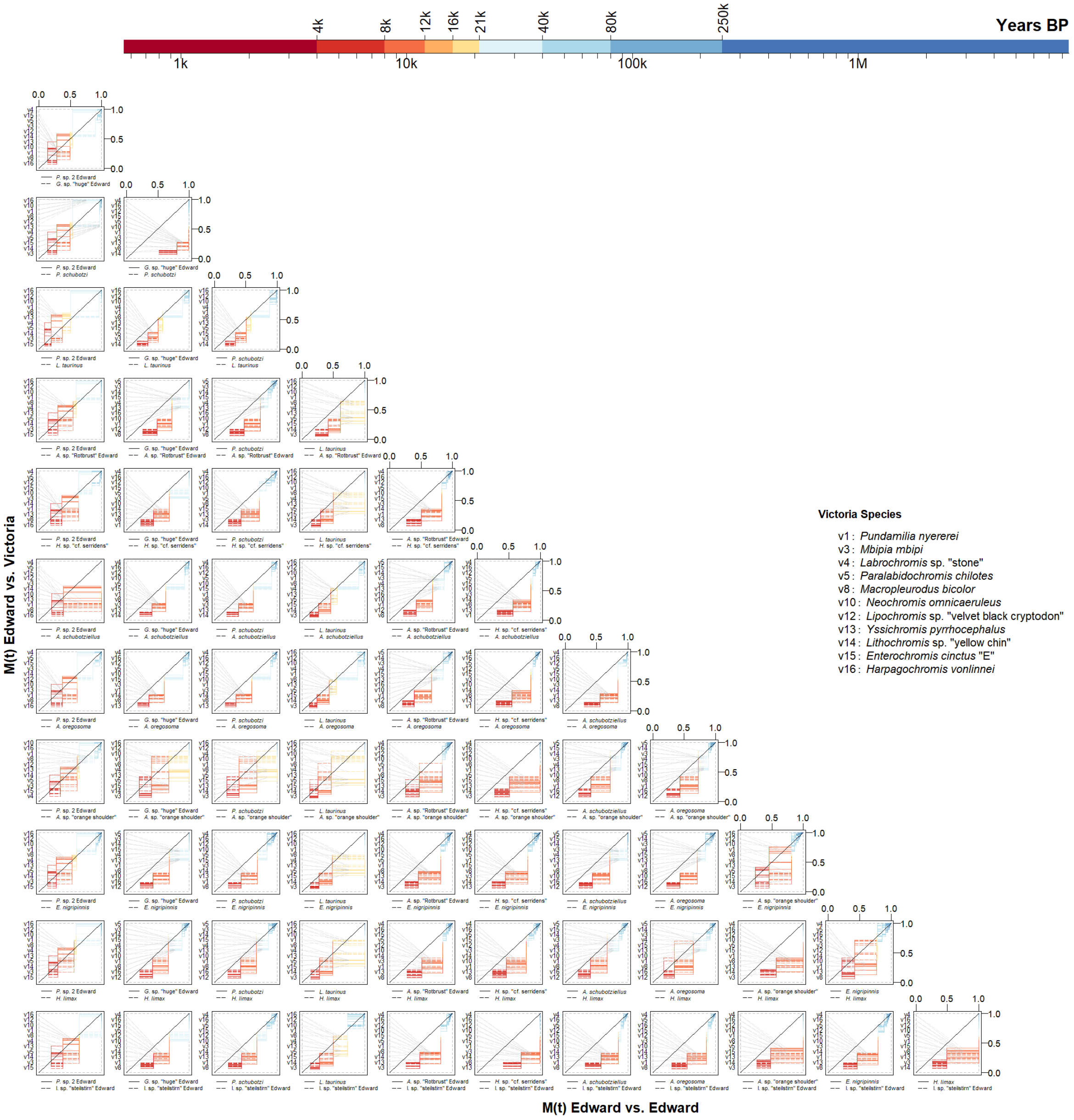

**Supplementary Fig. 11 | Observed lineage merging among Lake Edward cichlids from the same lake vs. from Lake Victoria, resolved by species.** Cumulative migration probability M(t) trajectories reconstructed from two haplotypes per species are shown for species pairs from within and between lakes. The color scale gives the timing of lineage merging.

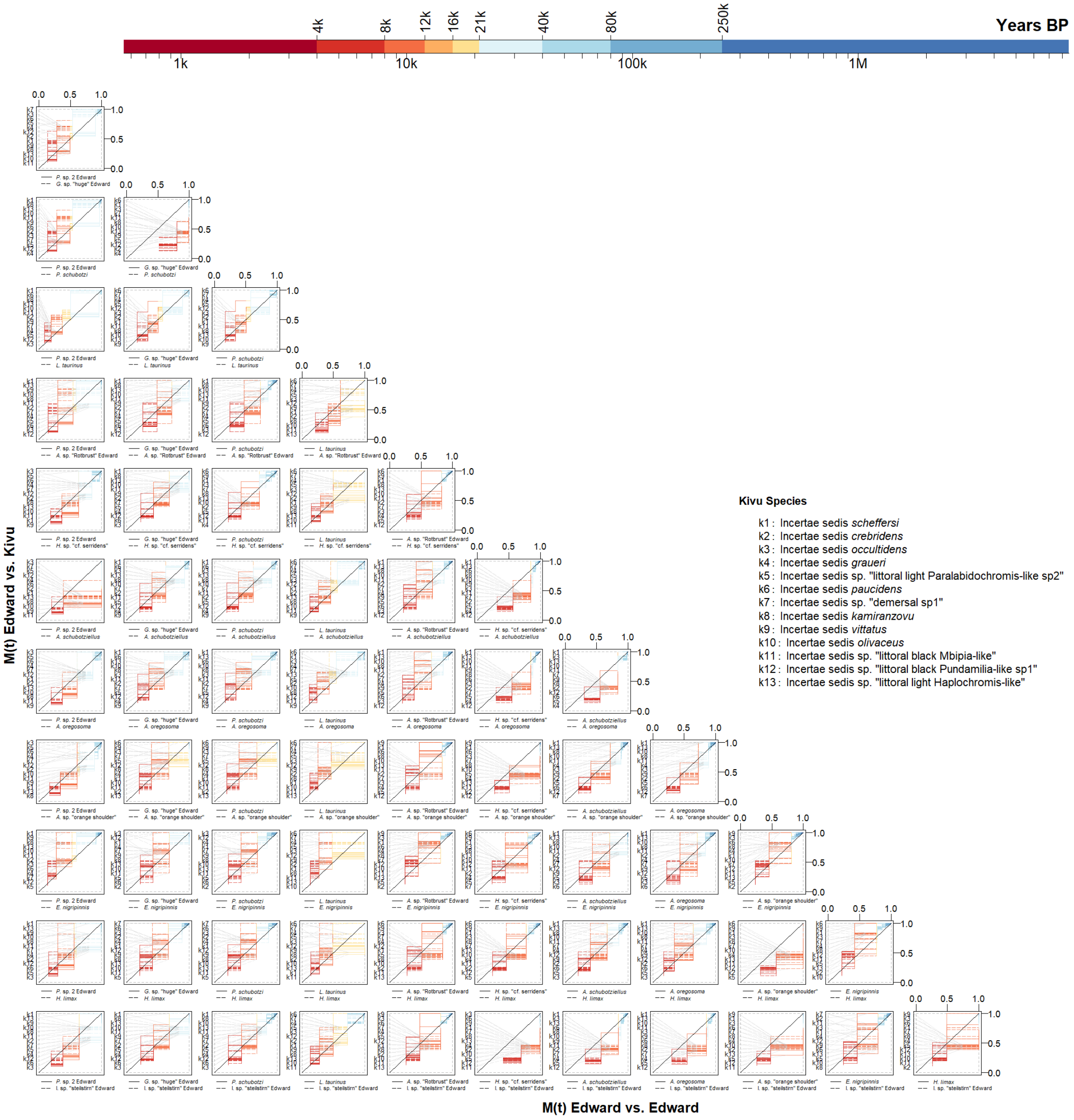

**Supplementary Fig. 12 | Observed lineage merging among Lake Edward cichlids from the same lake vs. from Lake Kivu, resolved by species.** Cumulative migration probability M(t) trajectories reconstructed from two haplotypes per species are shown for species pairs from within and between lakes. The color scale gives the timing of lineage merging.

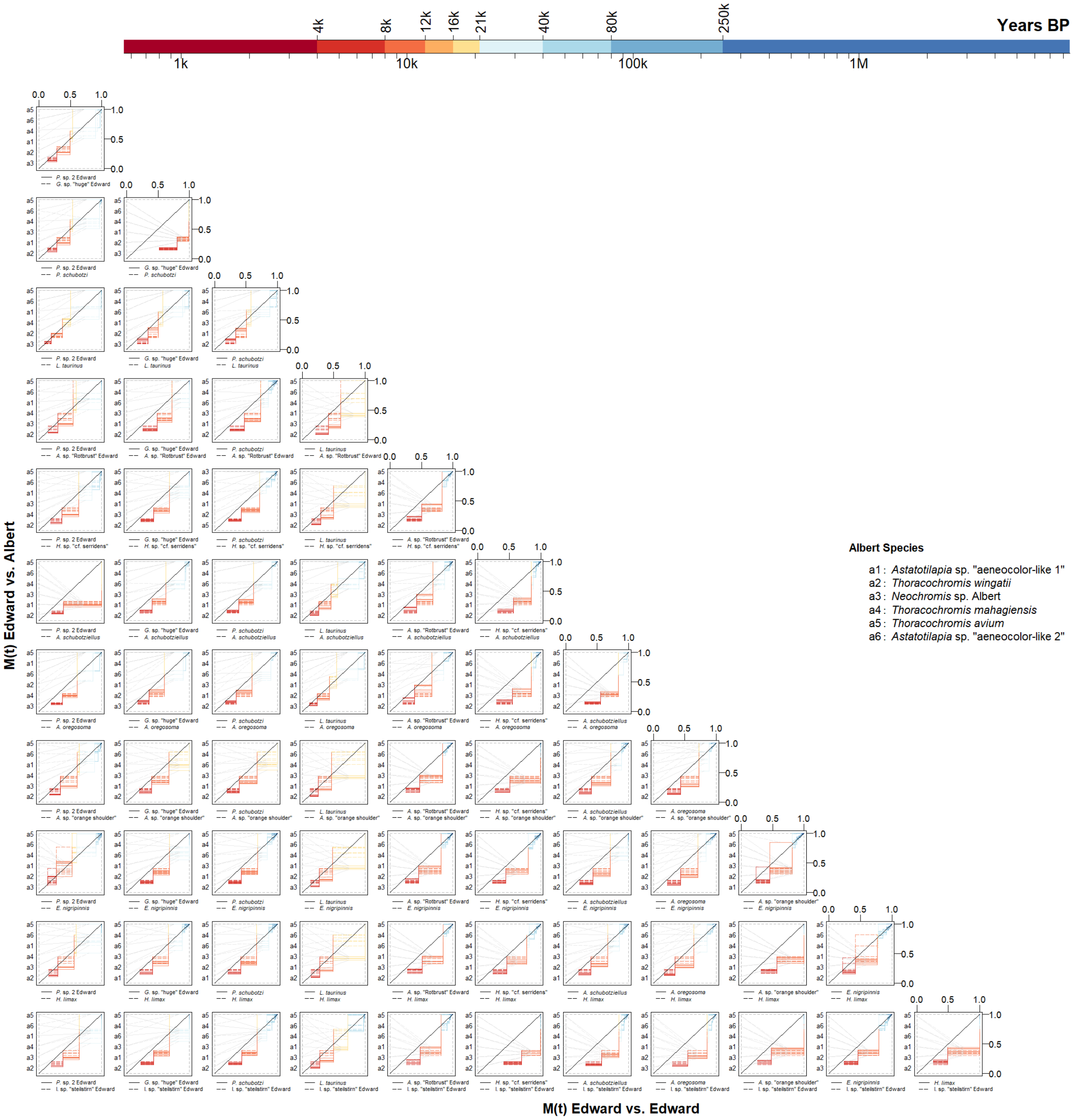

**Supplementary Fig. 13 | Observed lineage merging among Lake Edward cichlids from the same lake vs. from Lake Albert, resolved by species.** Cumulative migration probability M(t) trajectories reconstructed from two haplotypes per species are shown for species pairs from within and between lakes. The color scale gives the timing of lineage merging.

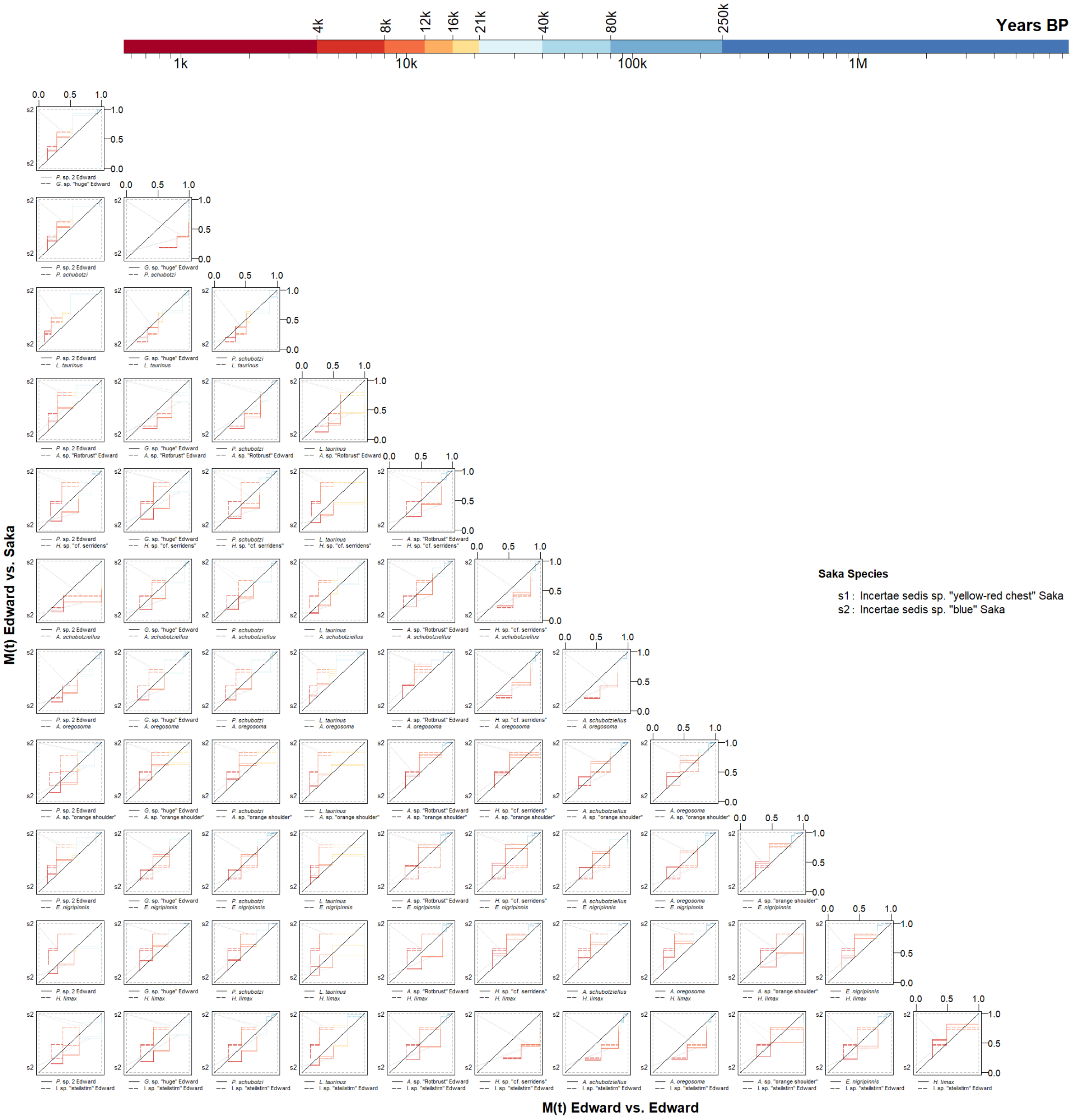

**Supplementary Fig. 14 | Observed lineage merging among Lake Edward cichlids from the same lake vs. from Lake Saka, resolved by species.** Cumulative migration probability M(t) trajectories reconstructed from two haplotypes per species are shown for species pairs from within and between lakes. The color scale gives the timing of lineage merging.

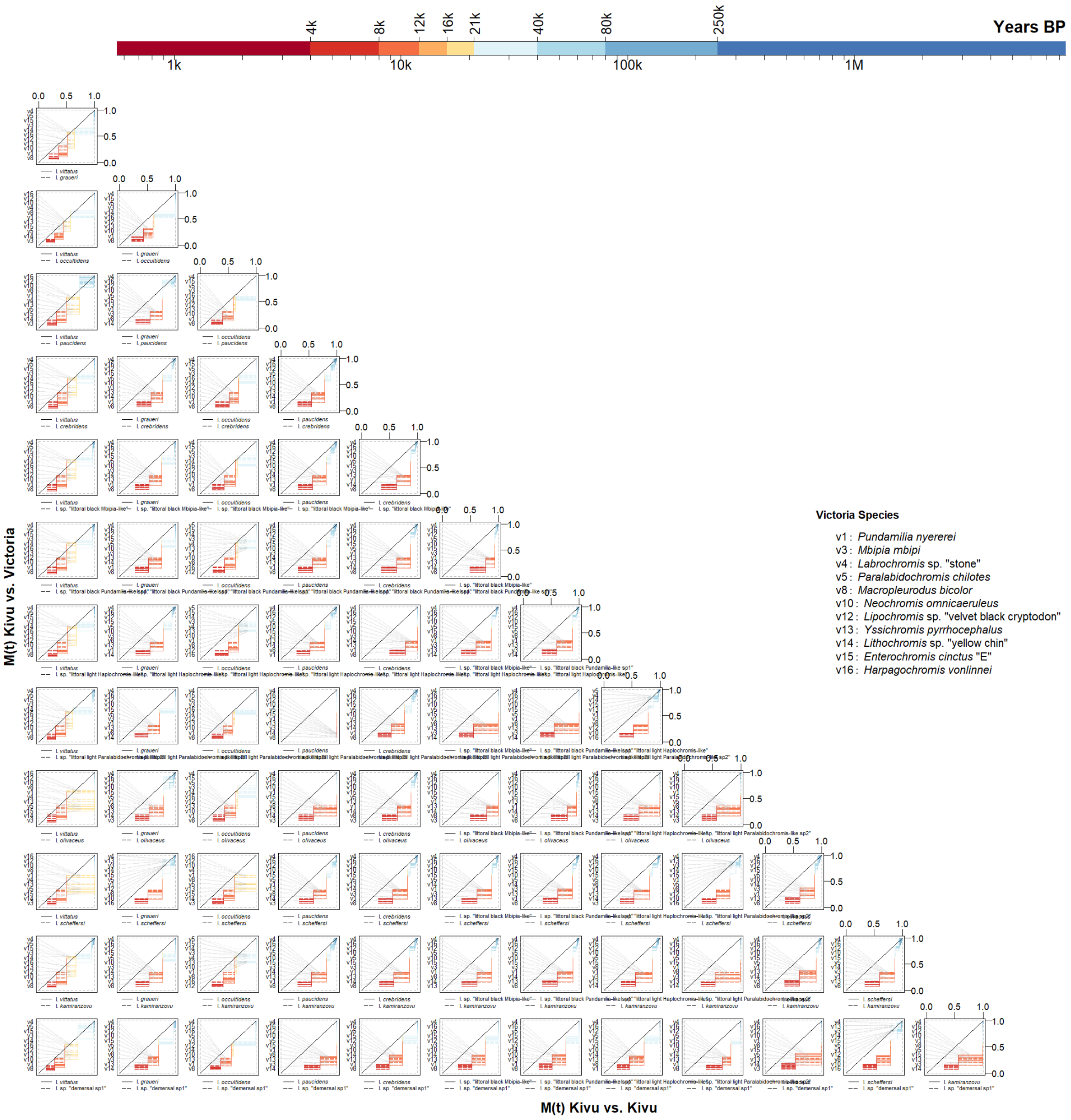

**Supplementary Fig. 15 | Observed lineage merging among Lake Kivu cichlids from the same lake vs. from Lake Victoria, resolved by species.** Cumulative migration probability M(t) trajectories reconstructed from two haplotypes per species are shown for species pairs from within and between lakes. The color scale gives the timing of lineage merging.

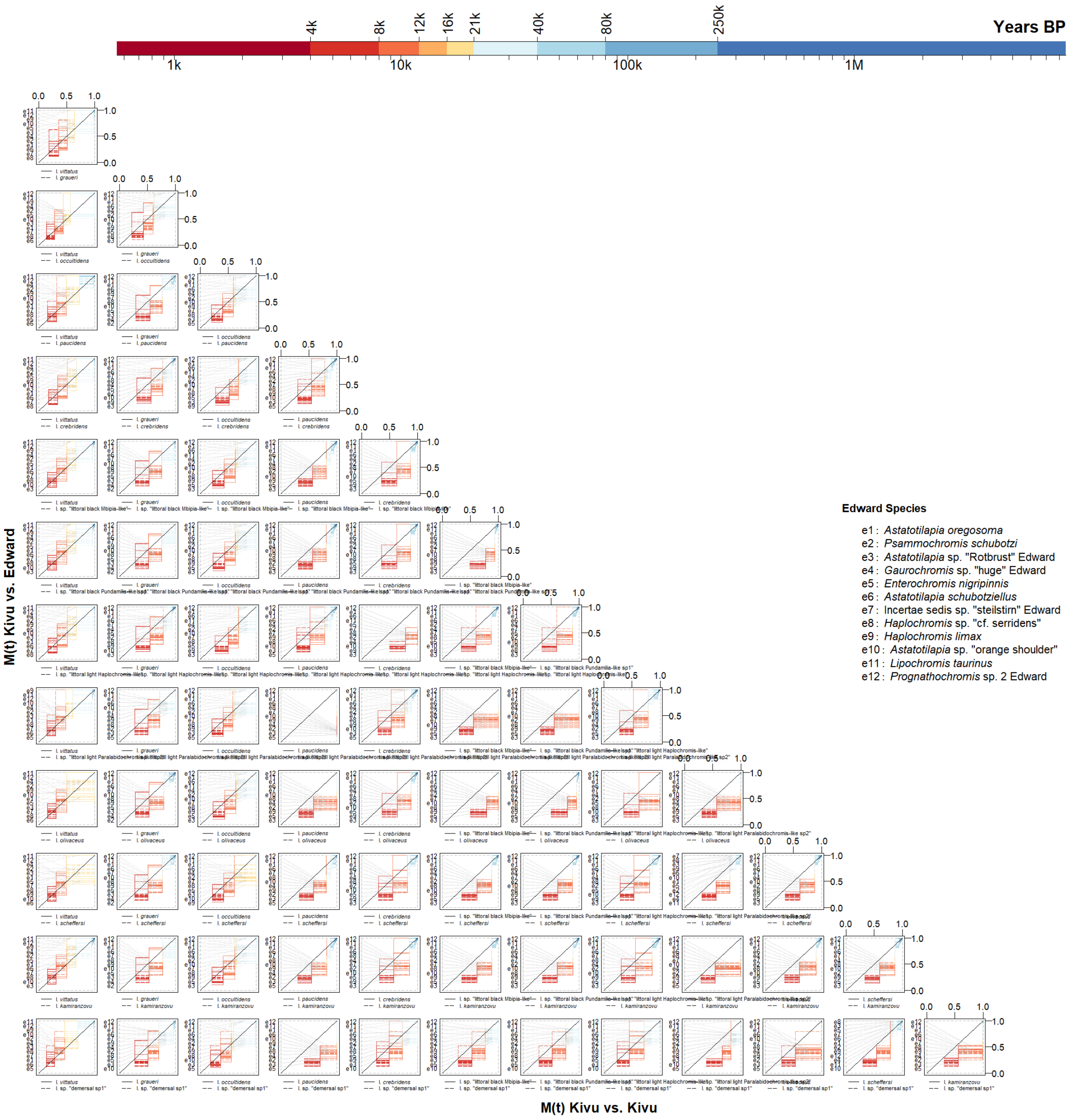

**Supplementary Fig. 16 | Observed lineage merging among Lake Kivu cichlids from the same lake vs. from Lake Edward, resolved by species.** Cumulative migration probability M(t) trajectories reconstructed from two haplotypes per species are shown for species pairs from within and between lakes. The color scale gives the timing of lineage merging.

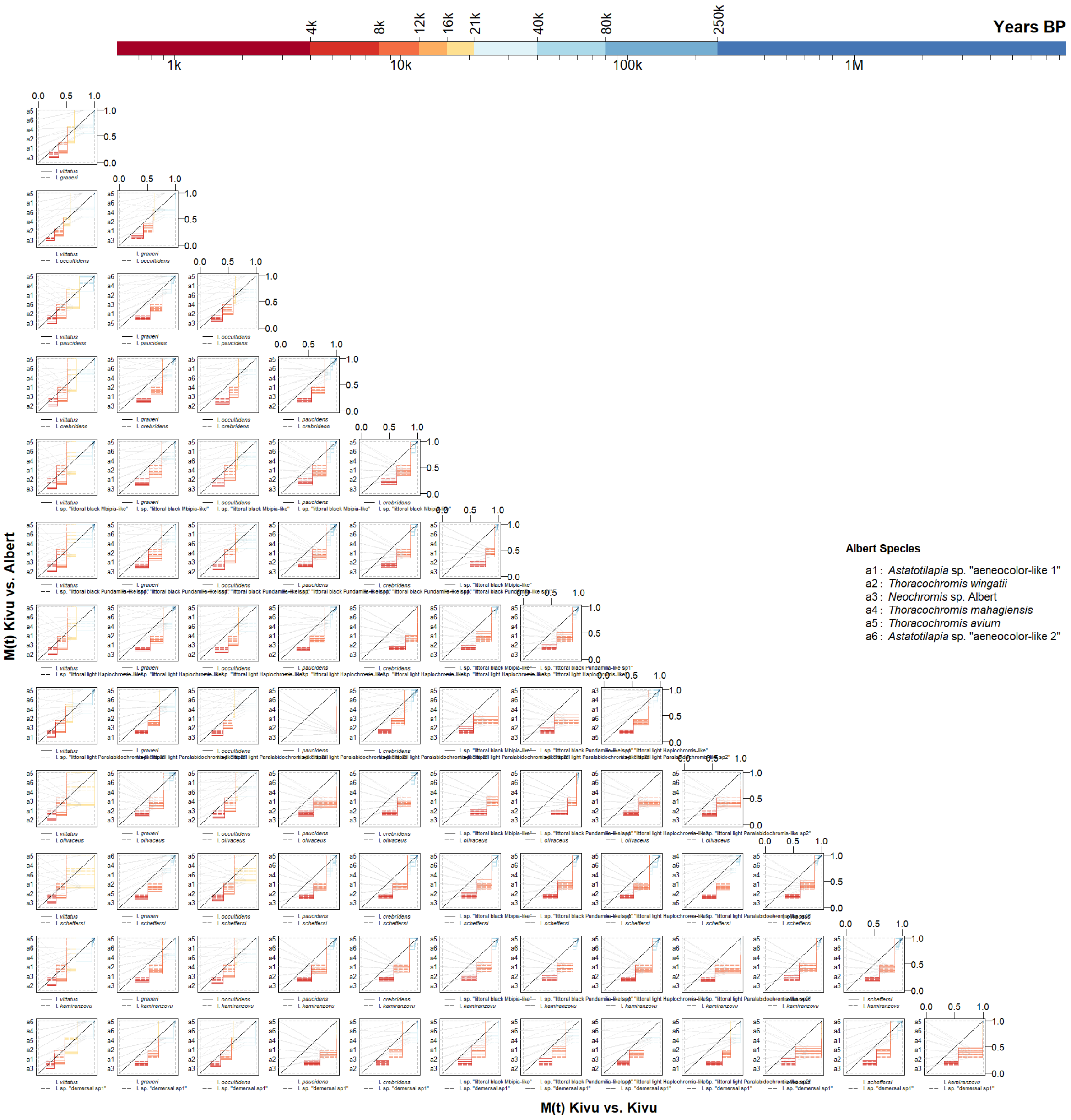

**Supplementary Fig. 17 | Observed lineage merging among Lake Kivu cichlids from the same lake vs. from Lake Albert, resolved by species.** Cumulative migration probability M(t) trajectories reconstructed from two haplotypes per species are shown for species pairs from within and between lakes. The color scale gives the timing of lineage merging.

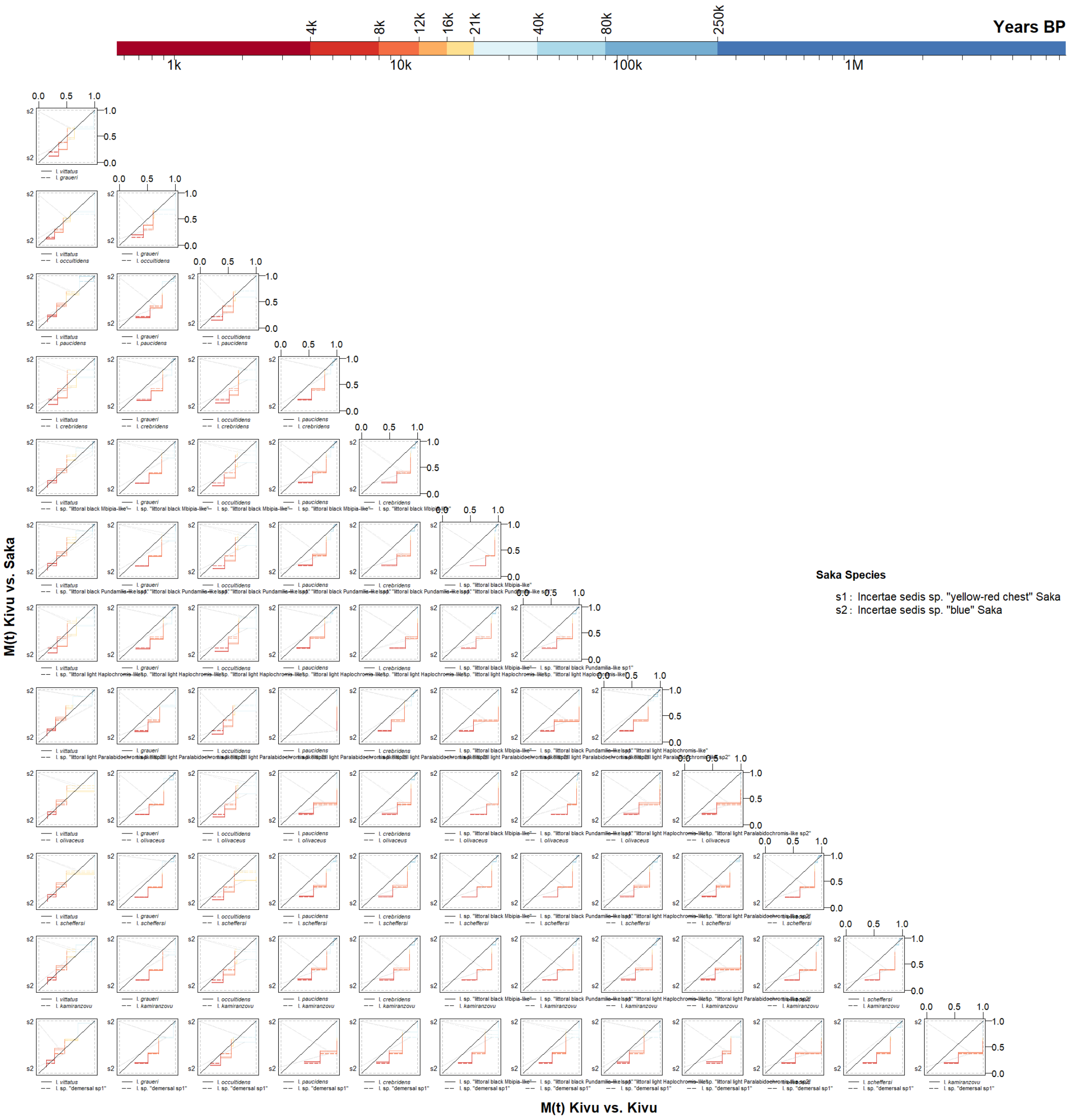

**Supplementary Fig. 18 | Observed lineage merging among Lake Kivu cichlids from the same lake vs. from Lake Saka, resolved by species.** Cumulative migration probability M(t) trajectories reconstructed from two haplotypes per species are shown for species pairs from within and between lakes. The color scale gives the timing of lineage merging.

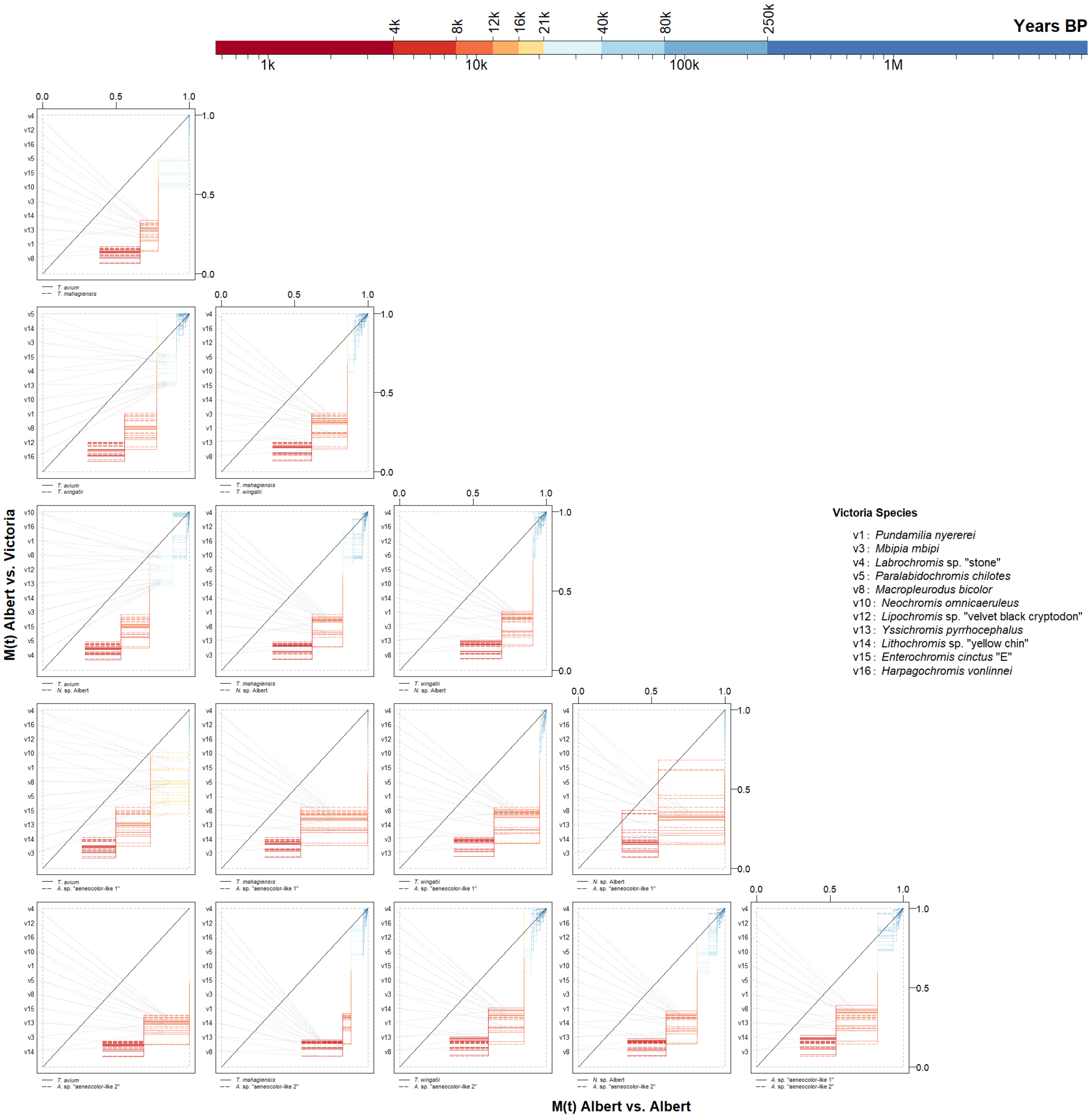

**Supplementary Fig. 19 | Observed lineage merging among Lake Albert cichlids from the same lake vs. from Lake Victoria, resolved by species.** Cumulative migration probability M(t) trajectories reconstructed from two haplotypes per species are shown for species pairs from within and between lakes. The color scale gives the timing of lineage merging.

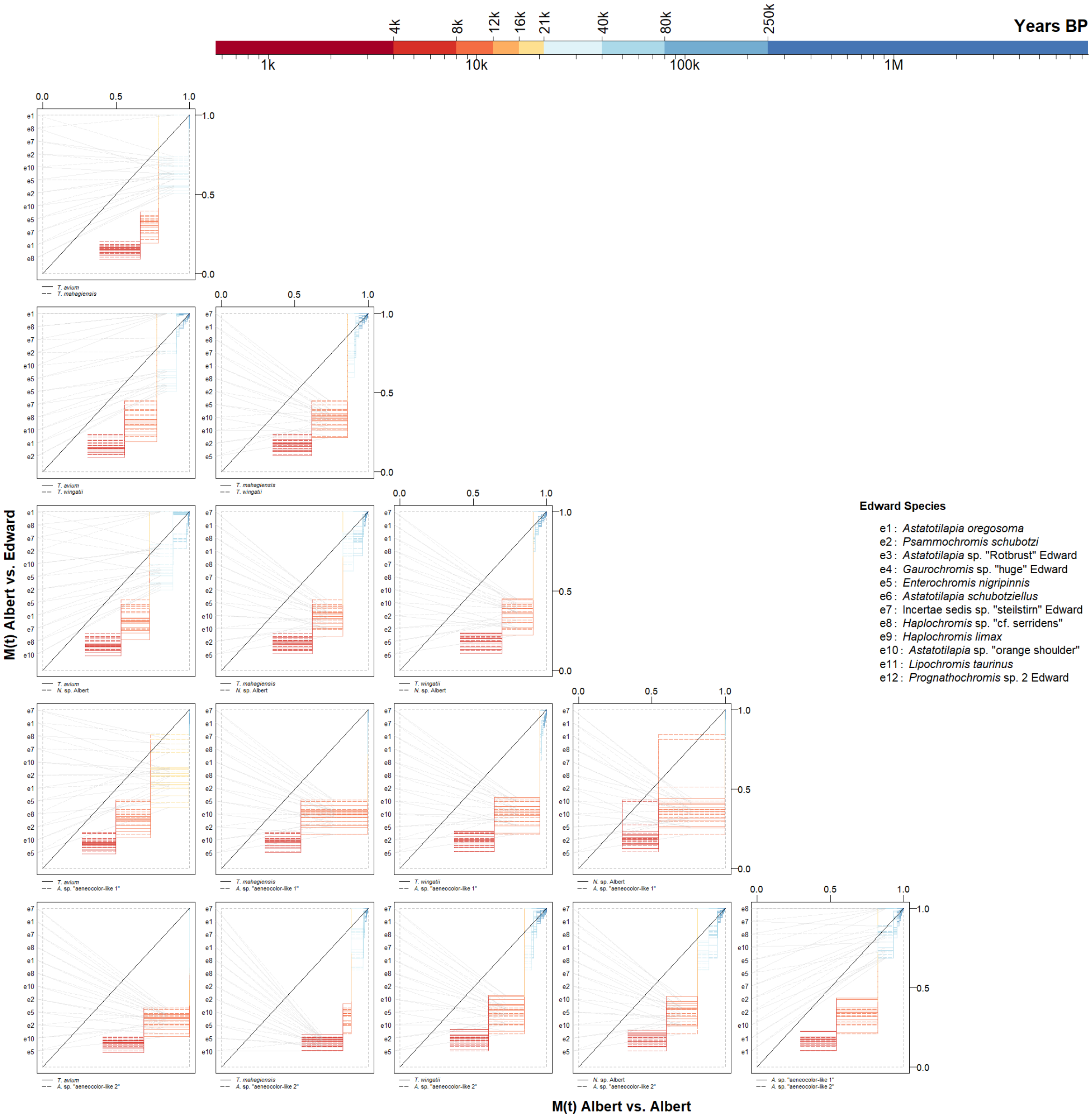

**Supplementary Fig. 20 | Observed lineage merging among Lake Albert cichlids from the same lake vs. from Lake Edward, resolved by species.** Cumulative migration probability M(t) trajectories reconstructed from two haplotypes per species are shown for species pairs from within and between lakes. The color scale gives the timing of lineage merging.

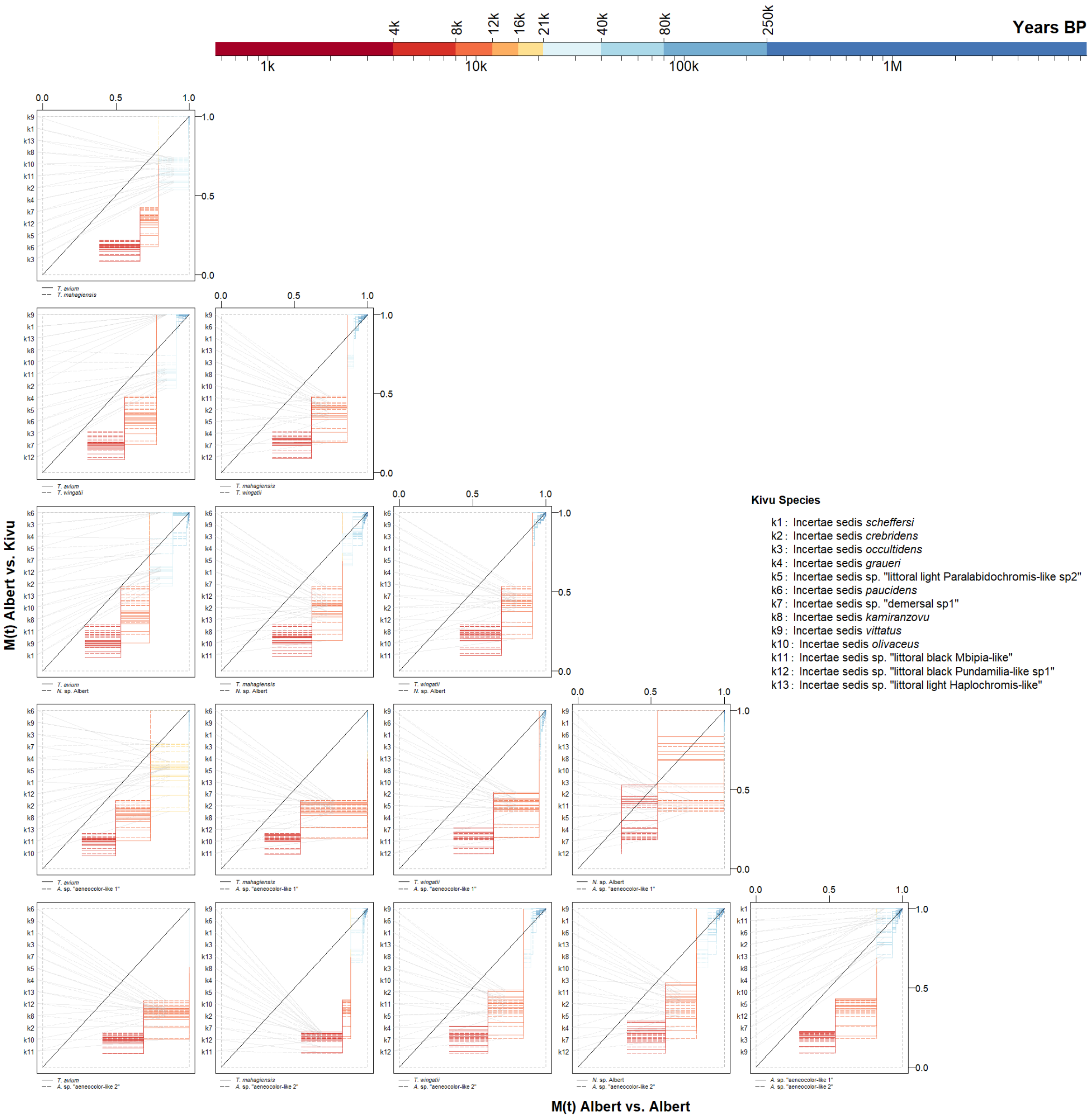

**Supplementary Fig. 21 | Observed lineage merging among Lake Albert cichlids from the same lake vs. from Lake Kivu, resolved by species.** Cumulative migration probability M(t) trajectories reconstructed from two haplotypes per species are shown for species pairs from within and between lakes. The color scale gives the timing of lineage merging.

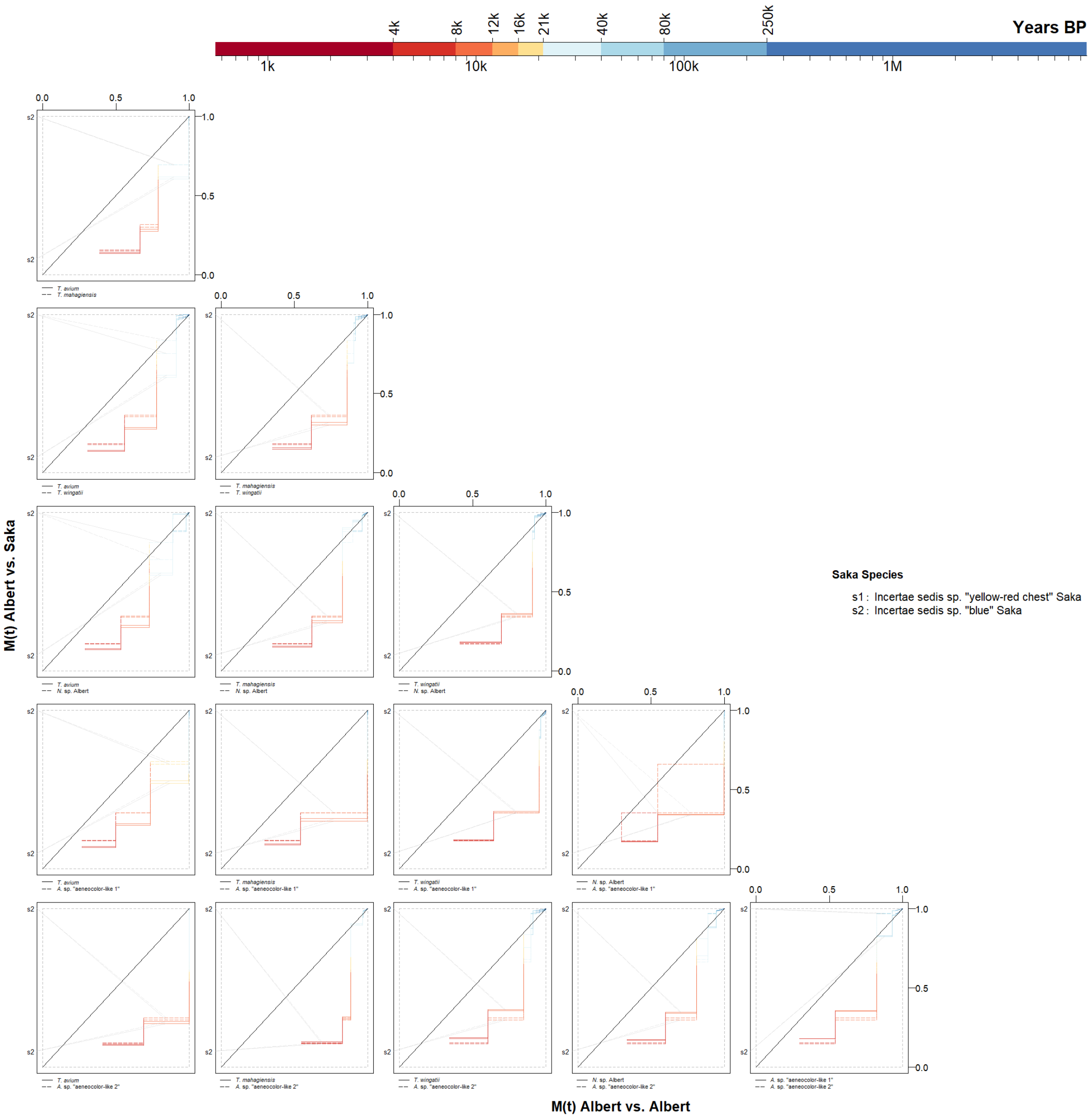

**Supplementary Fig. 22 | Observed lineage merging among Lake Albert cichlids from the same lake vs. from Lake Saka, resolved by species.** Cumulative migration probability M(t) trajectories reconstructed from two haplotypes per species are shown for species pairs from within and between lakes. The color scale gives the timing of lineage merging.

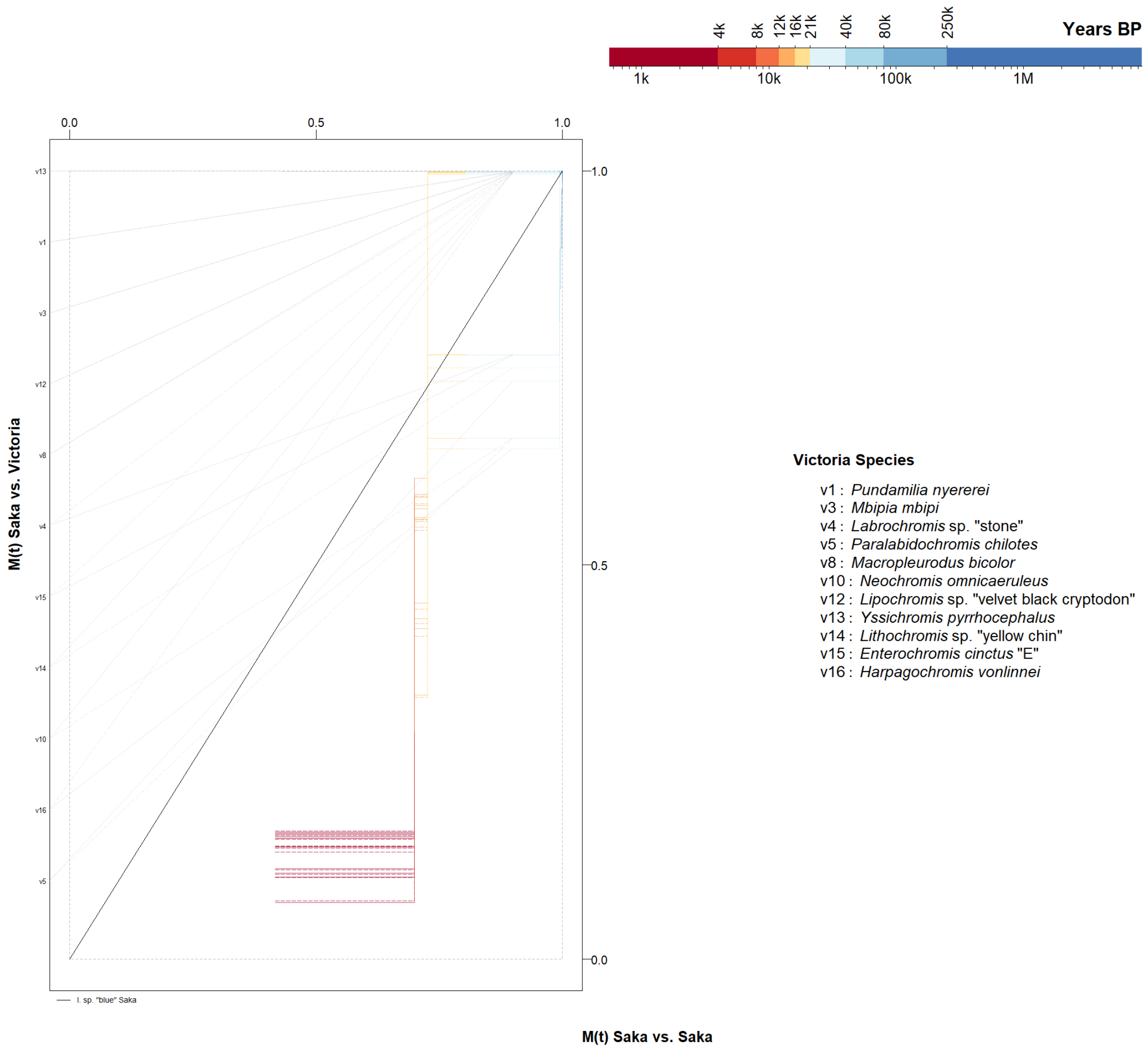

**Supplementary Fig. 23 | Observed lineage merging among Lake Saka cichlids from the same lake vs. from Lake Victoria, resolved by species.** Cumulative migration probability M(t) trajectories reconstructed from two haplotypes per species are shown for species pairs from within and between lakes. The color scale gives the timing of lineage merging.

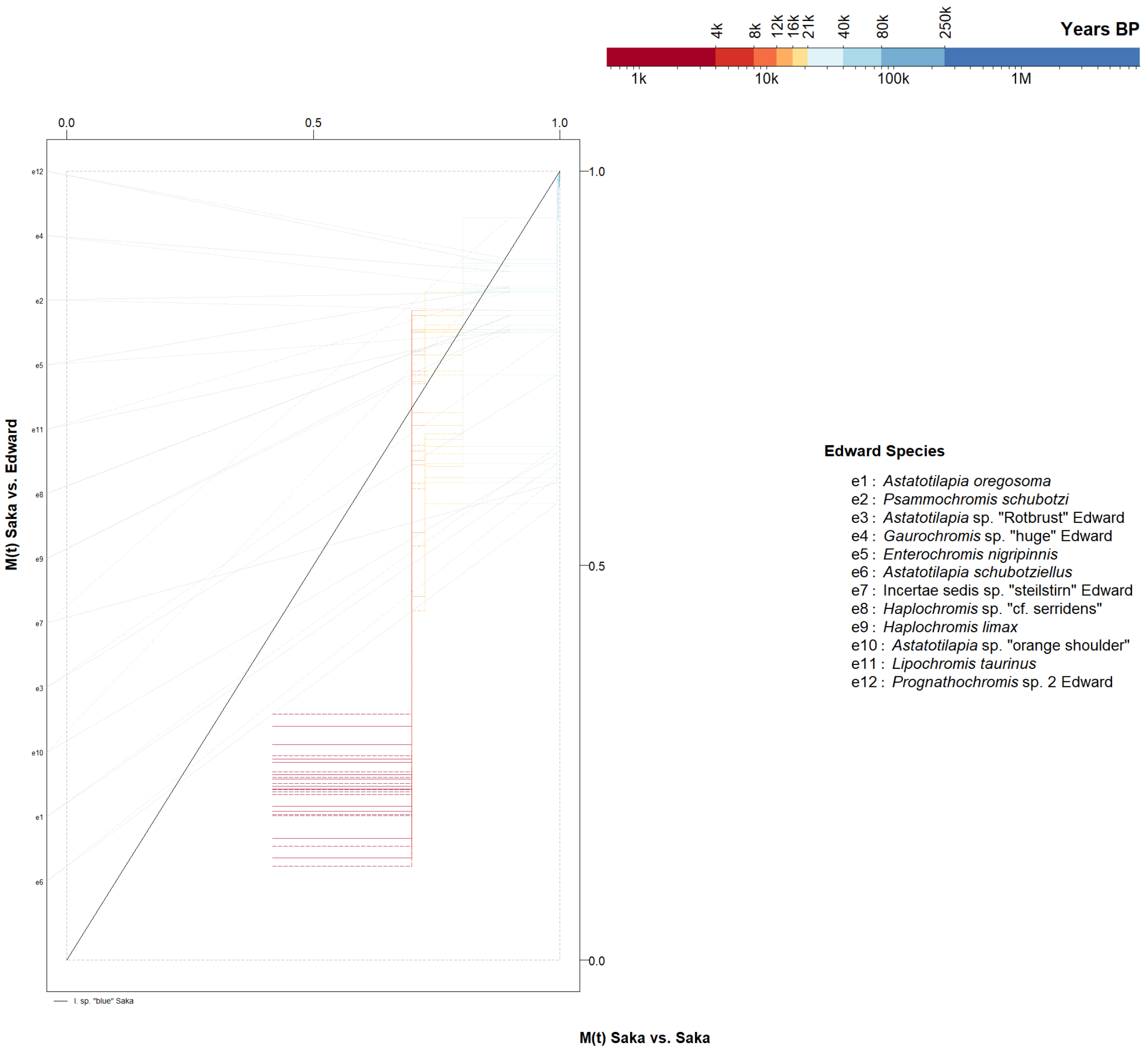

**Supplementary Fig. 24 | Observed lineage merging among Lake Saka cichlids from the same lake vs. from Lake Edward, resolved by species.** Cumulative migration probability M(t) trajectories reconstructed from two haplotypes per species are shown for species pairs from within and between lakes. The color scale gives the timing of lineage merging.

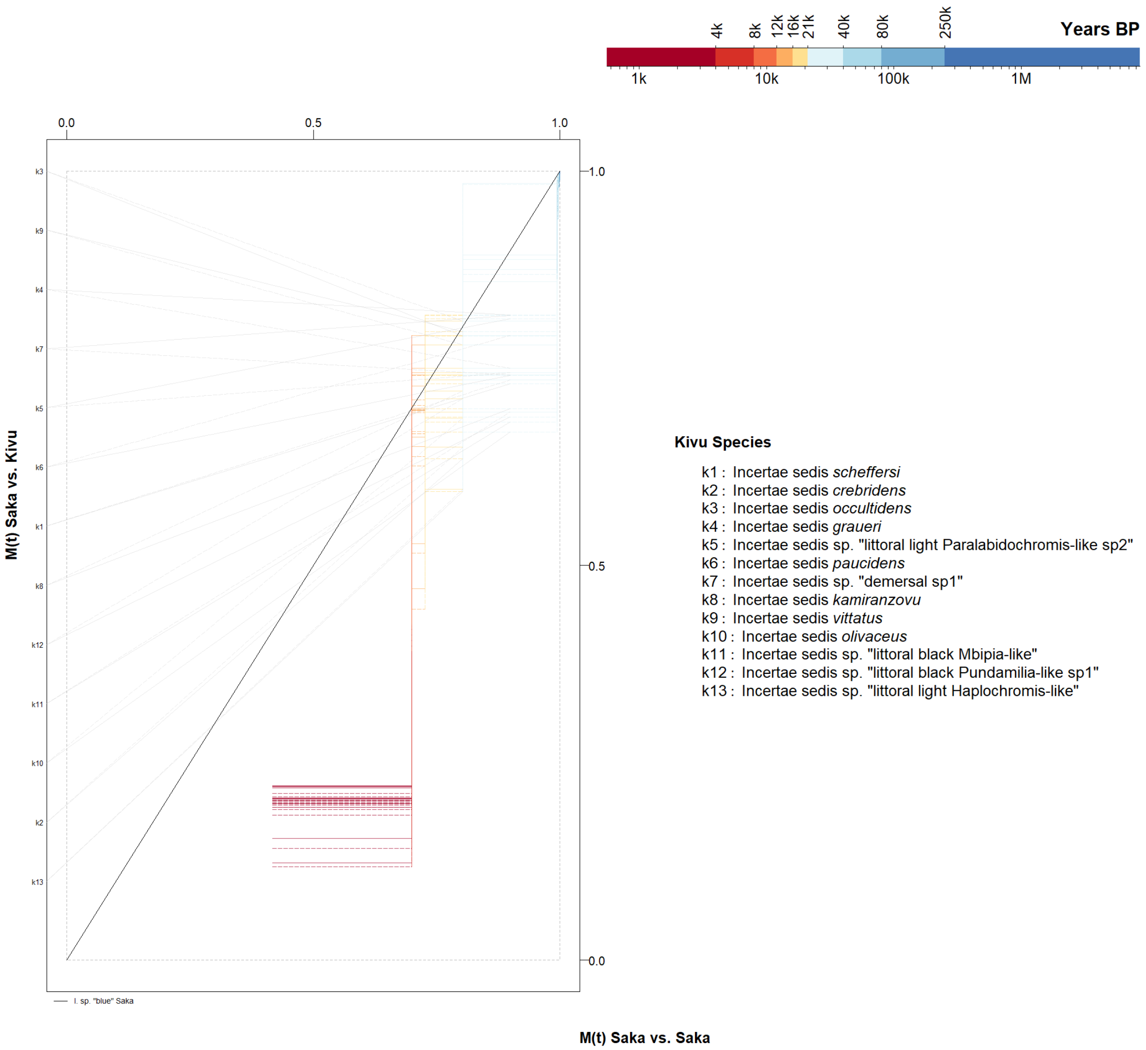

**Supplementary Fig. 25 | Observed lineage merging among Lake Saka cichlids from the same lake vs. from Lake Kivu, resolved by species.** Cumulative migration probability M(t) trajectories reconstructed from two haplotypes per species are shown for species pairs from within and between lakes. The color scale gives the timing of lineage merging.

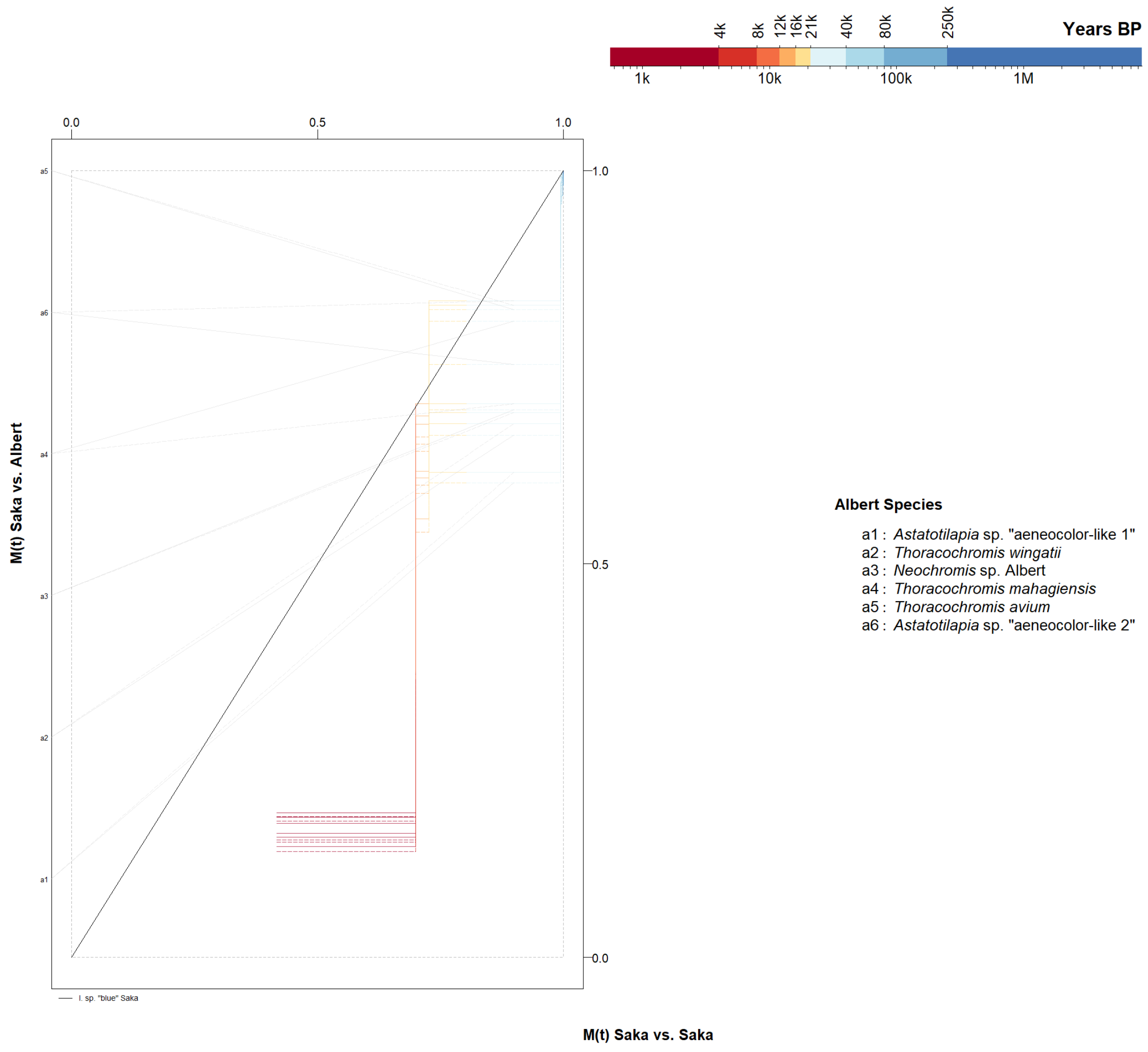

**Supplementary Fig. 26 | Observed lineage merging among Lake Saka cichlids from the same lake vs. from Lake Albert, resolved by species.** Cumulative migration probability M(t) trajectories reconstructed from two haplotypes per species are shown for species pairs from within and between lakes. The color scale gives the timing of lineage merging.

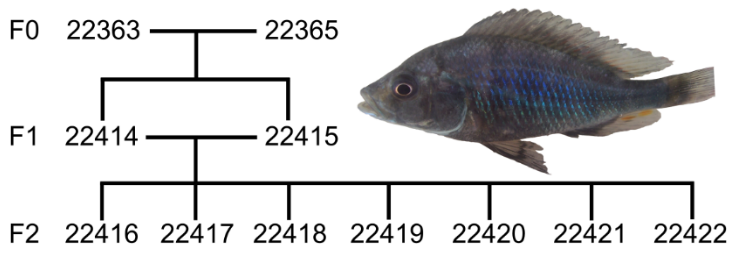

**Supplementary Fig. 27 | Experimental setup for mutation rate estimation in a Lake Victoria haplochromine cichlid.** Pedigree of the 11 member *Gaurochromis hiatus* family used to estimate the Lake Victoria haplochromine cichlid mutation rate, with numbers indicating individual identifiers (Supplementary Table 1). F0, F1 and F2 are the respective grandparent, parent and offspring generations. The picture shows a male displaying nuptial coloration of this species.

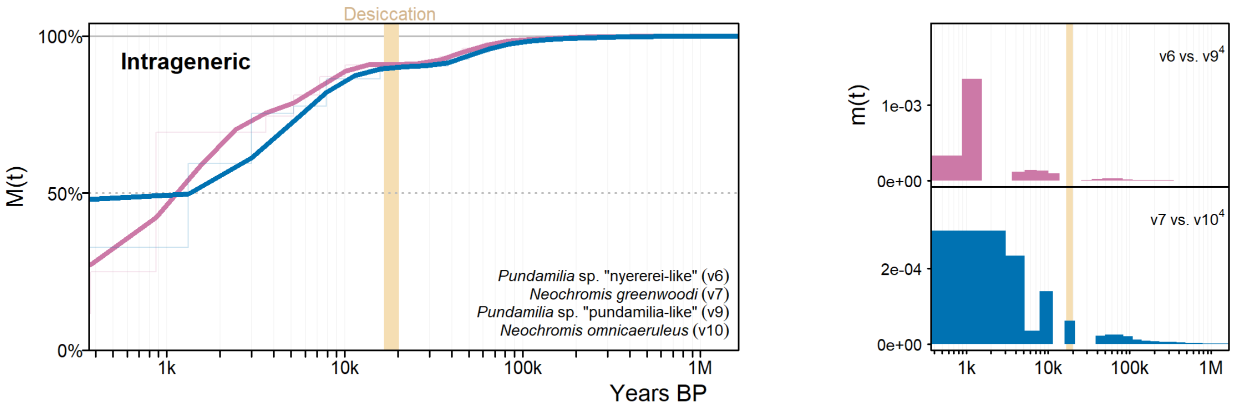

**Supplementary Fig. 28 | Observed lineage merging between closely related Victoria cichlids from the same genus.** The two species pairs, members of the same genus and the Victoria radiation, merge very recently into a common ancestor, much more recently than different guilds (Supplementary Fig. 1), confirming the recency of two speciation events. Cumulative migration probability M(t) and migration rates m(t) between species were estimated from six (*Neochromis*) and eight (*Pundamilia*) haplotypes per species, with the beige vertical bar denoting the period in which Lake Victoria dried out between 16,700-20,200 years ago. The reconstruction is at the lower inference limit of the MSMC-IM method.

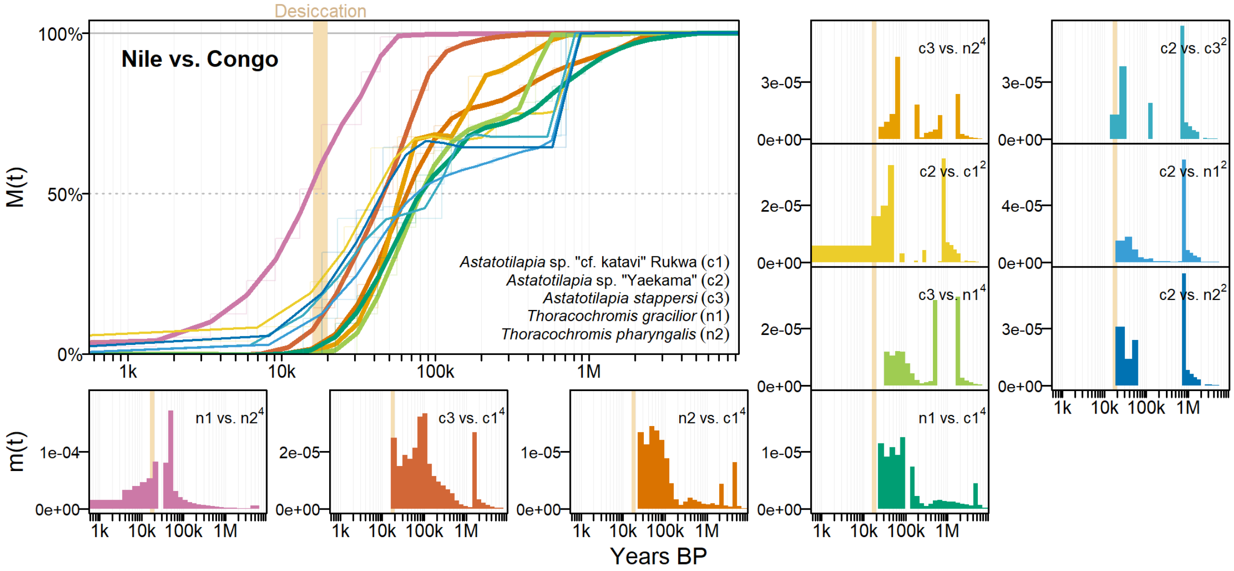

**Supplementary Fig. 29 | Observed lineage merging between haplochromine cichlids of the Congo and Upper Nile sister lineages to the LVRS cichlids.** The two representatives of the Congo and Upper Nile clades each merge around 40,000 and 100,000 years ago into a common ancestor, while between Upper Nile and Congo lineages, coalescence falls into two or more discrete migration rate m(t) episodes with considerable plateaus in cumulative migration probability M(t). The latter suggests considerable recent admixture between Congo and Upper Nile lineages, during the formation of the LVRS hybrid lineage, after an ancient origin of these two lineages 1-6 MY ago, consistent with phylogenetic divergence estimates for this group. The beige vertical bar denotes the period in which Lake Victoria dried out 16,700-20,200 years ago.

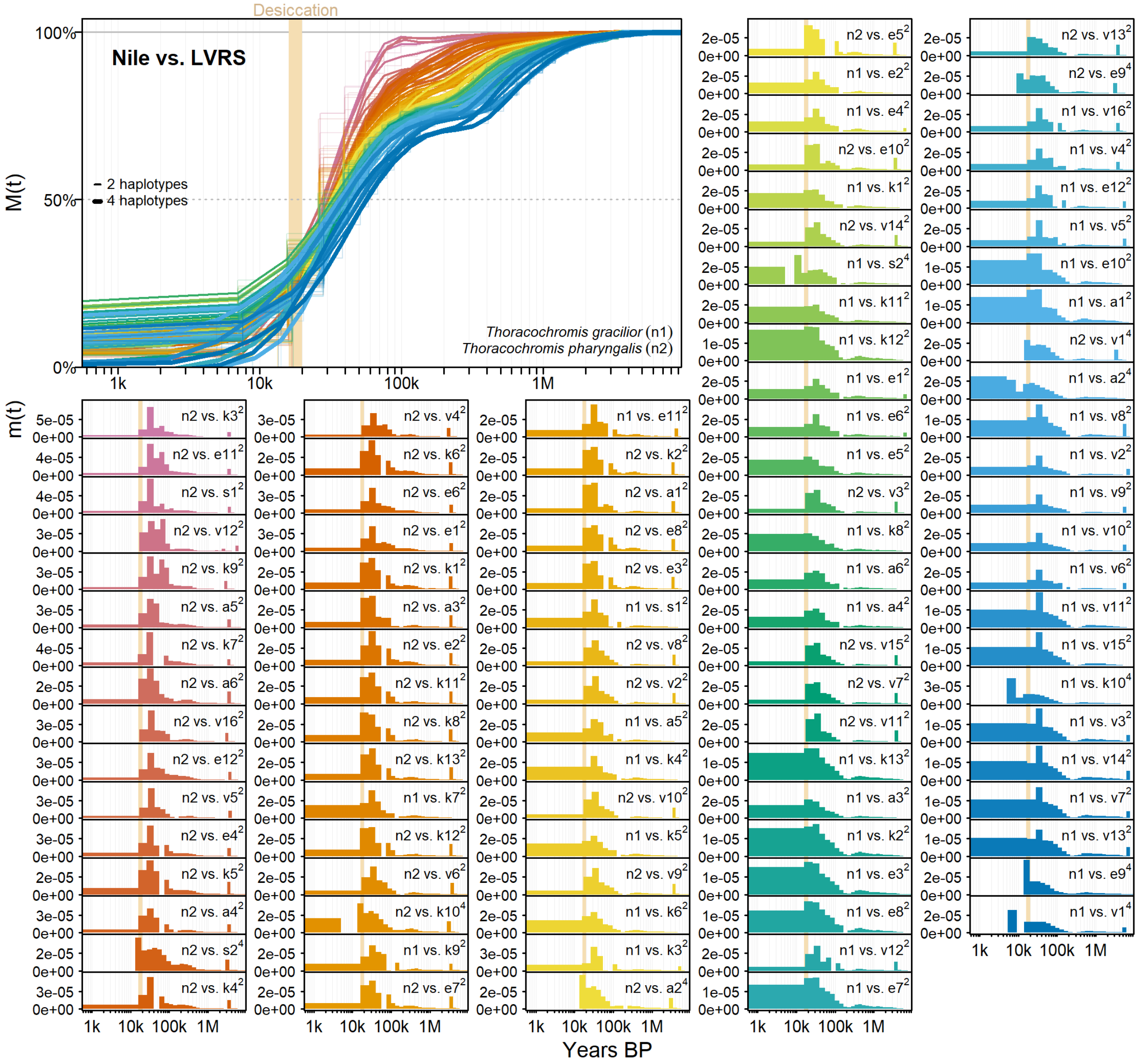

**Supplementary Fig. 30 | Observed lineage merging between haplochromine cichlids of the Nile sister lineages and members of the LVRS hybrid lineage.** The LVRS and Nile cichlids lineages merge most rapidly around 30,000-80,000 years ago, consistent with the formation of the LVRS hybrid swarm in this time range. Note the plateau in M(t) between this time range and complete coalescence only around 2 My ago, the likely split time between the contributing Congo and Nile lineages (Supplementary Note 4). Species numbers are explained in Fig. 1, Supplementary Figs. 1-5 and in Supplementary Table 1. Line widths and superscripts behind species numbers indicate whether two or four haplotypes per species were used to estimate cumulative migration probability M(t) and migration rates m(t). The beige vertical bare denotes the arid period 16,700-20,200 years ago that led to the desiccation of Lake Victoria.

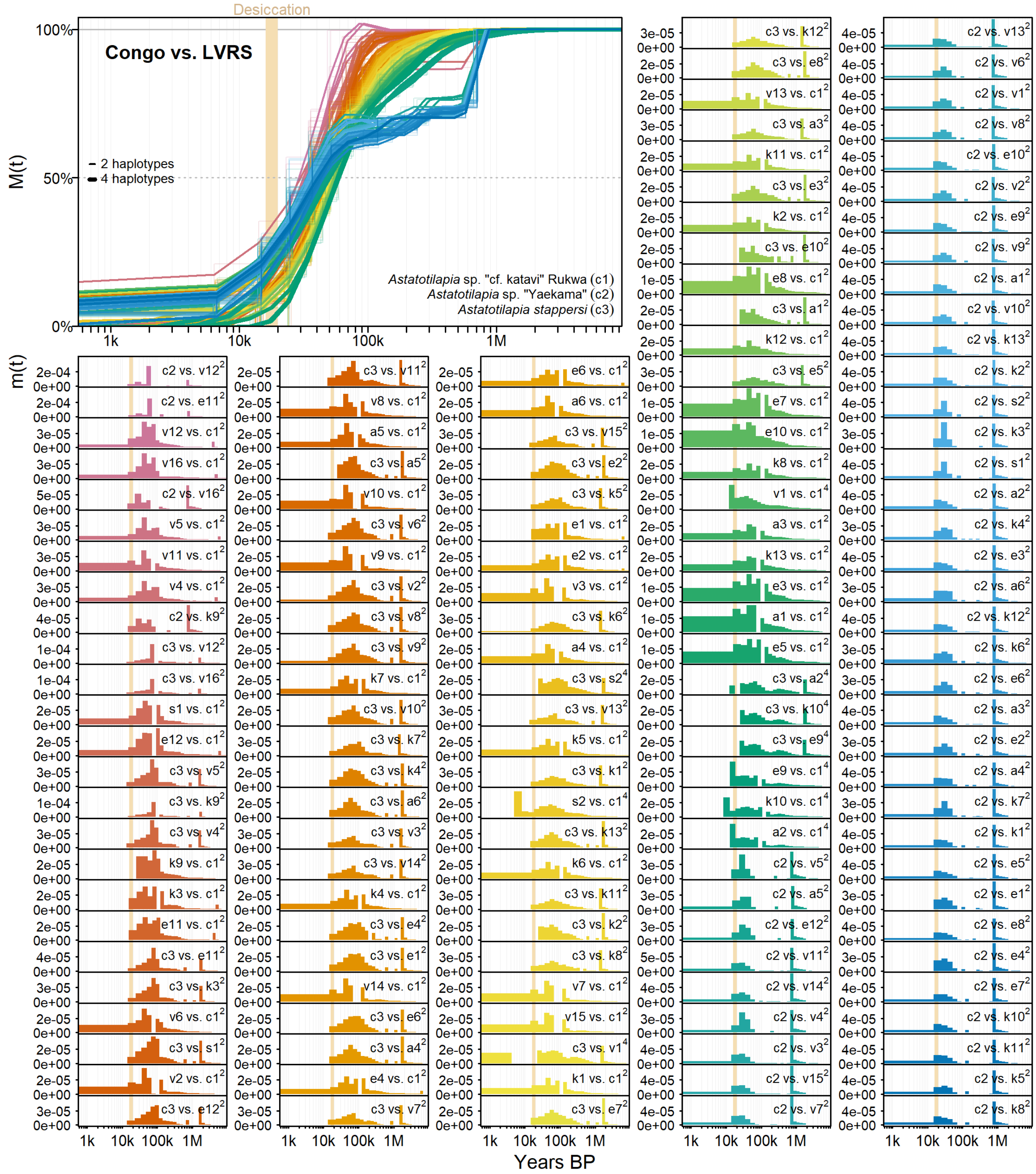

**Supplementary Fig. 31 | Observed lineage merging between haplochromine cichlids of the Congo sister lineage and members of the LVRS hybrid lineage.** LVRS and Congo cichlid lineages merged most rapidly around 30,000-80,000 years ago, consistent with the formation of the LVRS hybrid swarm in this time range. Note the plateau in M(t) between this time range and complete coalescence only around 2 My ago, the likely split time between the contributing Congo and Nile lineages (Supplementary Note 4). Species numbers are explained in Fig. 1, Supplementary Figs. 1-5 and in Supplementary Table 1. Line widths and superscripts behind species numbers indicate whether two or four haplotypes per species were used to estimate cumulative migration probability M(t) and migration rates m(t). The beige vertical bare denotes the arid period 16,700-20,200 years ago that led to the desiccation of Lake Victoria.

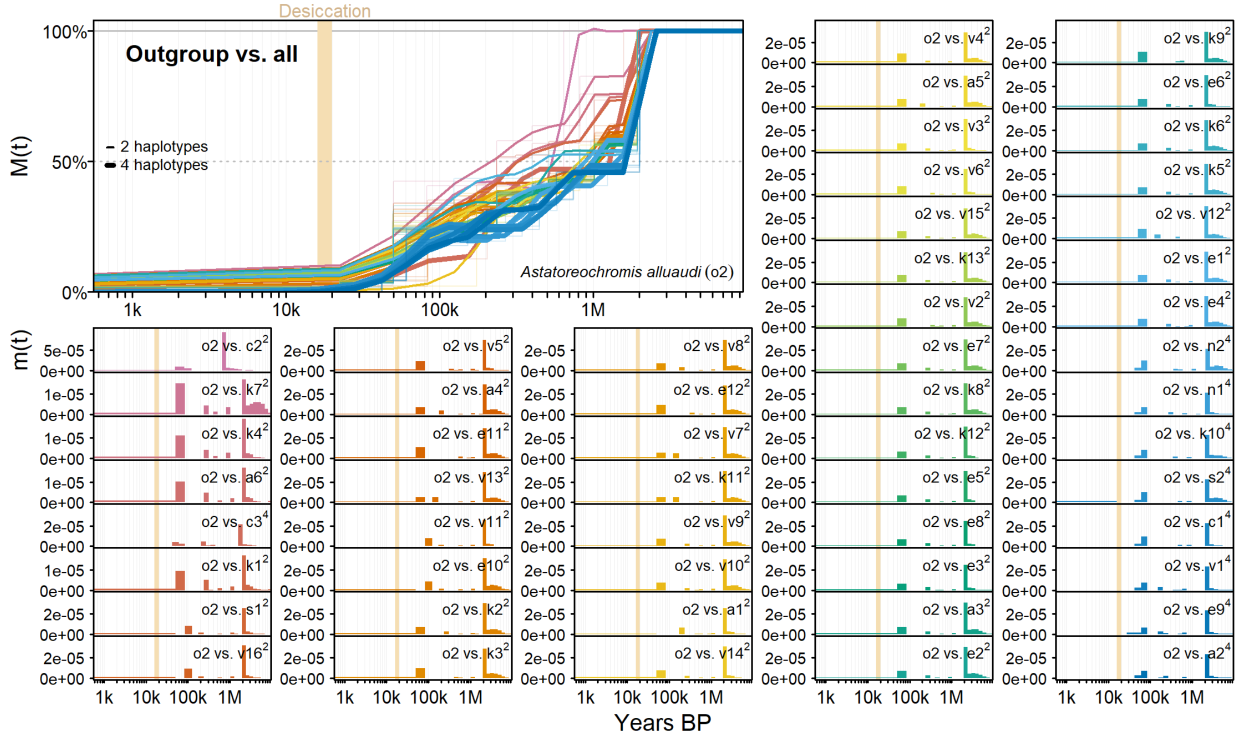

**Supplementary Fig. 32 | Lineage merging between the outgroup species *Astatoreochromis alluaudi* and LVRS, Congo and Nile haplochromine cichlids.** All haplochromine cichlids of the LVRS, Congo and Nile lineages merge simultaneously with a distant outgroup species, with minor migration around 50,000-80,000 years and major migration 2-6 MY ago, suggesting ancient divergence from the outgroup but surprisingly some minor recent gene flow. Species numbers are explained in Fig. 1, Supplementary Figs. 1-5 and in Supplementary Table 1. Line widths and superscripts behind species numbers indicate whether two or four haplotypes per species were used to estimate cumulative migration probability M(t) and migration rates m(t). The beige vertical bare denotes the arid period 16,700-20,200 years ago that led to the desiccation of Lake Victoria.

415 **Supplementary Tables**

416 **Supplementary Table 1 |** List of sequenced individuals, origin, sequencing effort, depth and analysis involvement. FishecID specifies unique identifiers used in
417 the Seehausen lab. Columns 2hap, 4hap, 6hap and 8hap indicate in which combinations individuals were used in MSMC2 / MSMC-IM analyses.

| FishecID | Species | sID | Sex | Year | Collectors | Location | Country | Lake | Group | Trophic Level | Trophic Group | Habitat | Number of Reads | Mean Depth | Accession | 2hap | 4hap | 6hap | 8hap |
|---|---|---|---|---|---|---|---|---|---|---|---|---|---|---|---|---|---|---|---|
| 71004 | *Astatotilapia* sp. "cf. katavi" Rukwa | c1 | U | 2018 | M. Kishe, TAFIRI | Rukwa | Tanzania | Rukwa | Congolese lineage | 3 | insectivore | littoral | 127,960,848 | 31.0 | SAMN32927018 | | x | | |
| 71006 | *Astatotilapia* sp. "cf. katavi" Rukwa | c1 | U | 2018 | M. Kishe, TAFIRI | Rukwa | Tanzania | Rukwa | Congolese lineage | 3 | insectivore | littoral | 133,618,788 | 32.1 | SAMN32927020 | x | x | | |
| 81309 | *Astatotilapia* sp. "Yaekama" | c2 | M | 2009 | U. Schliewen | Yaekama | DR Congo | Congo | Congolese lineage | 3 | insectivore | streams | 94,650,423 | 14.4 | SAMN32927054 | x | | | |
| 81343 | *Astatotilapia stappersi* | c3 | U | | W. Salzburger | Kalambo River | Zambia | Congo | Congolese lineage | 3 | insectivore | streams | 53,300,342 | 12.5 | SAMN32927055 | x | x | | |
| 81345 | *Astatotilapia stappersi* | c3 | U | | W. Salzburger | Kalambo River | Zambia | Congo | Congolese lineage | 3 | insectivore | streams | 50,717,112 | 10.5 | SAMN32927057 | | x | | |
| 80942 | *Astatotilapia* sp. "aeneocolor-like 1" | a1 | M | 2002 | S .B. Wandera | Albert | Uganda | Albert | LVRS Albert | 3 | insectivore | littoral | 143,239,396 | 35.2 | SAMN32927032 | x | x | | |
| 80955 | *Astatotilapia* sp. "aeneocolor-like 1" | a1 | M | 2002 | S .B. Wandera | Albert | Uganda | Albert | LVRS Albert | 3 | insectivore | littoral | 95,764,225 | 23.3 | SAMN32927033 | | x | | |
| 80978 | *Astatotilapia* sp. "aeneocolor-like 2" | a6 | M | 2002 | S .B. Wandera | Albert | Uganda | Albert | LVRS Albert | 3 | insectivore | littoral | 91,346,392 | 22.3 | SAMN32927037 | x | | | |
| 80984 | *Neochromis* sp. Albert | a3 | U | 2002 | S .B. Wandera | Albert | Uganda | Albert | LVRS Albert | 2 | epilithic algae browser | rocky | 123,322,941 | 30.3 | SAMN32927041 | x | | | |
| 80977 | *Thoracochromis avium* | a5 | M | 2002 | S .B. Wandera | Albert | Uganda | Albert | LVRS Albert | 4 | piscivore | littoral | 88,093,784 | 21.5 | SAMN32927036 | x | | | |
| 80980 | *Thoracochromis mahagiensis* | a4 | M | 2002 | S .B. Wandera | Albert | Uganda | Albert | LVRS Albert | 3 | snail crusher | littoral | 72,071,801 | 17.7 | SAMN32927039 | | x | | |
| 80981 | *Thoracochromis mahagiensis* | a4 | M | 2002 | S .B. Wandera | Albert | Uganda | Albert | LVRS Albert | 3 | snail crusher | littoral | 72,032,332 | 17.8 | SAMN32927040 | x | x | | |
| 80959 | *Thoracochromis wingatii* | a2 | U | 2002 | S .B. Wandera | Albert | Uganda | Albert | LVRS Albert | 3 | insectivore | littoral | 75,433,240 | 18.5 | SAMN32927034 | | x | | |
| 80964 | *Thoracochromis wingatii* | a2 | U | 2002 | S .B. Wandera | Albert | Uganda | Albert | LVRS Albert | 3 | insectivore | littoral | 95,963,065 | 23.6 | SAMN32927035 | x | x | | |
| 80994 | *Astatotilapia oregosoma* | e1 | U | 2000 | O. Seehausen, L. J. Chapman, C. A. Chapman | Edward | Uganda | Edward | LVRS Edward | unknown | unknown | littoral | 57,218,616 | 14.9 | SAMN32927042 | | x | | |
| 80996 | *Astatotilapia oregosoma* | e1 | U | 2000 | O. Seehausen, L. J. Chapman, C. A. Chapman | Edward | Uganda | Edward | LVRS Edward | unknown | unknown | littoral | 42,770,426 | 15.6 | SAMN32927043 | x | x | | |
| 81095 | *Astatotilapia schubotziellus* | e6 | F | 2003 | O. Seehausen | Edward | Uganda | Edward | LVRS Edward | 3 | insectivore | littoral | 71,034,667 | 16.8 | SAMN32927051 | x | | | |
| 81000 | *Astatotilapia* sp. "orange shoulder" | e10 | U | 2000 | O. Seehausen, L. J. Chapman, C. A. Chapman | Edward | Uganda | Edward | LVRS Edward | 3 | insectivore | littoral | 50,521,905 | 24.8 | SAMN32927044 | x | | | |
| 81077 | *Enterochromis nigripinnis* | e5 | M | 2003 | O. Seehausen | Edward | Uganda | Edward | LVRS Edward | 2 | phytoplankton | demersal | 56,492,184 | 14.7 | SAMN32927049 | x | | | |
| 81084 | *Gaurochromis* sp. "huge" Edward | e4 | M | 2003 | O. Seehausen | Edward | Uganda | Edward | LVRS Edward | 4 | insectivore & piscivore | demersal | 58,665,922 | 14.0 | SAMN47214501 | x | | | |
| 81103 | *Haplochromis limax* | e9 | U | 2003 | O. Seehausen | Edward | Uganda | Edward | LVRS Edward | 2 | epiphythic algae grazer | littoral | 29,496,806 | 29.8 | SAMN32927052 | x | x | | |
| 71015 | *Haplochromis limax* | e9 | U | 2003 | O. Seehausen | Edward | Uganda | Edward | LVRS Edward | 2 | epiphythic algae grazer | littoral | 63,832,973 | 16.1 | SAMN32927028 | | x | | |
| 81709 | *Haplochromis* sp. "cf. serridens" | e8 | U | | E. Schraml | Edward | Uganda | Edward | LVRS Edward | unknown | unknown | littoral | 62,576,932 | 19.4 | SAMN32927061 | x | | | |
| 81710 | Incertae sedis sp. "steilstirn" Edward | e7 | U | | E. Schraml | Edward | Uganda | Edward | LVRS Edward | 2 | epiphythic algae grazer | littoral | 46,063,871 | 21.8 | SAMN32927062 | x | | | |
| 81023 | *Lipochromis taurinus* | e11 | U | 2000 | O. Seehausen, L. J. Chapman, C. A. Chapman | Edward | Uganda | Edward | LVRS Edward | 4 | paedophage | littoral | 63,828,783 | 17.3 | SAMN32927046 | x | | | |
| 81027 | *Prognathochromis* sp. 2 Edward | e12 | U | 2000 | O. Seehausen, L. J. Chapman, C. A. Chapman | Edward | Uganda | Edward | LVRS Edward | 4 | piscivore | demersal | 67,655,137 | 16.1 | SAMN47214502 | x | | | |
| 81073 | *Psammochromis schubotzi* | e2 | M | 2003 | O. Seehausen | Edward | Uganda | Edward | LVRS Edward | 3 | insectivore | demersal | 97,714,216 | 22.0 | SAMN32927048 | x | | | |
| 81706 | *Ptyochromis* sp. "Rotbrust" Edward | e3 | U | | E. Schraml | Edward | Uganda | Edward | LVRS Edward | unknown | unknown | littoral | 54,996,812 | 19.6 | SAMN32927059 | x | | | |
| 64530 | Incertae sedis *scheffersi* | k1 | M | 2012 | G. Periat, O. Seehausen | Kivu | Rwanda | Kivu | LVRS Kivu | 3 | insectivore | unknown | 78,258,692 | 18.6 | SAMN32926981 | x | x | | |
| 64531 | Incertae sedis *scheffersi* | k1 | M | 2012 | G. Periat, O. Seehausen | Kivu | Rwanda | Kivu | LVRS Kivu | 3 | insectivore | unknown | 78,618,455 | 19.0 | SAMN32926982 | | x | | |
| 64236 | Incertae sedis sp. "demersal sp1" | k7 | M | 2012 | G. Periat, O. Seehausen | Kivu | Rwanda | Kivu | LVRS Kivu | unknown | unknown | demersal | 86,512,762 | 21.0 | SAMN32926959 | | x | | |
| 64609 | Incertae sedis sp. "demersal sp1" | k7 | M | 2012 | G. Periat, O. Seehausen | Kivu | Rwanda | Kivu | LVRS Kivu | unknown | unknown | demersal | 106,599,038 | 25.6 | SAMN32926988 | x | x | | |
| 64642 | Incertae sedis sp. "littoral black Mbipia-like" | k11 | M | 2012 | G. Periat, O. Seehausen | Kivu | Rwanda | Kivu | LVRS Kivu | 2 | epilithic algae | rocky | 111,775,638 | 25.9 | SAMN32926993 | x | | | |
| 64173 | Incertae sedis sp. "littoral black Pundamilia-like sp1" | k12 | M | 2012 | G. Periat, O. Seehausen | Kivu | Rwanda | Kivu | LVRS Kivu | unknown | unknown | rocky | 76,463,464 | 18.6 | SAMN32926958 | x | | | |
| 64111 | Incertae sedis sp. "littoral light Incertae sedis-like" | k13 | M | 2012 | G. Periat, O. Seehausen | Kivu | Rwanda | Kivu | LVRS Kivu | unknown | unknown | littoral | 96,942,725 | 23.6 | SAMN32926956 | x | | | |
| 64657 | Incertae sedis sp. "littoral light Paralabidochromis-like sp2" | k5 | M | 2012 | G. Periat, O. Seehausen | Kivu | Rwanda | Kivu | LVRS Kivu | 3 | insectivore | littoral | 83,499,327 | 20.1 | SAMN32926994 | x | | | |
| 64272 | Incertae sedis *occultidens* | k3 | M | 2012 | G. Periat, O. Seehausen | Kivu | Rwanda | Kivu | LVRS Kivu | 4 | paedophage | littoral | 114,287,477 | 27.6 | SAMN32926961 | x | | | |
| 64102 | Incertae sedis *crebridens* | k2 | M | 2012 | G. Periat, O. Seehausen | Kivu | Rwanda | Kivu | LVRS Kivu | 2 | epilithic algae | rocky | 122,710,091 | 30.0 | SAMN32926954 | x | x | | |
| 64421 | Incertae sedis *crebridens* | k2 | M | 2012 | G. Periat, O. Seehausen | Kivu | Rwanda | Kivu | LVRS Kivu | 2 | epilithic algae | rocky | 103,372,757 | 25.4 | SAMN32926972 | | x | | |
| 64106 | Incertae sedis *olivaceus* | k10 | M | 2012 | G. Periat, O. Seehausen | Kivu | Rwanda | Kivu | LVRS Kivu | 2 | epilithic algae | rocky | 106,837,897 | 25.7 | SAMN32926955 | x | x | | |
| 64488 | Incertae sedis *olivaceus* | k10 | M | 2012 | G. Periat, O. Seehausen | Kivu | Rwanda | Kivu | LVRS Kivu | 2 | epilithic algae | rocky | 94,250,057 | 22.8 | SAMN32926975 | | x | | |
| 64517 | Incertae sedis *paucidens* | k6 | M | 2012 | G. Periat, O. Seehausen | Kivu | Rwanda | Kivu | LVRS Kivu | 3 | insectivore | littoral | 97,852,538 | 23.7 | SAMN32926978 | x | | | |
| 64293 | Incertae sedis *vittatus* | k9 | M | 2012 | G. Periat, O. Seehausen | Kivu | Rwanda | Kivu | LVRS Kivu | 4 | piscivore | littoral | 85,041,291 | 20.8 | SAMN32926963 | x | x | | |
| 64394 | Incertae sedis *vittatus* | k9 | M | 2012 | G. Periat, O. Seehausen | Kivu | Rwanda | Kivu | LVRS Kivu | 4 | piscivore | littoral | 78,823,884 | 19.4 | SAMN32926969 | | x | | |
| 64241 | Incertae sedis *graueri* | k4 | M | 2012 | G. Periat, O. Seehausen | Kivu | Rwanda | Kivu | LVRS Kivu | 3 | insectivore | demersal | 83,112,135 | 20.2 | SAMN32926960 | x | | | |
| 64542 | Incertae sedis *kamiranzovu* | k8 | M | 2012 | G. Periat, O. Seehausen | Kivu | Rwanda | Kivu | LVRS Kivu | 2 | phytoplankton | pelagic | 94,408,319 | 22.4 | SAMN32926985 | x | x | | |
| 64594 | Incertae sedis *kamiranzovu* | k8 | M | 2012 | G. Periat, O. Seehausen | Kivu | Rwanda | Kivu | LVRS Kivu | 2 | phytoplankton | pelagic | 79,457,606 | 19.1 | SAMN32926987 | | x | | |
| Sa192 | Incertae sedis sp. "blue" Saka | s2 | M | 2000 | O. Seehausen, L. J. Chapman, C. A. Chapman | Saka | Uganda | Saka | LVRS Saka | 2 | phytoplankton | littoral & demersal | 99,636,378 | 24.6 | SAMN32927094 | | x | x | x |
| Sa193 | Incertae sedis sp. "blue" Saka | s2 | M | 2000 | O. Seehausen, L. J. Chapman, C. A. Chapman | Saka | Uganda | Saka | LVRS Saka | 2 | phytoplankton | littoral & demersal | 105,177,249 | 25.8 | SAMN32927095 | x | x | x | x |
| Sa195 | Incertae sedis sp. "blue" Saka | s2 | M | 2000 | O. Seehausen, L. J. Chapman, C. A. Chapman | Saka | Uganda | Saka | LVRS Saka | 2 | phytoplankton | littoral & demersal | 87,872,627 | 21.3 | SAMN32927096 | | | x | x |
| Sa75 | Incertae sedis sp. "blue" Saka | s2 | M | 2000 | O. Seehausen, L. J. Chapman, C. A. Chapman | Saka | Uganda | Saka | LVRS Saka | 2 | phytoplankton | littoral & demersal | 81,779,109 | 19.8 | SAMN32927099 | | | | x |
| Sa122 | Incertae sedis sp. "yellow-red chest" Saka | s1 | M | 2000 | O. Seehausen, L. J. Chapman, C. A. Chapman | Saka | Uganda | Saka | LVRS Saka | 2 | phytoplankton | littoral & demersal | 88,252,334 | 21.6 | SAMN32927092 | | x | x | x |
| Sa163 | Incertae sedis sp. "yellow-red chest" Saka | s1 | M | 2000 | O. Seehausen, L. J. Chapman, C. A. Chapman | Saka | Uganda | Saka | LVRS Saka | 2 | phytoplankton | littoral & demersal | 91,103,901 | 22.2 | SAMN32927093 | x | x | x | x |
| Sa73 | Incertae sedis sp. "yellow-red chest" Saka | s1 | M | 2000 | O. Seehausen, L. J. Chapman, C. A. Chapman | Saka | Uganda | Saka | LVRS Saka | 2 | phytoplankton | littoral & demersal | 79,056,665 | 19.2 | SAMN32927098 | | | | x |
| Sa85 | Incertae sedis sp. "yellow-red chest" Saka | s1 | M | 2000 | O. Seehausen, L. J. Chapman, C. A. Chapman | Saka | Uganda | Saka | LVRS Saka | 2 | phytoplankton | littoral & demersal | 87,207,808 | 21.4 | SAMN32927100 | | | x | x |
| 103658 | *Astatotilapia* sp. "nubila swamp red" | v11 | M | 2014 | J. van Rijssel, F. Moser, O. Seehausen | Sweya | Tanzania | Victoria | LVRS Victoria | 3 | insectivore | streams | 111,525,085 | 25.7 | SAMN32926781 | | | x | |
| 109429 | *Astatotilapia* sp. "nubila swamp red" | v11 | M | 2014 | J. van Rijssel, F. Moser, O. Seehausen | Sweya | Tanzania | Victoria | LVRS Victoria | 3 | insectivore | streams | 120,270,772 | 28.4 | SAMN32926844 | | x | x | |
| 109432 | *Astatotilapia* sp. "nubila swamp red" | v11 | M | 2014 | J. van Rijssel, F. Moser, O. Seehausen | Sweya | Tanzania | Victoria | LVRS Victoria | 3 | insectivore | streams | 128,515,290 | 30.5 | SAMN32926845 | x | x | x | |
| 104015 | *Enterochromis cinctus* "E" | v15 | M | 2014 | J. van Rijssel, F. Moser, O. Seehausen | MG Transect | Tanzania | Victoria | LVRS Victoria | 2 | detritivore | demersal | 106,463,663 | 25.0 | SAMN15891775 | | x | x | |
| 104016 | *Enterochromis cinctus* "E" | v15 | M | 2014 | J. van Rijssel, F. Moser, O. Seehausen | MG Transect | Tanzania | Victoria | LVRS Victoria | 2 | detritivore | demersal | 110,314,407 | 26.8 | SAMN32926801 | x | x | x | |
| 104020 | *Enterochromis cinctus* "E" | v15 | M | 2014 | J. van Rijssel, F. Moser, O. Seehausen | MG Transect | Tanzania | Victoria | LVRS Victoria | 2 | detritivore | demersal | 108,667,092 | 26.1 | SAMN32926804 | | | x | |
| 22363 | *Gaurochromis hiatus* | - | M | 2014 | J. van Rijssel, F. Moser, O. Seehausen | captive stock | Tanzania | Victoria | LVRS Victoria | 3 | insectivore | demersal | 325,086,660 | 76.2 | SAMN47214503 | | | | |
| 22365 | *Gaurochromis hiatus* | - | F | 2014 | J. van Rijssel, F. Moser, O. Seehausen | captive stock | Tanzania | Victoria | LVRS Victoria | 3 | insectivore | demersal | 182,267,480 | 41.4 | SAMN47214504 | | | | |
| 22415 | *Gaurochromis hiatus* | - | M | 2014 | J. van Rijssel, F. Moser, O. Seehausen | captive stock | Tanzania | Victoria | LVRS Victoria | 3 | insectivore | demersal | 250,875,502 | 59.8 | SAMN47214505 | | | | |
| 22414 | *Gaurochromis hiatus* | - | F | 2014 | J. van Rijssel, F. Moser, O. Seehausen | captive stock | Tanzania | Victoria | LVRS Victoria | 3 | insectivore | demersal | 226,660,986 | 53.4 | SAMN47214506 | | | | |
| 22416 | *Gaurochromis hiatus* | - | U | 2014 | J. van Rijssel, F. Moser, O. Seehausen | captive stock | Tanzania | Victoria | LVRS Victoria | 3 | insectivore | demersal | 155,558,260 | 36.5 | SAMN47214507 | | | | |
| 22417 | *Gaurochromis hiatus* | - | U | 2014 | J. van Rijssel, F. Moser, O. Seehausen | captive stock | Tanzania | Victoria | LVRS Victoria | 3 | insectivore | demersal | 212,509,158 | 48.4 | SAMN47214508 | | | | |
| 22418 | *Gaurochromis hiatus* | - | U | 2014 | J. van Rijssel, F. Moser, O. Seehausen | captive stock | Tanzania | Victoria | LVRS Victoria | 3 | insectivore | demersal | 198,901,975 | 45.4 | SAMN47214509 | | | | |
| 22419 | *Gaurochromis hiatus* | - | U | 2014 | J. van Rijssel, F. Moser, O. Seehausen | captive stock | Tanzania | Victoria | LVRS Victoria | 3 | insectivore | demersal | 224,790,021 | 52.3 | SAMN47214510 | | | | |

| FishecID | Species | sID | Sex | Year | Collectors | Location | Country | Lake | Group | Trophic Level | Trophic Group | Habitat | Number of Reads | Mean Depth | Accession | 2hap | 4hap | 6hap | 8hap |
|---|---|---|---|---|---|---|---|---|---|---|---|---|---|---|---|---|---|---|---|
| 22420 | *Gaurochromis hiatus* | - | U | 2014 | J. van Rijssel, F. Moser, O. Seehausen | captive stock | Tanzania | Victoria | LVRS Victoria | 3 | insectivore | demersal | 191,859,792 | 44.0 | SAMN47214511 | | | | |
| 22421 | *Gaurochromis hiatus* | - | U | 2014 | J. van Rijssel, F. Moser, O. Seehausen | captive stock | Tanzania | Victoria | LVRS Victoria | 3 | insectivore | demersal | 148,242,987 | 34.4 | SAMN47214512 | | | | |
| 22422 | *Gaurochromis hiatus* | - | U | 2014 | J. van Rijssel, F. Moser, O. Seehausen | captive stock | Tanzania | Victoria | LVRS Victoria | 3 | insectivore | demersal | 267,857,618 | 63.2 | SAMN47214513 | | | | |
| 11050 | *Harpagochromis vonlinnei* | v16 | F | 2010 | O. Selz, O. Seehausen | Makobe | Tanzania | Victoria | LVRS Victoria | 4 | piscivore | rocky | 113,338,652 | 26.8 | SAMN15891795 | | x | x | |
| 13135 | *Harpagochromis vonlinnei* | v16 | M | 2010 | O. Selz, O. Seehausen | Makobe | Tanzania | Victoria | LVRS Victoria | 4 | piscivore | rocky | 124,180,472 | 30.4 | SAMN32926921 | x | x | x | |
| 14128 | *Harpagochromis vonlinnei* | v16 | M | 2010 | O. Selz, O. Seehausen | Makobe | Tanzania | Victoria | LVRS Victoria | 4 | piscivore | rocky | 115,208,811 | 26.6 | SAMN32926938 | | | x | |
| 13405 | *Labrochromis* sp. "stone" | v4 | M | 2010 | O. Selz, O. Seehausen | Makobe | Tanzania | Victoria | LVRS Victoria | 3 | snail crusher | rocky | 83,876,777 | 20.5 | SAMN15891826 | | | x | |
| 14259 | *Labrochromis* sp. "stone" | v4 | M | 2010 | O. Selz, O. Seehausen | Makobe | Tanzania | Victoria | LVRS Victoria | 3 | snail crusher | rocky | 138,598,302 | 32.9 | SAMN32926942 | x | x | x | |
| 14262 | *Labrochromis* sp. "stone" | v4 | M | 2010 | O. Selz, O. Seehausen | Makobe | Tanzania | Victoria | LVRS Victoria | 3 | snail crusher | rocky | 111,794,584 | 27.2 | SAMN32926943 | | x | x | |
| 10629 | *Lipochromis* sp. "velvet black cryptodon" | v12 | M | 2010 | O. Selz, O. Seehausen | Makobe | Tanzania | Victoria | LVRS Victoria | 4 | paedophage | rocky | 81,113,331 | 19.1 | SAMN32926828 | | | x | |
| 11045 | *Lipochromis* sp. "velvet black cryptodon" | v12 | M | 2010 | O. Selz, O. Seehausen | Makobe | Tanzania | Victoria | LVRS Victoria | 4 | paedophage | rocky | 106,042,099 | 25.8 | SAMN32926855 | x | x | x | |
| So28 | *Lipochromis* sp. "velvet black cryptodon" | v12 | M | 1995/96 | O. Seehausen | Sozihe | Tanzania | Victoria | LVRS Victoria | 4 | paedophage | rocky | 84,969,781 | 20.5 | SAMN15891866 | | x | x | |
| 11015 | *Lithochromis* sp. "yellow chin" | v14 | M | 2010 | O. Selz, O. Seehausen | Makobe | Tanzania | Victoria | LVRS Victoria | 3 | insectivore | rocky | 99,129,459 | 24.0 | SAMN15891791 | | x | x | |
| 14165 | *Lithochromis* sp. "yellow chin" | v14 | M | 2010 | O. Selz, O. Seehausen | Makobe | Tanzania | Victoria | LVRS Victoria | 3 | insectivore | rocky | 71,403,099 | 16.8 | SAMN32926939 | | | x | |
| 109320 | *Lithochromis* sp. "yellow chin" | v14 | M | 2014 | J. van Rijssel, F. Moser, O. Seehausen | Makobe | Tanzania | Victoria | LVRS Victoria | 3 | insectivore | rocky | 119,600,780 | 28.9 | SAMN32926843 | x | x | x | |
| 79628 | *Macropleurodus bicolor* | v8 | M | 2005 | I. Magalhaes, S. Mwaiko, O. Seehausen | Igombe | Uganda | Victoria | LVRS Victoria | 3 | snail sheller | littoral | 73,582,561 | 17.8 | SAMN15891847 | | | x | |
| Ig140 | *Macropleurodus bicolor* | v8 | F | 2005 | I. Magalhaes, S. Mwaiko, O. Seehausen | Igombe | Uganda | Victoria | LVRS Victoria | 3 | snail sheller | littoral | 113,589,472 | 26.0 | SAMN32927081 | | x | x | |
| Ig158 | *Macropleurodus bicolor* | v8 | M | 2005 | I. Magalhaes, S. Mwaiko, O. Seehausen | Igombe | Uganda | Victoria | LVRS Victoria | 3 | snail sheller | littoral | 115,847,506 | 27.3 | SAMN32927082 | x | x | x | |
| 10561 | *Mbipia mbipi* | v3 | M | 2010 | O. Selz, O. Seehausen | Makobe | Tanzania | Victoria | LVRS Victoria | 2 | epilithic algae grazer | rocky | 104,615,964 | 25.0 | SAMN15891779 | | | x | |
| 11003 | *Mbipia mbipi* | v3 | M | 2010 | O. Selz, O. Seehausen | Makobe | Tanzania | Victoria | LVRS Victoria | 2 | epilithic algae grazer | rocky | 107,706,199 | 26.3 | SAMN32926851 | x | x | x | |
| 11322 | *Mbipia mbipi* | v3 | M | 2010 | O. Selz, O. Seehausen | Makobe | Tanzania | Victoria | LVRS Victoria | 2 | epilithic algae grazer | rocky | 114,636,450 | 26.2 | SAMN32926865 | | x | x | |
| 10888 | *Neochromis greenwoodi* | v7 | M | 2010 | O. Selz, O. Seehausen | Anchor | Tanzania | Victoria | LVRS Victoria | 2 | epilithic algae browser | rocky | 62,328,850 | 14.9 | SAMN32926837 | | | x | |
| 10897 | *Neochromis greenwoodi* | v7 | M | 2010 | O. Selz, O. Seehausen | Anchor | Tanzania | Victoria | LVRS Victoria | 2 | epilithic algae browser | rocky | 76,162,885 | 17.0 | SAMN32926840 | | x | x | |
| 12319 | *Neochromis greenwoodi* | v7 | M | 2010 | O. Selz, O. Seehausen | Python | Tanzania | Victoria | LVRS Victoria | 2 | epilithic algae browser | rocky | 76,127,818 | 17.9 | SAMN15891805 | x | x | x | |
| 10253 | *Neochromis omnicaeruleus* | v10 | M | 2010 | O. Selz, O. Seehausen | Makobe | Tanzania | Victoria | LVRS Victoria | 2 | epilithic algae browser | rocky | 39,934,327 | 9.8 | SAMN32926769 | | | x | |
| 106816 | *Neochromis omnicaeruleus* | v10 | M | 2014 | J. van Rijssel, F. Moser, O. Seehausen | Makobe | Tanzania | Victoria | LVRS Victoria | 2 | epilithic algae browser | rocky | 156,963,507 | 36.8 | SAMN47214514 | x | x | x | |
| N_O | *Neochromis omnicaeruleus* | v10 | M | 2015 | lab bred (M. McGee, O. Seehausen) | Makobe | Tanzania | Victoria | LVRS Victoria | 2 | epilithic algae browser | rocky | 84,652,663 | 20.6 | SAMN15891865 | | x | x | |
| 10610 | *Paralabidochromis chilotes* | v5 | M | 2010 | O. Selz, O. Seehausen | Makobe | Tanzania | Victoria | LVRS Victoria | 3 | insectivore | rocky | 79,736,081 | 19.4 | SAMN15891781 | | x | x | |
| 10618 | *Paralabidochromis chilotes* | v5 | M | 2010 | O. Selz, O. Seehausen | Makobe | Tanzania | Victoria | LVRS Victoria | 3 | insectivore | rocky | 81,301,679 | 19.7 | SAMN32926825 | x | x | x | |
| 10631 | *Paralabidochromis chilotes* | v5 | U | 2010 | O. Selz, O. Seehausen | Makobe | Tanzania | Victoria | LVRS Victoria | 3 | insectivore | rocky | 77,156,731 | 19.0 | SAMN32926829 | | | x | |
| 11591 | *Pundamilia nyererei* | v1 | M | 2010 | O. Selz, O. Seehausen | Makobe | Tanzania | Victoria | LVRS Victoria | 3 | zooplankton | rocky | 75,927,888 | 18.1 | SAMN15891801 | | x | x | |
| 11593 | *Pundamilia nyererei* | v1 | M | 2010 | O. Selz, O. Seehausen | Makobe | Tanzania | Victoria | LVRS Victoria | 3 | zooplankton | rocky | 67,031,671 | 15.8 | SAMN05711163 | | | x | |
| 103528 | *Pundamilia nyererei* | v1 | M | 2005 | lab bred (S. Mwaiko, O. Seehausen) | Makobe | Tanzania | Victoria | LVRS Victoria | 3 | zooplankton | rocky | 123,541,998 | 29.5 | SAMN47214515 | x | x | x | |
| 10554 | *Pundamilia pundamilia* | v2 | M | 2010 | O. Selz, O. Seehausen | Makobe | Tanzania | Victoria | LVRS Victoria | 3 | insectivore | rocky | 99,650,223 | 23.9 | SAMN32926821 | x | x | x | |
| 10560 | *Pundamilia pundamilia* | v2 | M | 2010 | O. Selz, O. Seehausen | Makobe | Tanzania | Victoria | LVRS Victoria | 3 | insectivore | rocky | 73,943,413 | 17.4 | SAMN05711175 | | x | x | |
| 11297 | *Pundamilia pundamilia* | v2 | M | 2010 | O. Selz, O. Seehausen | Makobe | Tanzania | Victoria | LVRS Victoria | 3 | insectivore | rocky | 67,998,763 | 15.6 | SAMN05711178 | | | x | |
| 11546 | *Pundamilia* sp. "nyererei-like" | v6 | M | 2010 | O. Selz, O. Seehausen | Python | Tanzania | Victoria | LVRS Victoria | 3 | zooplankton | rocky | 77,695,351 | 17.8 | SAMN05711235 | | | | x |
| 11719 | *Pundamilia* sp. "nyererei-like" | v6 | M | 2010 | O. Selz, O. Seehausen | Python | Tanzania | Victoria | LVRS Victoria | 3 | zooplankton | rocky | 104,447,964 | 24.5 | SAMN05711237 | | x | x | x |
| 11992 | *Pundamilia* sp. "nyererei-like" | v6 | M | 2010 | O. Selz, O. Seehausen | Python | Tanzania | Victoria | LVRS Victoria | 3 | zooplankton | rocky | 96,650,879 | 22.9 | SAMN05711242 | | | x | x |
| 12170 | *Pundamilia* sp. "nyererei-like" | v6 | M | 2010 | O. Selz, O. Seehausen | Python | Tanzania | Victoria | LVRS Victoria | 3 | zooplankton | rocky | 104,141,970 | 24.6 | SAMN32926891 | x | x | x | x |
| 11545 | *Pundamilia* sp. "pundamilia-like" | v9 | M | 2010 | O. Selz, O. Seehausen | Python | Tanzania | Victoria | LVRS Victoria | 3 | insectivore | rocky | 104,976,312 | 25.7 | SAMN32926877 | x | x | x | x |
| 11725 | *Pundamilia* sp. "pundamilia-like" | v9 | M | 2010 | O. Selz, O. Seehausen | Python | Tanzania | Victoria | LVRS Victoria | 3 | insectivore | rocky | 74,937,130 | 17.4 | SAMN05711261 | | x | x | x |
| 11728 | *Pundamilia* sp. "pundamilia-like" | v9 | M | 2010 | O. Selz, O. Seehausen | Python | Tanzania | Victoria | LVRS Victoria | 3 | insectivore | rocky | 66,706,943 | 15.9 | SAMN05711263 | | | | x |
| 11729 | *Pundamilia* sp. "pundamilia-like" | v9 | M | 2010 | O. Selz, O. Seehausen | Python | Tanzania | Victoria | LVRS Victoria | 3 | insectivore | rocky | 69,445,446 | 16.7 | SAMN05711264 | | | x | x |
| 103754 | *Yssichromis pyrrhocephalus* | v13 | M | 2014 | J. van Rijssel, F. Moser, O. Seehausen | MG Transect | Tanzania | Victoria | LVRS Victoria | 3 | zooplankton | pelagic | 129,653,309 | 31.0 | SAMN32926790 | x | x | x | |
| 109761 | *Yssichromis pyrrhocephalus* | v13 | M | 2014 | J. van Rijssel, F. Moser, O. Seehausen | MG Transect | Tanzania | Victoria | LVRS Victoria | 3 | zooplankton | pelagic | 82,711,452 | 19.6 | SAMN32926791 | | | x | |
| Y_P | *Yssichromis pyrrhocephalus* | v13 | M | 2015 | lab bred (M. McGee, O. Seehausen) | MG Transect | Tanzania | Victoria | LVRS Victoria | 3 | zooplankton | pelagic | 91,545,341 | 21.8 | SAMN15891869 | | x | x | |
| 103218 | *Astatoreochromis alluaudi* | o2 | M | 2014 | J. van Rijssel, F. Moser, O. Seehausen | Makobe | Tanzania | Victoria | Outgroup | 3 | snail crusher | rocky | 111,077,227 | 24.5 | SAMN32926772 | x | x | | |
| 106754 | *Astatoreochromis alluaudi* | o2 | F | 2014 | J. van Rijssel, F. Moser, O. Seehausen | Makobe | Tanzania | Victoria | Outgroup | 3 | snail crusher | rocky | 105,216,414 | 23.7 | SAMN32926832 | | x | | |
| 106985 | *Pseudocrenilabrus multicolor* | o1 | M | 2014 | J. van Rijssel, F. Moser, O. Seehausen | Sweya | Tanzania | Victoria | Outgroup | 3 | insectivore | streams | 93,251,161 | 22.0 | SAMN47214516 | x | | | |
| 64538 | *Thoracochromis gracilior* | n1 | F | 2012 | G. Periat, O. Seehausen | Kivu | Rwanda | Kivu | Upper Nile lineage | 3 | insectivore | littoral | 96,270,424 | 22.6 | SAMN32926983 | | x | | |
| 64539 | *Thoracochromis gracilior* | n1 | F | 2012 | G. Periat, O. Seehausen | Kivu | Rwanda | Kivu | Upper Nile lineage | 3 | insectivore | littoral | 104,301,265 | 24.4 | SAMN32926984 | x | x | | |
| 71001 | *Thoracochromis pharyngalis* | n2 | U | 2019 | J. Fogby, K. Pedersen | Edward | Uganda | Edward | Upper Nile lineage | 3 | insectivore | littoral | 53,362,986 | 13.0 | SAMN32927015 | | x | | |
| 71014 | *Thoracochromis pharyngalis* | n2 | U | 2019 | J. Fogby, K. Pedersen | Edward | Uganda | Edward | Upper Nile lineage | 3 | insectivore | littoral | 49,355,724 | 18.9 | SAMN32927027 | x | x | | |

**Supplementary Table 2 |** *De novo* single base pair mutations in a two-generation *Gaurochromis hiatus* family. The first column shows confirmed grandparent germline mutations (inheritance to the offspring via parents in brackets) and putative parent germline mutations (no brackets). The last column indicates which variant and genotype caller detected a *de novo* single base pair mutation: FB = freebayes, HC = HaplotypeCaller + GVCF pipeline, jHC = joint HaplotypeCaller.

| ID of trio offspring with *de novo* mutation (IDs of F2 inheriting the mutation) | Chromosome:Position | *de novo* Mutation | CpG site | Caller |
|---|---|---|---|---|
| F1: 22414 (F2: 22420, 22421) | chr19:24,638,775 | G>A | | FB+jHC |
| F1: 22414 (F2: 22417, 22418, 22421) | chr18:9,066,532 | G>A | | HC+FB+jHC |
| F1: 22415 (F2: 22417, 22418, 22420) | chr22:11,049,346 | G>A | | FB+jHC |
| F1: 22415 (F2: 22419, 22420, 22421) | scaffold_2854:3,907 | T>C | | FB+jHC |
| F1: 22415 (F2: 22417, 22418, 22422) | scaffold_3519:12 | T>C | | FB+jHC |
| F1: 22415 (F2: 22417, 22420, 22421, 22422) | chr16:23,341,321 | T>C | | HC+FB+jHC |
| F1: 22415 (F2: 22419) | chr5:14,128,976 | C>T | | HC+FB+jHC |
| F1: 22415 (F2: 22416, 22417, 22418, 22420, 22421) | chr8:21,889,646 | G>A | | HC+FB+jHC |
| F1: 22415 (F2: 22419, 22420, 22421, 22422) | chr7:10,901,715 | G>C | | jHC |
| F2: 22416 | chr3:21,331,290 | T>G | | FB+jHC |
| F2: 22416 | chr1:22,182,978 | A>G | | HC+FB+jHC |
| F2: 22417 | chr10:14,100,562 | C>T | X | HC+FB+jHC |
| F2: 22417 | chr9:16,637,941 | T>A | | HC+FB+jHC |
| F2: 22417 | scaffold_245:85,600 | C>T | X | HC+FB+jHC |
| F2: 22417 | scaffold_80:794,765 | A>G | | HC+FB+jHC |
| F2: 22418 | chr16:27,161,429 | G>T | | HC+FB+jHC |
| F2: 22418 | chr8:16,246,805 | T>C | | HC+FB+jHC |
| F2: 22419 | chr12:25,942,673 | A>C | | FB |
| F2: 22419 | chr12:25,942,668 | G>C | | HC+FB |
| F2: 22419 | chr5:5,890,923 | C>T | X | HC+FB+jHC |
| F2: 22420 | chr8:4,876,210 | A>G | | FB |
| F2: 22420 | chr15:11,665,656 | A>G | | HC+FB+jHC |
| F2: 22421 | chr2:19,907,439 | A>G | | HC+FB+jHC |
| F2: 22422 | chr19:26,028,892 | C>T | | HC+FB+jHC |
| F2: 22422 | chr9:9,574,545 | A>G | | HC+FB+jHC |
